# Hill-Robertson interference may bias the inference of fitness effects of new mutations in highly selfing species

**DOI:** 10.1101/2024.02.06.579142

**Authors:** Austin Daigle, Parul Johri

**Affiliations:** Department of Biology, University of North Carolina, Chapel Hill, NC 27599; Department of Genetics, University of North Carolina, Chapel Hill, NC 27599; Curriculum in Bioinformatics and Computational Biology, University of North Carolina, Chapel Hill, NC 27599; Integrative Program for Biological & Genome Sciences, University of North Carolina, Chapel Hill, NC 27599

**Keywords:** distribution of fitness effects, self-fertilization, Hill-Robertson interference, background selection, population structure

## Abstract

The accurate estimation of the distribution of fitness effects (DFE) of new mutations is critical for population genetic inference but remains a challenging task. While various methods have been developed for DFE inference using the site frequency spectrum of putatively neutral and selected sites, their applicability in species with diverse life history traits and complex demographic scenarios is not well understood. Selfing is common among eukaryotic species and can lead to decreased effective recombination rates, increasing the effects of selection at linked sites, including interference between selected alleles. We employ forward simulations to investigate the limitations of current DFE estimation approaches in the presence of selfing and other model violations, such as linkage, departures from semidominance, population structure, and uneven sampling. We find that distortions of the site frequency spectrum due to Hill-Robertson interference in highly selfing populations lead to mis-inference of the deleterious DFE of new mutations. Specifically, when inferring the distribution of selection coefficients, there is an overestimation of nearly neutral and strongly deleterious mutations and an underestimation of mildly deleterious mutations when interference between selected alleles is pervasive. In addition, the presence of cryptic population structure with low rates of migration and uneven sampling across subpopulations leads to the false inference of a deleterious DFE skewed towards effectively neutral/mildly deleterious mutations. Finally, the proportion of adaptive substitutions estimated at high rates of selfing is substantially overestimated. Our observations apply broadly to species and genomic regions with little/no recombination and where interference might be pervasive.

## INTRODUCTION

The distribution of fitness effects (DFE) of new mutations describes the continuum of selective effects of spontaneous mutations in a population and is critical for understanding the effects of natural selection on genetic variation (Eyre-Walker and Keightley 2007). Because of the considerable impact deleterious mutations can have on allele frequency patterns in a population, knowledge of the DFE is often necessary to produce accurate evolutionary models, which are required for unbiased estimates of both the demographic history of a population and the proportion of adaptive nucleotide substitutions in a lineage (Charlesworth 1994; Eyre-Walker and Keightley 2007; Bank et al. 2014; Johri, Aquadro, et al. 2022). While the DFE can be estimated with experimental approaches using random mutagenesis or mutation-accumulation lines (*e.g*., Wloch *et al*. 2001; Estes *et al*. 2004; Sanjuán *et al*. 2004), it can also be inferred from the distribution of allele frequencies from population genetic data (Williamson et al. 2005; Eyre-Walker et al. 2006; Keightley and Eyre-Walker 2007; Galtier 2016). As many organisms cannot easily be cultured in laboratories, are not genetically manipulatable, and/or have large generation times, a population genetic approach remains an important and the only possible approach to estimating the DFE in some species.

Population genetic methods use putatively neutral sites to fit the demographic history or to account for other potential nuisance parameters, while selected sites are used to infer the DFE (by inferring the parameters of a prespecified probability distribution), conditional on the demographic history obtained from neutral sites (reviewed in Johri *et al*. 2022b). While most two-step population genetic approaches that infer the DFE make a series of assumptions, like random mating and the absence of effects of selection at linked sites, they appear to be robust to certain model violations like mutation rate variability (Tataru et al. 2017), background selection (Huber et al. 2017; Kim et al. 2017; Huang et al. 2021), population substructure (Kim et al. 2017), and some complex demographic changes (Kousathanas and Keightley 2013). This is because two-step inference methods are usually employed using interdigitated neutral and selected sites and might therefore correct for model violations that skew the neutral and selected SFS similarly. As an aside, because demographic history is unlikely to affect the SFS of neutral and very strongly deleterious mutations equally (Eyre-Walker et al. 2006), this might lead to slight inaccuracies in inference using methods that do not explicitly model population size changes.

Despite significant progress, not much work has been done on understanding the effects of non-random mating, in particular self-fertilization, on the inference of the DFE using population genetic approaches. Self-fertilization or selfing (*i.e.*, an individual mating with itself) is common across all eukaryotic species - about 35% of all seed plants and ∼50% of all animals exhibit moderate to high rates (50-100%) of selfing (Goodwillie et al. 2005; Jarne and Auld 2006). A transition from random mating to partial selfing has several effects on the population. Firstly, the increase of nonrandom mating in a partially selfing population decreases the effective population size (*N*_*e*_), due to a reduction in the amount of independently sampled gametes (Pollak 1987). There is no change in the probability of fixation of semidominant mutations, but the probability of fixation of recessive or partially recessive beneficial mutations is increased by selfing, and the probability of fixation of recessive or partially recessive deleterious mutations is slightly reduced (Caballero and Hill 1992; Charlesworth 1992). Thirdly, increased homozygosity in the genomes of partially selfing organisms results in a higher proportion of recombination events occurring on nearly identical chromosomes, which decreases the effective rate of recombination (Nordborg 2000). This effect increases the extent of background selection (BGS; Charlesworth *et al*. 1993; Charlesworth and Charlesworth 1998; Charlesworth 2003) and selective sweeps (Hedrick 1980; Hartfield and Bataillon 2020) experienced by the population.

Finally, as the effective rate of recombination is lowered drastically in highly selfing populations, there can be a further decrease in the efficacy of selection due to Hill-Robertson interference (HRI) between selected loci (Hill and Robertson 1966; McVean and Charlesworth 2000; Comeron and Kreitman 2002; Hartfield and Glémin 2014; Hartfield and Glémin 2016).

Hill-Robertson interference (HRI) describes how the efficacy of selection at an allele at one locus can be reduced due to the segregation of selected alleles at a second linked locus (Hill and Robertson 1966; Felsenstein 1974). While BGS and selective sweeps are also a type of HRI, those models assume that selection is strong and that there is no interference between selected sites (reviewed in Charlesworth and Jensen 2021). However, because weakly selected mutations can segregate in the population at higher frequencies, interference between them can result in reduced efficacy of selection and has been termed interference selection (Comeron et al. 1999; Comeron and Kreitman 2002) or weak selection HRI (McVean and Charlesworth 2000; Kaiser and Charlesworth 2009). Moreover, the effects of such interference have been shown to extend to more strongly selected mutations in cases where recombination is greatly reduced (Kaiser and Charlesworth 2009). We therefore refer to the effects of interference selection/weak selection HRI as simply HRI in this study. Since the entire genomes of populations with high selfing rates have greatly reduced effective recombination rates (Nordborg 2000) and deleterious mutations are prevalent across the genome (Bank et al. 2014), HRI is likely to play an important role in determining the dynamics of deleterious mutations in such populations.

Unlike the model of classical BGS, where deleterious alleles are at deterministic mutation-selection equilibrium, selected sites in populations or genomic regions with pervasive HRI depart from the standard equilibrium (Comeron and Kreitman 2002; Kaiser and Charlesworth 2009; O’Fallon et al. 2010; Seger et al. 2010; Nicolaisen and Desai 2013). The site frequency spectra (SFS) of neutral and selected sites affected by HRI are skewed towards rare variants and become more similar to each other as the strength of HRI increases, and these changes cannot be summarized as a simple change in effective population size (Comeron and Kreitman 2002; Kaiser and Charlesworth 2009; O’Fallon et al. 2010; Seger et al. 2010; Nicolaisen and Desai 2013; Good et al. 2014). More recent work assuming equal selection coefficients across deleterious alleles has shown that when HRI is sufficiently common, the nucleotide site diversity can be entirely described by the variation of fitness between individuals in the population. However, populations with different combinations of average selection coefficients, recombination, and mutation rates can have the same nucleotide diversity in this regime (Good et al. 2014), suggesting difficulties in estimating a distribution of selection coefficients along with other population genetic parameters in populations with little/no recombination. While, a recent study suggested that the population-scaled fitness effects of mutations are mis-inferred due to linked effects of selection when the probability of selfing is sufficiently high (Gilbert et al. 2022), the role of HRI (specifically interference) remains unclear. Here, we thoroughly characterize how HRI and other linked effects of selection might bias the inference of both population-scaled as well as direct estimates of selection coefficients in highly selfing species and in regions of low recombination for a variety of DFE shapes. To investigate other potential causes of DFE mis-inference in partially selfing natural populations, we explore the effects of changing the dominance coefficients, coding density, and number of selected sites, as well as the effects of cryptic population structure and sampling on DFE inference. Moreover, we characterize the accuracy of inference of both the DFE of new deleterious mutations as well as the proportion of adaptive substitutions.

## METHODS

### Simulations

All simulations were performed using the forward simulator SLiM (version 4.0.1; Haller and Messer 2023). The genome architecture and population parameters were modeled to mimic the genomes of *C. elegans* populations because they have a high rate of selfing, which has been estimated to be 98%-99.9% using laboratory observations (Hodgkin 1983; Anderson et al. 2010), field work, and population genetic analyses (Barrière and Félix 2005; Barrière and Félix 2007). A chromosome with an intron-exon structure was simulated such that each gene consisted of 6 exons (250 bp in length) and 5 introns (100 bp in length). Genes were interspersed with intergenic regions of length 3000 bp each. These numbers were close to the median of the length of exons, introns, and intergenic regions observed in the *C. elegans* genome with a gene density of ∼1 gene every 5kb (*C. elegans* Sequencing Consortium 1998). The *C. elegans* genome comprises 30% coding, 25% intronic, and 45% intergenic regions (*C. elegans* Sequencing Consortium 1998). In line with these genome-wide characteristics, our simulated genome comprised 30% coding, 10% intronic, and 60% intergenic regions, and a total of 500 genes, resulting in a full chromosomal length of 2,503,000 bp. All introns and intergenic regions were assumed to be neutral, while exons experienced selection.

To take synonymous sites into account, 25% (= *f_neu_*) of all mutations in exons were assumed to be fully neutral (with *s*=0) while the rest were under selection. Fitness was calculated multiplicatively across loci. Epistasis, which could increase or decrease HRI due to changes in the mean equilibrium frequency of deleterious mutations, was not modeled in this study. Of the selected mutations, a fraction *f_pos_* were assumed to be beneficial, where *f_*pos*_* was assumed to be either 0.0, 0.01, or 0.001, and the rest (1 − *f_pos_* − *f_neu_*) were deleterious. The DFE of deleterious mutations was assumed to be a gamma distribution, parametrized by the shape parameter (*β*) and the mean population-scaled selection coefficient 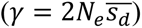, where 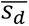 is the mean selective disadvantage of the mutant homozygote relative to wildtype. Three different deleterious DFEs were modeled, representing fitness effects skewed towards mildly, moderately, and strongly deleterious classes respectively (Table 1). The DFE of beneficial mutations was assumed to be exponential with mean 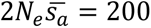, where 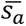 is the mean selective advantage of the mutant homozygote relative to wildtype. All mutations were assumed to be semidominant (where *h* = 0.5) unless otherwise specified.

**Table 1:** Parameters of the simulated DFEs of deleterious mutations in terms of the proportion of mutations in effectively neutral (*f*_0_), weakly deleterious (*f*_1_), moderately deleterious (*f*_2_) and strongly deleterious (*f*_3_) classes. DFEs consisted of predominantly mildly deleterious (DFE1), moderately deleterious (DFE2), or strongly deleterious (DFE3) mutations.

| | $\beta$ | $2Ns_d$ | $f_0$ | $f_1$ | $f_2$ | $f_3$ |
| --- | --- | --- | --- | --- | --- | --- |
| DFE1 | 0.9 | 5 | 0.204 | 0.655 | 0.140 | 0.000 |
| DFE2 | 0.5 | 50 | 0.113 | 0.233 | 0.497 | 0.157 |
| DFE3 | 0.3 | 1000 | 0.098 | 0.097 | 0.192 | 0.614 |

A Wright-Fisher population under equilibrium was simulated with a population size (*N*) of 500,000 diploid individuals (following Gilbert et al. 2022) and with varying probabilities of selfing – 0%, 50%, 80%, 90%, 95%, and 99%. Mutation (*μ*) and recombination (*r*) rates were assumed to be constant at 3.3 × 10^−9^per site/generation (Denver et al. 2004; Konrad et al. 2019; Saxena et al. 2019) and 3.12 × 10^−8^ per site/generation (Rockman and Kruglyak 2009), respectively. Simulations were conducted with a population size of 5000 diploid individuals, with a rescaling factor (*Q*) of 100, and rates of mutation and recombination as well as selection coefficients were correspondingly adjusted so that the population-scaled parameters *Nμ*, *Nr*, and *Ns* were equivalent to a population of 500,000. Simulations were run for a total of 14*N* generations, where 10*N* generations corresponded to the burn-in period and 4*N* generations were used to calculate the number of substitutions and where *N* represents the scaled population size. Five replicates of each evolutionary scenario were simulated and the DFE was estimated for each replicate separately.

#### Single-locus simulations with selfing

To investigate the effects of selfing on DFE inference without linkage, selected, semidominant mutations were simulated at single sites with probabilities of selfing of 0%, 50%, and 99%. As allele frequencies of neutral mutations are unaffected by linkage in the absence of selected mutations, 187,500 linked neutral sites were simulated, equivalent to the number of neutral sites present in 500 *C. elegans* genes. 50,000 selected sites were simulated for each scenario (compared to 562,500 selected sites in previous simulations), then paired with neutral mutations simulated with the appropriate selfing rate. The simulations were rescaled by a factor of 100 and run for 14*N* generations with 10*N* burn-in generations.

#### Simulations under low recombination rates with no selfing

To clarify if the lower effective recombination rates in a partially selfing population were responsible for DFE mis-inference, we conducted simulations with low recombination rates and 0% selfing. As before, these simulations were rescaled by a factor of 100. The *C. elegans* rescaled recombination rate of 3.12 × 10^−6^ was multiplied by 0.5, 0.1, 0.05, 0.01, 0.005, or 0.001. Beneficial mutations were not included in these simulations.

#### Simulations with varying dominance coefficients

All selected mutations shared a single dominance coefficient of 0.1, 0.25, or 0.75. Populations with selfing probabilities of 0%, 50%, and 99% were simulated and compared to the original simulations (where *h* = 0.5).

#### Simulations under varying coding density and chromosome size

Simulations were performed with varying coding densities by varying the intergenic lengths - 1500 bp and 500 bp. The total number of genes was not increased in these simulations. Thus, in the simulations with 1500 bp intergenic regions, the simulated genome was comprised of ∼43% coding, ∼14% intronic, and ∼43% intergenic regions, with a total length of 1,752,100 bp. In the simulations with 500 bp intergenic regions, the genome was comprised of ∼55% coding, ∼18% intronic, and ∼27% intergenic regions, with a total length of 1,376,650 bp. These simulations were run with probabilities of selfing equal to 0%, 50%, and 99%.

To study the effects of a more realistically sized chromosome on DFE inference in selfers, the total number of genes was increased from 500 to 3000, corresponding to approximately the number of genes observed on the first chromosome of the *C. elegans* genome, with a total simulated length of 15,018,000 bp, close to the true size of *C. elegans* chromosome I (15,072,434 bp) (https://www.ncbi.nlm.nih.gov/datasets/genome/GCF_000002985.6/; last accessed in Dec. 2023).

#### Simulations with varying rescaling factors

To investigate the impact of rescaling of population sizes, simulations were conducted with the rescaling factor *Q* = 50, 20, and 10, where the population-scaled parameters *Nμ*, *Nr*, and *Ns* were equivalent to a population of 500,000.

Simulations were run for a total of 14*N* generations, where 10*N* generations corresponded to the burn-in period and 4*N* generations were used to calculate the number of substitutions and where *N* represents the scaled population size. Due to the excessive run time of unscaled simulations, simulations were only run with selfing rates of 95% and 99%. For *Q* = 10, only one replicate of DFE3 at 95% selfing and three replicates each for DFE2 and DFE3 at 99% selfing reached completion.

### Quantifying the effect of BGS in simulated data

To determine the theoretically expected neutral nucleotide diversity (*π*) in simulations with partial selfing, we calculated the expected effective population size after correcting for selfing (*N*_*self*_) as *N*_*self*_ = *N*/(1 + *F*), where *F* = *P*_*self*_/(2 − *P*_*self*_), and *P*_*self*_ represents the proportion of selfing individuals in a population per generation (Pollak 1987). Without the linked effects of selection, the neutral *π* is expected to be approximated by *π* ∼ *θ* = 4*N*_*self*_μ. In simulations with deleterious mutations, the extent of BGS was quantified as the observed nucleotide diversity (*π*_*obs*_) at neutral sites relative to that expected under strict neutrality, *i.e*.,

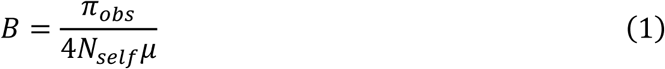

### Simulations with population structure

To study the effects of population structure on DFE inference in selfers, a finite island model (Maruyama 1970) with five demes and an equal probability of exchange of migrants between all demes was considered. Note that in this model the migration rate is independent of the selfing rate, which applies well to animals but is not necessarily the case in plant species because selfing can prevent pollen dispersal (reviewed in Sicard and Lenhard 2011). Scenarios with different migration rates were simulated such that the effective metapopulation size remained constant and equivalent to the size of the panmictic population simulated above. Genome structure was identical to that in the previous simulations mimicking *C. elegans* genome architecture. In a metapopulation with *d* demes and a deme effective population size *N*_*deme*_, the mean coalescence time for a pair of alleles sampled from the same deme (*T_S_*) is given by the following equation (Nei 1973; Slatkin and Voelm 1991; Charlesworth and Charlesworth 2010):

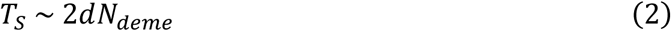

The mean coalescence time for a pair of alleles sampled from different demes (*T_B_*) is given by (Nei 1973; Slatkin and Voelm 1991; Charlesworth and Charlesworth 2010):

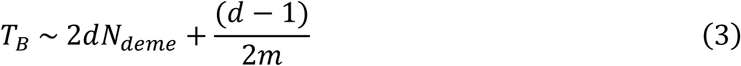

where the migration rate *m* is the probability that an allele in one deme is derived by migration from any other deme in a generation (which means that the migration rate between any two demes is equal to 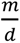). Combining these two equations gives the mean coalescence time (*T_T_*) for a pair of alleles drawn from the entire metapopulation (Nei 1973; Slatkin and Voelm 1991; Charlesworth and Charlesworth 2010):

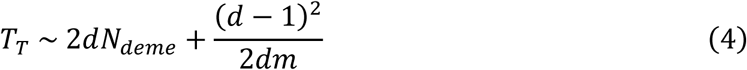

To keep the effective size of the metapopulation (0.5*T*_*T*_) equivalent to the rescaled size of 5,000 individuals used in previous simulations, *N*_*deme*_ and *m* were varied such that *T*_*T*_ remained equal to 10,000. To allow a wide range of migration rates, *N*_*deme*_*m* was varied from 10, 2, 1, 0.5, and 0.1 in populations with 0%, 50%, and 99% selfing, with separate simulations for DFE1, DFE2, and DFE3. The exact parameters used in each simulation can be found in Supplementary Table 7. Parameters were chosen so that the predicted *F*_*ST*_ of the entire metapopulation 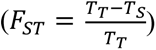 would cover a range of values (from 0.02 to 0.76) matching previously estimated values for *C. elegans* populations in the literature (from 0.15 to 0.43; Cutter 2006). Neutral simulations with identical genome structure and migration parameters were used to verify the joint effects of population structure and selfing on neutral nucleotide diversity and *F*_*ST*_, and were found to yield expected results (Supplementary Table 8). The expected neutral nucleotide diversity for partially selfing populations was calculated assuming that the effective size of each deme will be equivalent to 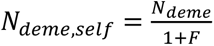, which was used to calculate the appropriate *T_T_* and *T_S_* values for each evolutionary scenario. In SLiM, *m* represents the proportion of individuals in a deme that are migrants each generation. Therefore, *m* still represents the probability that an allele is derived from migration each generation, and *N*_*deme*_*m* represents the number of diploid migrants per deme per generation.

Twenty genomes were sampled from each deme (totaling 100 genomes from the metapopulation) for DFE inference. To understand the effects of uneven sampling on DFE inference, two alternative sampling schemes were employed. In the first, 60 genomes were sampled from one deme, while 10 genomes were drawn from the remaining four demes. In the second, 35 genomes were drawn from two demes, while 10 genomes were drawn from the remaining three demes.

### Calculating the SFS and the number of adaptive substitutions from simulations

The full output provided by SLiM was used to generate the site frequency spectra of neutral and selected mutations in exonic regions individually. The fixed class was obtained as the number of substitutions that occurred after the burn-in period (= 10*N* generations). The zero class was calculated by subtracting the number of polymorphic and fixed sites from the total number of sites for both neutral (= 0.25 × number of exonic sites) and selected (= 0.75 × number of exonic sites) mutations respectively. Scripts to calculate the SFS are available at https://github.com/JohriLab/DFESelfing/tree/main/calculateSFS.

The true fraction of beneficial substitutions (*α*) in simulations was obtained as follows:

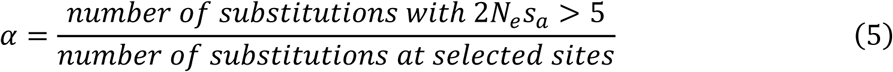

Only substitutions that occurred after the burn-in with 2*N*_*e*_*s*_*a*_ > 5 were included when calculating the true value of *α* in order to match the assumptions of GRAPES.

### DFE-alpha

*Est_dfe* (Keightley and Eyre-Walker 2007) was employed to infer the DFE of deleterious mutations. We fitted a two-epoch model, with starting values of *t*2 (time of change), *β*, and 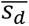 equal to 50, 0.5, and -0.01 respectively. *N*2 (current size) and *t*2 were set to be variable. In the absence of positive selection, the unfolded SFS was used as input, and the fold setting was set to 0. In the presence of beneficial mutations, the folded SFS was used in addition to the unfolded SFS because DFE-alpha has been shown to be sensitive to high frequency beneficial alleles in the SFS (Tataru et al. 2017). DFE-alpha provides the most likely estimate of the expected value of 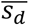, the shape parameter (*β*) of the gamma distribution, and a weighted effective population size (*N*_*w*_). These parameters were used to obtain the proportion of mutations (*f*_0_, *f*_1_, *f*_2_, *f*_3_) in the four classes- effectively neutral (0 ≤ 2*N*_*w*_*s*_*d*_ < 1), weakly deleterious (1 ≤ 2*N*_*w*_*s*_*d*_ < 10), moderately deleterious (10 ≤ 2*N*_*w*_*s*_*d*_ < 100), and strongly deleterious (100 ≤ 2*N*_*w*_*s*_*d*_), respectively.

## GRAPES

The program GRAPES (Eyre-Walker and Keightley 2009; Galtier 2016), Version 1.1.1 (https://github.com/BioPP/grapes/releases/tag/v1.1.1) was used to infer the full DFE using simulations with and without positive selection. GRAPES provides the maximum likelihood estimate of the shape parameter (*β*) and the expected population-scaled mean strength of selection in terms of 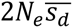, in order to characterize the gamma distribution representing the deleterious DFE. GRAPES also infers the parameters of the beneficial DFE, including the proportion of substitutions that are beneficial mutations (*α*) and the mean population-scaled strength of selection favoring new advantageous mutations 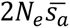, assuming that advantageous selection coefficients are exponentially distributed. The unfolded SFS was provided as input for both scenarios (with and without positive selection). In simulations without positive selection, the default parameters for the GammaZero model were used (*i.e*., parameters of the beneficial DFE were not inferred). In the presence of positive selection, the estimated beneficial DFE was assumed to follow an exponential distribution (*i.e*., the GammaExpo model was used). As GRAPES can use either the folded or the unfolded SFS to provide estimates of the beneficial DFE, we performed inference using the unfolded and folded SFS separately to compare the results of both estimation methods. These parameters were accordingly used to obtain the values *f*_0_, *f*_1_, *f*_2_, *f*_3_, representing the classes of the deleterious DFE as described above.

### Comparison of the expected and estimated DFE when mutations are semidominant

In a partially selfing population, diffusion theory provides the following mean (*M*_Δ*q*_) and variance (*V*_Δ*q*_) of the change in allele frequency (*q*) assuming that the change (Δ*q*) in allele frequency is small (Caballero et al. 1991):

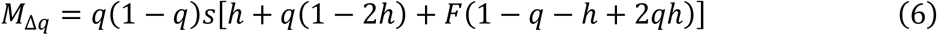

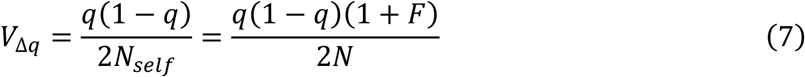

For completely additive mutations (*h* = 0.5), the mean reduces to:

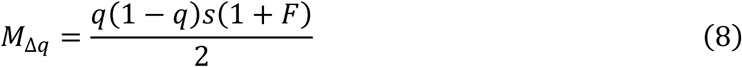

In a panmictic population (of size *N*) with random mating, the mean and variance of change in allele frequency (assuming semidominance) is given by:

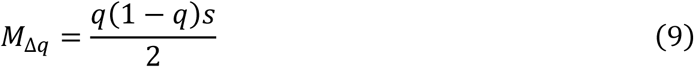

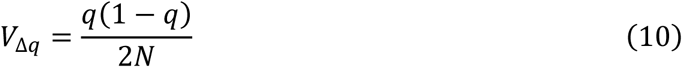

Thus, by comparing the above set of equations, a selfing population of size *N* experiences more genetic drift than a randomly mating population of the same size (Caballero and Hill 1992).

However, the mean change in allele frequency in selfing populations is also larger, by the same factor, (1 + *F*), and thus the fixation probability of mutations remains the same in a selfing *vs*. a randomly mating population (Charlesworth and Charlesworth 1987; Caballero and Hill 1992; Charlesworth 1992). Thus, for the case of additive mutations and not taking selection at linked sites into account, the expected DFE would be determined by 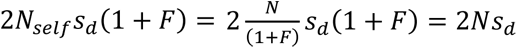. To account for the relative decrease in population size due to BGS or selective sweeps, the simulated DFE was adjusted and presented as 2*NBs*_*d*_, where *B* was calculated using equation 1.

In the case of the estimated DFE, because DFE-alpha only infers a relative increase or decrease in population size 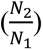, these estimates of demographic parameters should not be affected by selfing rates (where the factor 1 + *F* will cancel out). We thus present our inferred DFE as 2*N*_*w*_*s*_*d*_.

### Comparison of the expected and estimated DFE when mutations are not semidominant

When mutations were not additive, our expected DFE was parameterized by 2*N*_*self*_*s*_*d*_*h*_*self*_, where

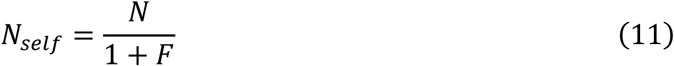

and

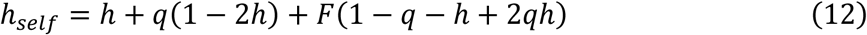

Assuming that *q* << 1, this equation simplifies to:

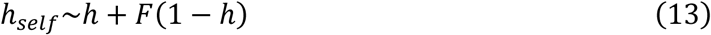

Note that this approximation is less accurate when selection is weak (2*Ns*_*d*_∼1) and thus might not be as accurate for the DFE comprising predominantly weakly deleterious mutations. The estimated DFE (using DFE-alpha and GRAPES) was parameterized by 2*N*_*w*_*s*_*d*_*h* where *h* = 0.5 is assumed by both the programs.

### Calculation of summary statistics from simulated data

In order to quantify the effects of HRI, the skew in the SFS towards rare alleles was measured using the summary statistic 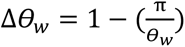, which has been demonstrated to be less sensitive than Tajima’s *D* to the number of SNPs used in the calculation (Becher et al. 2020). Here, *π* is the nucleotide diversity (Nei and Tajima 1981) and *θw* denotes Watterson’s *θ* (Watterson 1975). In addition, the number of singletons (*sing*) and haplotype diversity (hapdiv; Ferrer-Admetlla et al. 2014) were calculated. To summarize linkage disequilibrium, *r*^2^(Hill and Robertson 1968),

*D*, and *D*′ (Lewontin 1964) were calculated using minor alleles for pairs of polymorphisms at frequencies between 1% and 5%. Only low-frequency SNPs were used for *D*′ calculations to correct for the allele frequency dependence of linkage disequilibrium statistics (Lewontin 1988). All summary statistics were calculated separately for neutral and selected polymorphisms in exonic regions in nonoverlapping 5000 bp windows using *pylibseq* v. 0.2.3 (Thornton 2003).

### Estimation of the extent of background selection in highly selfing species

To estimate the levels of background selection (*B*) in highly selfing species, we collated previous estimates for the mutation (*μ*) and recombination rate (*r*) per site/generation, the fraction of the genome that is coding (*f*_*coding*_), and selfing rates for highly selfing species that have publicly available genome assemblies. In species where the mutation rate had not been measured, we used closely-related species to approximate the rate (see Supplementary Table 9 for details). Estimates of *B* were obtained for each species using 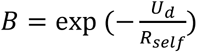 (Nordborg *et. al* 1996), where *U_d_* (the genome-wide mutation rate for a diploid individual) was estimated as *U*_*d*_ = 2 × *L* × *f*_*coding*_ ∗ *μ* × 0.7, where *L* is the total length of the genome, 0.7 represents the fraction of nonsynonymous sites, and *R*_*self*_ = *R* × *F*, where *R* was estimated as *R* = *r* × *L*.

## RESULTS

### Inference of the DFE of new deleterious mutations in self-fertilizing populations

We simulated a single *C. elegans*-like population with varying rates of selfing – 0, 50, 80, 90, 95, and 99% and three different DFEs (following a gamma distribution; parameters specified in Table 1) skewed towards mildly (DFE1), moderately (DFE2), and strongly deleterious (DFE3) classes respectively. Here we assumed that all new mutations were semidominant. We used DFE-alpha and GRAPES to infer the parameters of the deleterious DFE, specified to be gamma-distributed. As expected, when there is no selfing, inference was accurate (Figure 1, Supplementary Figure 1, Supplementary Table 1, Supplementary Table 2). As the selfing rate increased (≥ 90%), the estimated DFE (*i*.*e*., distribution of population-scaled selection coefficients) was skewed strongly towards the class of mildly deleterious mutations (Figure 1), when compared to the simulated DFE, with a particularly substantial skew in the most highly selfing population, consistent with Gilbert *et al*. (2022). While the inferred 2*Ns̅*_*d*_was highly underestimated (Supplementary Figure 1)., the *β* parameter of the gamma distribution was also mis-inferred as the selfing rate increased, with the inferred *β* becoming 2-3× smaller than the simulated value at 99% selfing (Supplementary Table 2). The mean population-scaled selection coefficients remained slightly biased even when selfing rates were as low as 80% (*e.g*. for DFE2; Figure 1, Supplementary Table 2).

**Figure 1:**
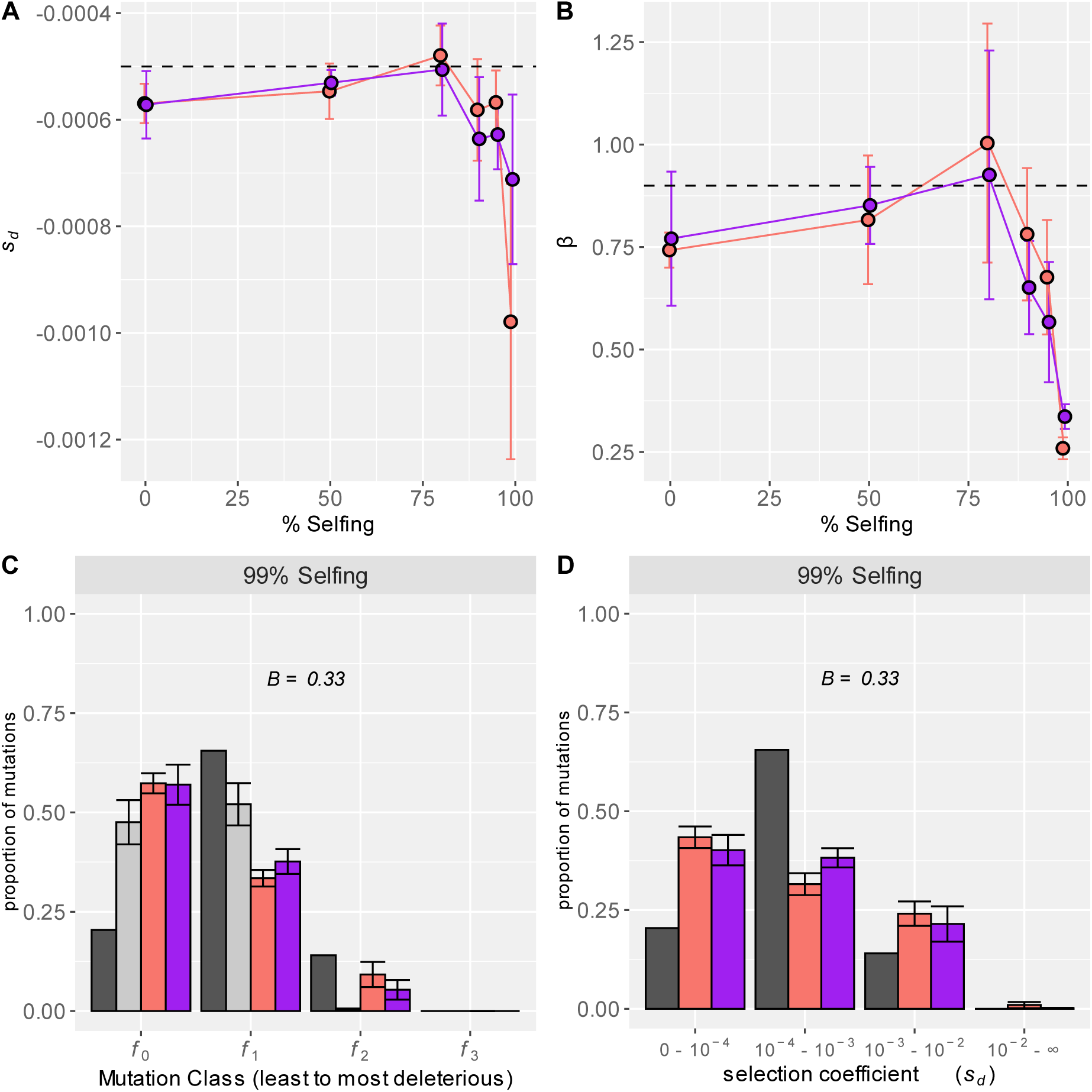
Effects of selfing on the inference of the DFE of new deleterious mutations using DFE-alpha (red bars) and GRAPES (purple bars). (**A**) The mean selection coefficients 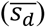 and (**B**) shape parameters (*β*) of the inferred gamma distributions for varying selfing rates are shown. (C) A comparison of the simulated and inferred DFE at 99% selfing rate is shown in terms of the proportion of mutations in the effectively neutral (*f*_0_), weakly deleterious (*f*_1_), moderately deleterious (*f*_2_) and strongly deleterious (*f*_3_) classes of mutations. The simulated (black bars) and adjusted (grey bars) DFE represents the distribution of 2*Ns*_*d*_ and 2*NBs*_*d*_ respectively. (**D)** The simulated and inferred distribution of *s*_*d*_, which was created by dividing the mean of the gamma distribution (2*N*_*e*_*s̅*_*d*_) by 2*BN*. The nucleotide site diversity with background selection (*B*) relative to its expectation under strict neutrality is shown in **C** and **D**. In all panels, the error bars denote the standard deviation of proportions estimated from 5 independent replicates.

Because BGS will lead to a decrease in the effective population size, we hypothesized that the true population-scaled selection coefficients in our simulated populations are likely to be smaller than those simulated (*i*.*e*., 2*Ns*_*d*_). To account for the reduction in the efficacy of purifying selection due to BGS, we corrected the simulated DFE with the relative rate of coalescence (*B*) with and without BGS observed in the simulated population. Note that this adjusted DFE accounts for the relative decrease in population size due to linked effects of selection only, not due to selfing (see details in Methods). While the inferred DFE matches the corrected expected DFE substantially well for moderate levels of selfing (Figure 1; Supplementary Figure 1), the DFE accounting for BGS is somewhat inaccurate at high selfing levels (Figure 1C; Supplementary Figure 1).

We next estimated the distribution of selection coefficients by accounting for the decrease in the effective population size due to BGS. Consistent with the observation above, the distribution of selection coefficients is accurately inferred when selfing rates are below 0.8 (Supplementary Figure 2) but remains inaccurate at extremely high selfing rates (0.95-0.99; Figure 1D; Supplementary Figure 2). That is, the inferred DFE (in terms of the selection coefficients) tends to have both more strongly deleterious mutations and nearly neutral mutations and results in a more platykurtic than the simulated distribution (Figure 1D; Supplementary Figure 2). This suggests that a simple rescaling of the effective population size can correct for the misinference when selfing rates are low/moderate (when the population is likely experiencing classical BGS) but not when rates are high (when the population is likely in the interference selection regime). This effect is especially substantial when a higher proportion of the DFE is composed of weakly and moderately deleterious mutations (*e.g.* DFE1 and DFE2), which are more likely to experience HRI(Figure 1D; Supplementary Figure 2).

The simulated DFEs were accurately estimated from single-locus simulations with no linkage even in highly selfing populations (Supplementary Figure 3), despite a significant reduction in the total number of segregating sites at high selfing (Supplementary Table 3). Concordantly, the observed SFSs at selected and neutral sites were highly similar across different selfing levels, confirming that selfing does not drastically influence the proportions of allele frequencies in a population without linkage (Supplementary Figure 4), assuming that deleterious mutations are additive. Importantly, as high rates of selfing increase the levels of homozygosity, the site frequency spectrum (SFS) obtained from a random sample of diploid *individuals* deviated starkly from one obtained by a random sampling of *genomes* (an example shown in Supplementary Figure 5A). The inferred DFEs were very similar when we sampled individuals *vs*. genomes (Supplementary Figure 5B) and thus all results that follow were obtained by sampling genomes.

### The effects of linkage on DFE inference in regions of low recombination

HRI is more likely in genomic regions of low recombination and may similarly bias DFE inference. With decreasing rates of recombination, the extent of mis-inference of the DFE increased (Figure 2A, B). As before, while the DFE accounting for BGS explained the estimated DFEs quite well for higher recombination rates, at very low recombination rates, it did not match the inferred DFE (Figure 2C). The inferred distribution of selection coefficients tended to overestimate the proportion of strongly deleterious and nearly neutral mutations and underestimated the proportion of mildly deleterious mutations (Figure 2D; Supplementary Figure 6). As with high selfing, the estimated distribution of selection coefficients at low recombination was observed to be more platykurtic than that simulated, and mis-inference was much stronger at the lowest recombination rates, even for the most strongly deleterious DFE (Supplementary Figure 7). This again reflects the fact that a simple rescaling effect of BGS (*i.e*., the decrease in the rate of coalescence) is unlikely to explain mis-inference observed in regions of very low recombination due to HRI. Note that the relative decrease in nucleotide diversity 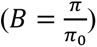 due to background selection, measured in simulations of regions with low recombination (*B*∼0.05 − 0.92) captured the range of *B* values measured in the simulations with selfing and linkage (*B*∼0.08 − 0.95).

**Figure 2:**
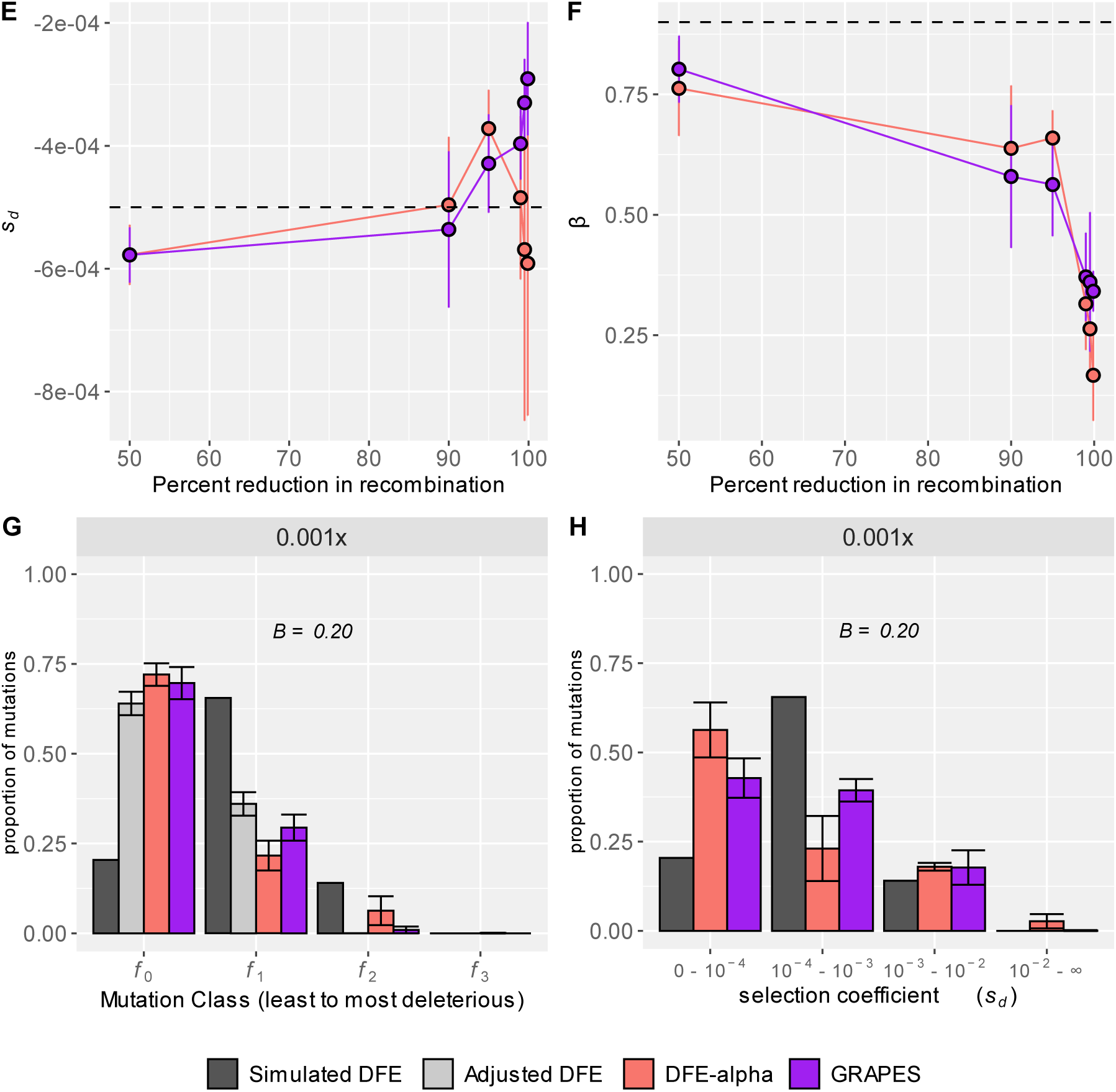
Effects of reduced recombination rates on the inference of the DFE of new deleterious mutations using DFE-alpha (red bars) and GRAPES (purple bars). (**A**) The mean selection coefficients 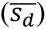 and (**B**) shape parameters (*β*) of the inferred gamma distributions are plotted for varying recombination rates, reduced from an initial rate of *r* = 3.12 × 10^−8^ per site/generation. (**C**) A comparison of the simulated and inferred DFE at a recombination rate reduced by 99.9% is shown in terms of the proportion of mutations in effectively neutral (*f*_0_), weakly deleterious (*f*_1_), moderately deleterious (*f*_2_) and strongly deleterious (*f*_3_) classes of mutations. The simulated (black bars) and adjusted (grey bars) DFE represents the distribution of 2*Ns*_*d*_ and 2*NBs*_*d*_ respectively. (**D**) The simulated and inferred distribution of *s*_*d*_, which was created by dividing the mean of the gamma distribution (2*N*_*e*_*s*_*d*_) by 2*BN*. The nucleotide site diversity with background selection (*B*) relative to its expectation under strict neutrality is shown in **C** and **D**. In all panels, the error bars denote the standard deviation of proportions estimated from 5 independent replicates.

### Parameters that modulate the extent of HRI

Population-scaled parameters such as *NU*, *NR*, and *Ns* all influence the extent of interference between selected alleles (Good et al. 2014). When the ratio of mutation to recombination is sufficiently high and the strength of purifying selection is low, interference between selected mutations becomes sufficiently common to cause them to diverge from the predictions of single site models. To investigate whether these parameters similarly determine the extent of DFE misinference, we explored how 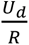 and 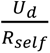 relates to the degree of misinference of *s* and *β* (Figure 3A) in the simulations with low recombination and selfing respectively, where *U*_*d*_ is the genome-wide deleterious mutation rate, *R* is the map length of the simulated chromosome (assuming that recombination scales linearly with distance), and *R*_*self*_ = *R* × (1 − *F*), where *F* = *P*_*self*_/(2 − *P*_*self*_) (Nordborg 2000). We found that error is relatively small at low 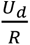 ratios, but when 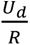 is greater than ∼5, error began to rapidly increase, with the largest error present in the DFE with predominantly weakly deleterious mutations. Interestingly, we found that the extent of BGS, as measured by *B*, was by itself a good predictor of the extent of this error, finding that as *B* decreased, the error in both *s̅*_*d*_ and *β* increased, and that the relationship between *B* and error was very similar in the low recombination and selfing simulations (Figure 3B). Although this might help determine when inference could be biased, it is difficult to obtain true values of *B* from empirical data especially if the genome-wide rate of recombination is low.

**Figure 3:**
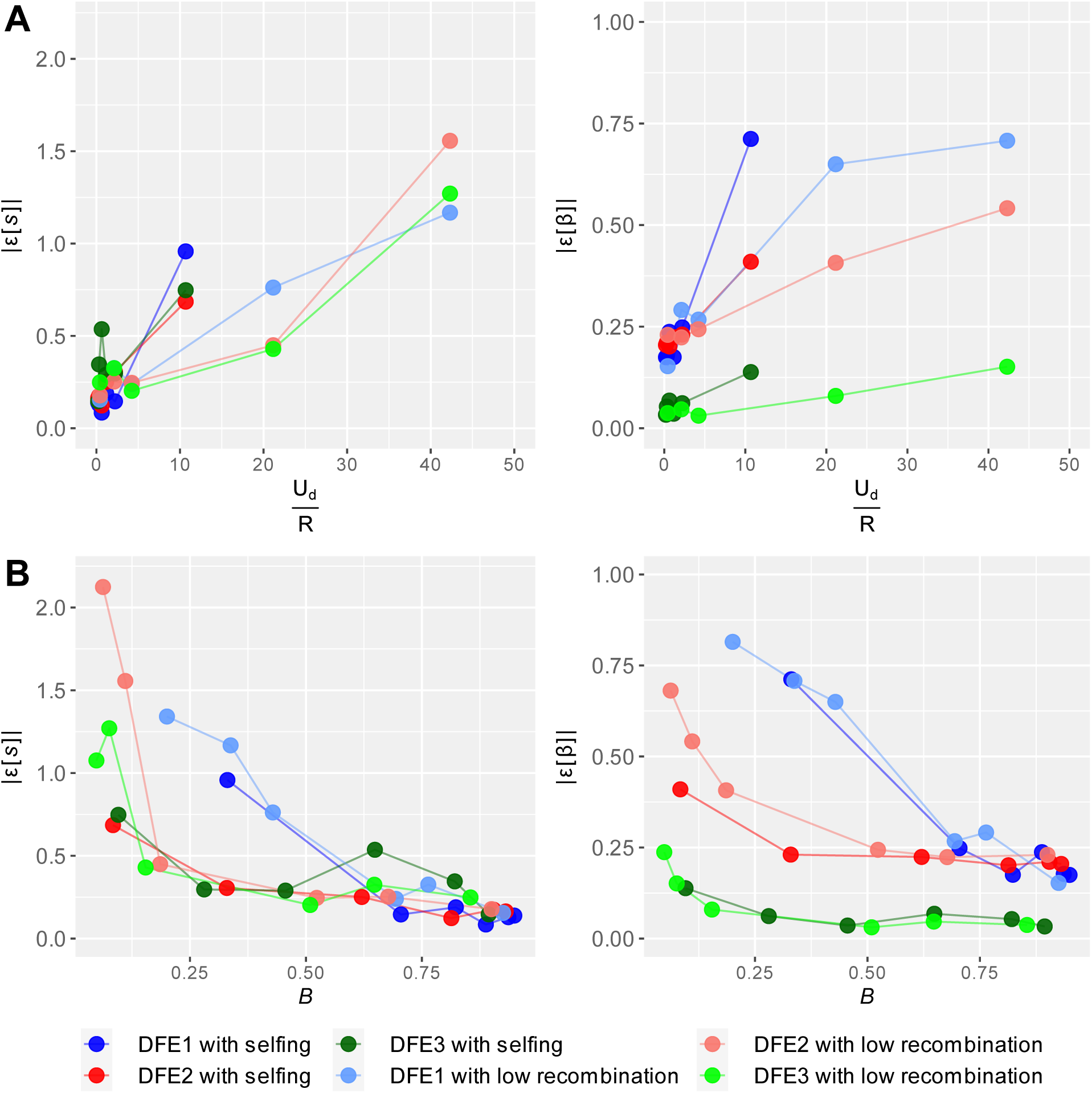
Correlation of error 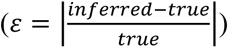 of parameters in DFE inference by DFE-alpha with (**A**) the ratio of number of deleterious mutations per diploid genome per generation (*U*_*d*_) and number of effective recombination events per individual per generation (*R*) and (**B**) the neutral nucleotide site diversity (B) with background selection relative to that expected under neutrality (*B*). Error in the inference of the parameters of the DFE and levels of background selection were averaged over five independent replicates.

### The quantification of HRI using summary statistics

Much theoretical and computational work on HRI has suggested that the decrease in the efficacy of selection results in a decrease in the nucleotide diversity at selected and linked neutral sites as well as an increase in divergence at selected sites due to the increased fixation of deleterious mutations (Comeron et al. 1999; McVean and Charlesworth 2000). Moreover, the site frequency spectrum for selected and neutral sites is expected to be skewed towards low-frequency variants as the strength of BGS and HRI increases (mimicking the genealogy of an expanding population; Zeng and Charlesworth 2011; Walczak *et al*. 2012; Nicolaisen and Desai 2013; Good *et al*. 2014), and the nucleotide diversity and site frequency spectrum of selected mutations are expected to become more similar to those of neutral mutations (Comeron and Kreitman 2002; Kaiser and Charlesworth 2009; O’Fallon et al. 2010; Seger et al. 2010; Nicolaisen and Desai 2013). Because interference results in a gene genealogy with longer external branches than expected under a standard neutral coalescent (Walczak *et al*. 2012; Nicolaisen and Desai 2013), and consistent with a skew towards low-frequency alleles, new mutations are more likely to occur on separate branches of the genealogy, generating negative LD. Thus signed measures of linkage disequilibrium between selected mutations and nearby neutral sequences become more negative as HRI becomes stronger (Hill and Robertson 1966; McVean and Charlesworth 2000; Comeron and Kreitman 2002; Garcia and Lohmueller 2021). In addition, in populations with pervasive HRI, the SFS has been observed to show non-monotonic behavior at high frequencies, resulting in a U-shape, which deviates from the predictions of the classical BGS theory (Seger et al. 2010; Neher and Hallatschek 2013; Good et al. 2014; Zeng and Corcoran 2015; Cvijović et al. 2018).

In our simulations, the SFSs of neutral and selected sites shifted towards lower frequency mutations and became more similar at 90% selfing and above (Figure 4A, Supplementary Figures 8A & 9A), which is consistent with the extent of DFE mis-inference and with previously simulated results in highly selfing populations (Charlesworth *et al*. 1993). In addition, the SFS at high frequency alleles showed non-monotonic (U-shaped) behavior (Figure 4B, Supplementary Figures 8B & 9B). Note that the SFS of neutral sites at 0% selfing was very similar to the unlinked neutral SFS (Supplementary Figure 4), consistent with high *B* values. Trends in Δ*θ*_*w*_, a measure of the skewness of the SFS (with positive values indicating a relative increase in rare alleles), were concordant with the patterns in the SFSs, with Δ*θ*_*w*_ values for neutral and selected sites sharply increasing and converging towards each other at 90% selfing and above (Figure 5). Values of Δ*θ*_*w*_ at neutral sites nearly converge with values at selected sites at 99% selfing, reflecting the reduced efficacy of selection and almost complete linkage. Similar patterns were observed in the simulations with low recombination, with values of Δ*θ*_*w*_ increasing sharply and converging at low recombination rates (Supplementary Figure 10).

**Figure 4:**
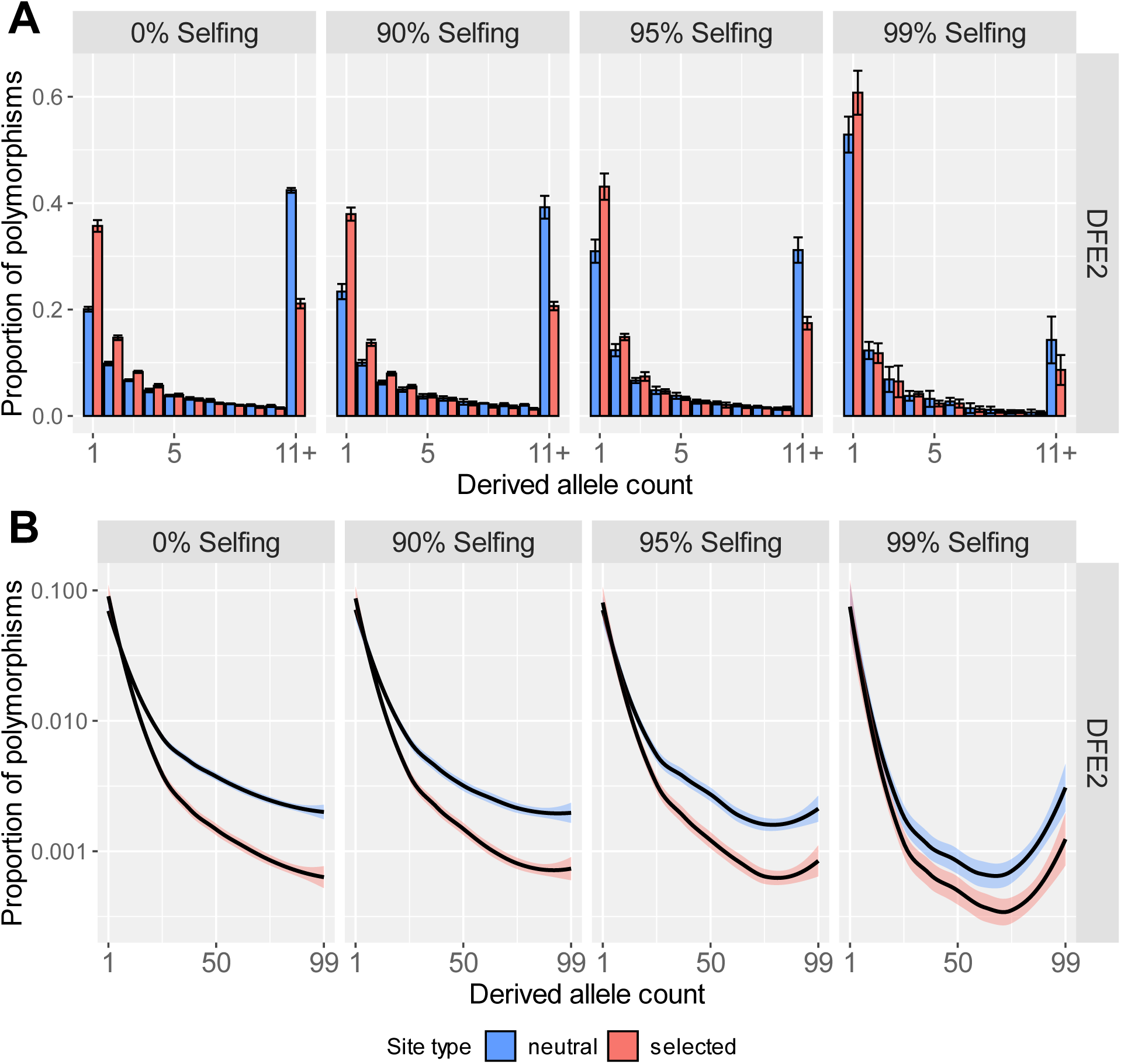
Site frequency spectra of neutral and selected alleles from 100 genomes sampled from simulations where all selected mutations were deleterious, for the moderately deleterious DFE (DFE2). For **(A)** the y-axis represents the proportion of segregating polymorphisms that fall into the given derived allele count. The last class (11+) refers to the derived allele counts 11-99. The error bars denote the standard deviation of proportions estimated from 5 independent replicates. For **(B)** the y-axis is plotted on a log scale, with all derived allele counts displayed. Lines were smoothed with LOESS.

**Figure 5:**
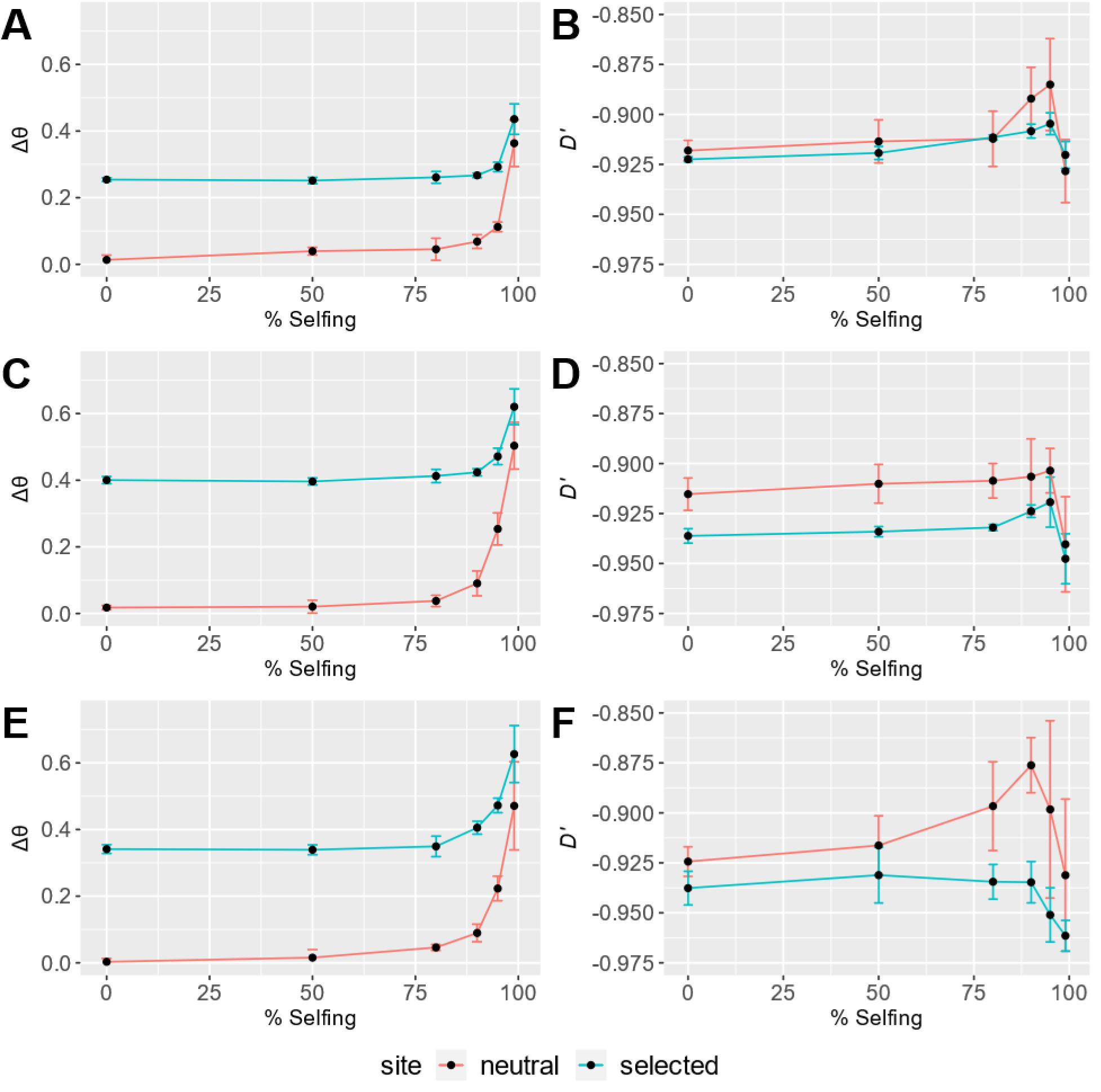
Effects of selfing on the SFS and linkage disequilibrium (LD) at neutral (red lines) and selected (blue lines) sites. The skew in the SFS was quantified using Δ*θ* for DFE1 (**A**), DFE2 (**C**), and DFE3 (**E**), calculated using alleles at all frequencies, while LD was summarized by the statistic *D*′ for DFE1 (**B**), DFE2 (**D**), and DFE3 (**F**), calculated for alleles at frequencies of 1%- 5%. The error bars denote the standard deviation of the genome-wide average of each summary statistic estimated from 5 independent replicates.

For selfing rates between 80% and 95%, *D*′ tended to become closer to 0 at both selected and neutral sites (Figure 4B). This is similar to an effect observed in simulations by Kamran-Disfani and Agarwal (2014), where partially selfing populations with intermediate selfing rates showed increases in linkage disequilibria. This phenomenon is expected at higher selfing levels if the efficacy of selection is not overwhelmed by HRI, because of correlations in homozygosity at different loci across the genome (Roze and Lenormand 2005). In contrast, *D*′ remained relatively flat for simulations with intermediate recombination rates (Supplementary Figure 10). At 99% selfing and at very low recombination rates, *D*′ values became considerably more negative for selected and neutral sites, consistent with a population experiencing pervasive HRI.

To understand how common summary statistics relate to errors in DFE misinference, we investigated how nucleotide diversity (*π*), *θH* (Fay and Wu 2000), singleton density (*sing*), haplotype diversity (*hapdiv*), and two LD statistics (*r*^2^and *D*) compared with the errors 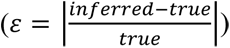 of the inferred parameters of the DFE (2*Ns̅*_*d*_ and *β*). In general, we found that SFS-based statistics as well as *hapdiv* at nonsynonymous sites divided by synonymous sites positively correlated with error in DFE inference (Supplementary Figure 11). On the other hand, LD-based statistics were not found to be particularly informative of the extent of misinference. However, we found that the values for each statistic vary depending on the DFE, suggesting that the value of any one summary statistic cannot be used to determine if pervasive interference is occurring in a population. Moreover, other factors such as assumptions about the fitness model, epistasis, and non-equilibrium demography will also affect the LD statistics (discussed in Johri and Charlesworth 2024), masking the effect of HRI.

### Simulation rescaling alters the effects of HRI

Due to computational time constraints, it was necessary to rescale the parameters *Nμ*, *Nr*, and *Ns* for most simulations in this study by lowering *N* and increasing *μ*, *r*, and *s* by the rescaling factor *Q* = 100. While equivalent equilibria are expected according to diffusion theory if the population-scaled parameters are held constant (Hill and Robertson 1966; Crow and Kimura, M. 1970; Ewens 1979; Li 1987), it has previously been observed that high levels of rescaling can increase the effects of HRI (Comeron and Kreitman 2002). To understand the effects of rescaling at high selfing rates, we once again simulated *C. elegans*-like populations with 500 genes and selfing rates of 95% and 99% and the three deleterious DFEs, with *Q* = 50, 20, and 10, though at *Q* = 10 only a limited number of simulations were able to be completed within a reasonable timeframe. We find that as *Q* decreases from the original value of 100, the effects of background selection and HRI decrease (Supplementary Figure 12). While the extent of DFE mis-inference decreases as *Q* is lowered, the DFEs estimated at low *Q* values and 99% selfing were still mis-inferred, especially the DFEs with more weakly and moderately deleterious mutations. The SFS at higher selfing rates and low *Q* was still skewed towards lower frequency mutations and U-shaped, though less than at high *Q* (Supplementary Figures 13-14). Thus, it is likely that fully unscaled simulations (*Q* = 1) would have even less HRI effects, especially for the strongly deleterious DFE, though it is not unrealistic to expect significant effects of BGS and interference between selected mutations based on the trends observed here. For instance, we observed substantially more mis-inference of the DFE when larger genomic regions were simulated (Supplementary Figure 15), suggesting that full length chromosomes simulated at *Q* = 1 will likely have more pervasive linked effects of selection. Indeed, Gilbert *et al*. (2022) conducted unscaled simulations of the entire genome, including all full-length chromosomes and a reduced recombination rate in chromosome centers, with *C. elegans-*like parameters and found a drastic decrease in population size due to background selection (see figure 4 of Gilbert *et al*.), so exploration of our scaled simulations with pervasive HRI is still relevant to highly selfing populations.

### Extent of mis-inference does not depend on coding density

The effects of HRI are expected to increase when there is a higher density of selected sites, and gene density varies along the *C. elegans* chromosomes, with higher gene densities found in the chromosome centers than in the chromosome arms (*C. elegans* Sequencing Consortium 1998). Moreover, the rate of recombination in *C. elegans* is higher in the gene-poor chromosome arms than in the gene-rich cores (Rockman and Kruglyak 2009). We however observed no obvious changes in the extent of DFE inference when coding density was 2× and 6× higher (Supplementary Figure 16), although local gene densities could increase more than we simulated.

### The impact of dominance on DFE inference

Dominance coefficients of new deleterious mutations vary considerably and are partially recessive on average, with strongly deleterious mutations tending to be highly recessive (Simmons and Crow 1977; Agrawal and Whitlock 2011; Manna et al. 2011). Most DFE inference methods assume that new mutations are semidominant and thus provide estimates of the joint distribution of the selection and dominance coefficients, “*s*_*d*_*h*”. A recent study by Wade *et al*. (2023) suggests that, if there are recessive lethal mutations, current DFE inference approaches can lead to underestimating the true proportion of strongly deleterious mutations, as more strongly deleterious mutations are likely to persist in the population if they are recessive. Furthermore, since selfing increases the levels of homozygosity in the genome, the efficacy of selection against large effect and highly recessive deleterious mutations is expected to be greater (Charlesworth and Charlesworth 1998), leading to more efficient purging of deleterious mutations. Thus, with higher rates of selfing one might instead expect to observe a decrease in the allele frequencies of mildly and moderately deleterious mutations, leading to the false inference of the presence of more strongly deleterious mutations in the population.

At 0% selfing, as new deleterious mutations became more recessive (especially when *h* = 0.1), the inferred distribution of the population-scaled product of selection and dominance coefficients slightly underestimated the proportion of strongly deleterious mutations (Supplementary Figure 17, Supplementary Table 5), consistent with Wade *et al*. (2023).

Additionally, the frequency of the nearly neutral class of mutations was underestimated in all three DFEs. In contrast, when more dominant mutations were simulated, DFE inference resulted in mis-inference towards both weakly and strongly deleterious mutations, though this effect was fairly small. Mis-inference was stronger when the true DFE was skewed towards mildly deleterious mutations, as these mutations are more likely to be present as homozygotes in the population.

As selfing increases homozygosity, the dominance coefficient is a less important factor for the efficacy of selection in highly selfing populations (Caballero and Hill 1992; Charlesworth 1992). Thus, at 50% selfing, the DFE was inferred more accurately (Supplementary Figure 18, Supplementary Table 5) and the dominance coefficient played a negligible role in DFE inference when the selfing rate was 99% (Supplementary Figure 17). Thus, at high rates of selfing, HRI effects were found to be primarily responsible for DFE mis-inference.

Increased purging of recessive deleterious mutations has been hypothesized to reduce their average allele frequencies, leading to DFE mis-inference (Arunkumar et al. 2015; Gilbert et al. 2022). However, observable effects of purging are likely to occur only when mutations are highly recessive (Nei 1968) and strongly deleterious (Glémin 2003). Concordantly, we observed that the distribution of allele frequencies of partially recessive (*h* = 0.25) mutations at intermediate selfing levels was the same as those of semidominant mutations (Supplementary Figure 19), suggesting that increased purging of recessive deleterious alleles did not have a substantial effect on the average allele frequencies in a population sample at equilibrium.

### Inference of the full DFE of new deleterious and beneficial mutations

Both BGS and HRI lead to a decrease in the efficacy of selection at linked selected sites, thus reducing the probability of fixation of beneficial mutations while increasing the probability of fixation of deleterious mutations (Charlesworth 1992; Kamran-Disfani and Agrawal 2014). It is therefore likely that the proportion of fixed mutations at selected sites that are beneficial (*α*) is mis-estimated when effective recombination rates are low. We conducted simulations with selfing, where 0.1% and 1% of new mutations in exons were beneficial, following an exponential distribution with 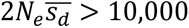. Due to recurrent selective sweeps in this scenario as well as BGS, there is likely to be a substantial decrease in effective population size experienced by selected alleles (and thus a reduced 2*N*_*e*_*s*_*d*_). Concordantly, the population-scaled selection coefficients of deleterious mutations were estimated to be heavily skewed towards mildly deleterious mutations even at much lower selfing rates (Supplementary Figures 20 and 21). When 0.1% of new exonic mutations were beneficial, and selfing rates were above 90%, the estimated distributions of selection coefficients were biased towards higher proportions of nearly neutral and strongly deleterious mutations, as observed previously (Figure 6A). However, the inferred distributions were skewed more towards strongly deleterious mutations compared to the simulations with only deleterious mutations. Mis-inference of selection coefficients was particularly substantial when the simulated deleterious DFE consisted of more weakly deleterious mutations, suggesting that HRI effects might cause mis-inference even at moderate levels of selfing if beneficial mutations are reasonably prevalent. The selected SFSs resembled the neutral SFSs in simulations with DFE mis-inference (Supplementary Figure 22). With a high rate of beneficial mutations entering the population, hitchhiking can reduce the efficacy of selection on weakly deleterious alleles (Campos and Charlesworth 2019), leading to the mis-inference of the DFE for the more weakly deleterious DFEs. In addition, the failure of DFE inference programs to fully correct for the presence of mildly beneficial alleles in the SFS could also have contributed to the observed mis-inference (Tataru et al. 2017).

**Figure 6:**
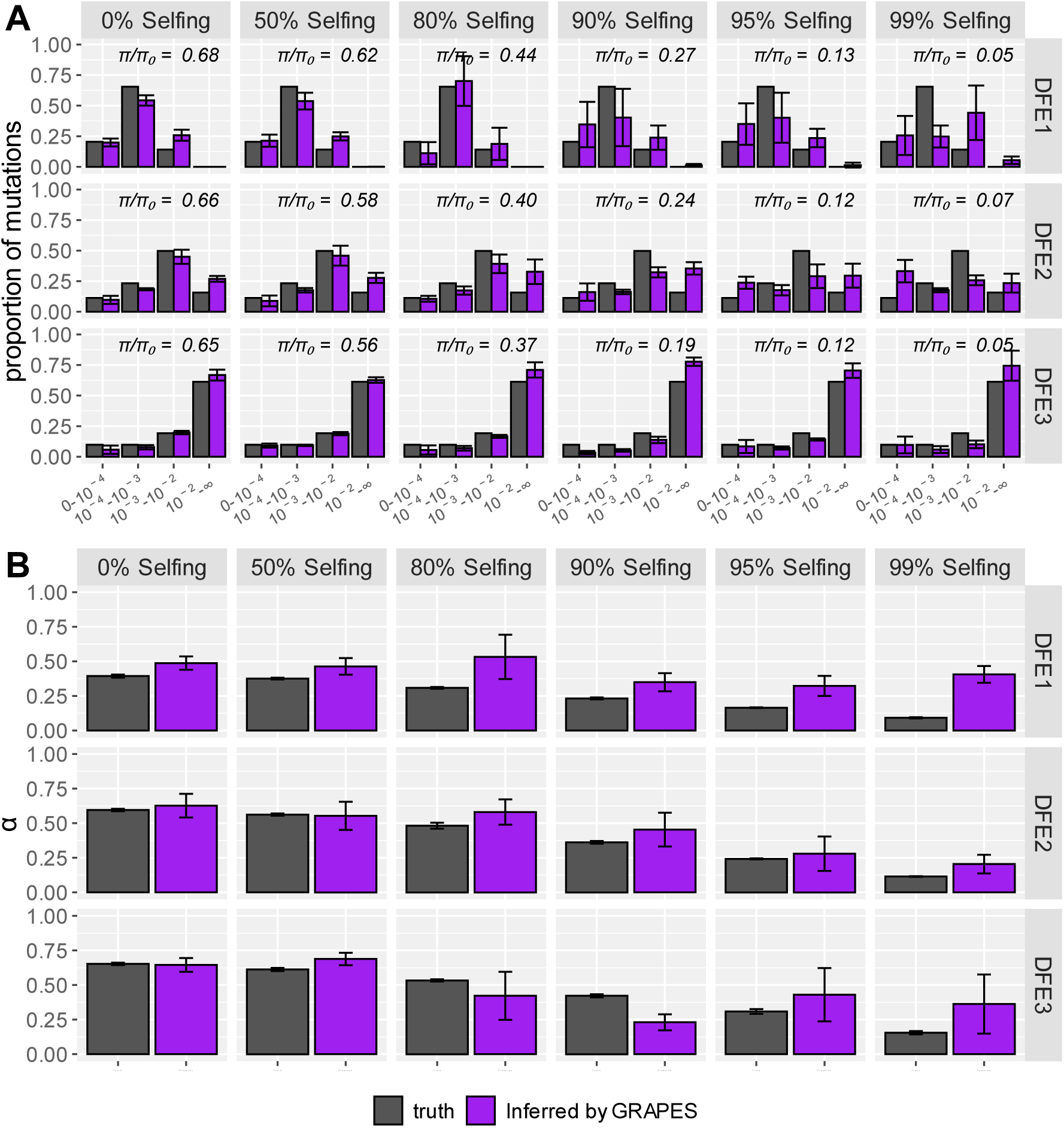
Effects of selfing on the inference of (A) the DFE of new deleterious mutations and (**B**) the proportion of beneficial fixations (α) using GRAPES (shown in purple bars). (**A**) Here the DFE is displayed as the distribution of *s*_*d*_, which was created by dividing the mean of the gamma distribution (2*N*_*e*_*s*_*d*_) by 2(*π*_*obs*_⁄*π*_*exp*_)*N*, where *π*_*obs*_ is the neutral nucleotide site diversity observed in each scenario and *π*_*exp*_is the neutral nucleotide site diversity expected in a population of size *N* at strict neutrality. Results are shown in terms of the proportion of mutations in effectively neutral (0 ≤ *s*_*d*_ < 1 × 10^−4^), weakly deleterious (1 × 10^−4^ ≤ *s*_*d*_ < 1 × 10^−3^), moderately deleterious (1 × 10^−3^ ≤ *s*_*d*_ < 1 × 10^−2^) and strongly deleterious (1 × 10^−2^ ≤ *s*_*d*_ < 1) classes. Selfing rates were varied between 0 and 99 %. *π*⁄*π*_0_, the neutral nucleotide diversity with selection relative to that expected under neutrality is shown. For both (**A**) and (**B**), 0.1% of new exonic mutations were beneficial and exponentially distributed with a mean 2*Ns*_*a*_ = 200. The error bars denote the standard deviation of proportions estimated from 5 independent replicates.

As the folded SFS might provide better estimates of the deleterious DFE when beneficial mutations are segregating at high frequencies (Keightley and Eyre-Walker 2007; Eyre-Walker and Keightley 2009), we also tested the inference using the folded SFS. Overall, the deleterious DFE inferred by GRAPES was comparable when using the unfolded and folded SFS (Supplementary Figures 23A & 24A). However, the DFEs estimated by DFE-alpha using the folded SFS were biased towards weakly deleterious mutations even with no selfing, with much larger uncertainty in the estimated DFE at high selfing levels, especially when 1% of all new selected mutations were beneficial. This indicates that folding the SFS is inadequate to correct for the mis-inference of the deleterious DFE when the rate of new beneficial mutations is extremely high.

As expected, the true fraction of beneficial substitutions (*α*) decreased as the selfing rate increased (Figure 6B, Supplementary Figure 20B). Changes in this parameter were driven by both increases in the number of deleterious substitutions (∼2 − 4 × higher at 99% selfing relative to 0% selfing) as well as the decreases in the number of beneficial substitutions (∼3 × lower at 99% selfing relative to 0% selfing; Supplementary Table 6). In simulations with both 0.1% beneficial mutations and 1% beneficial mutations, *α* tended to be overestimated for the weakly deleterious DFE at all selfing levels, with mis-inference increasing steadily as the selfing rate increased. For the moderately and strongly deleterious DFEs, *α* was only overestimated at high selfing levels, with the strongly deleterious DFE corresponding with the most accurate inference of *α*. This is consistent with the expectation that a decrease in the efficacy of selection will increase the fixation of weakly deleterious alleles, which will be misinterpreted as beneficial fixations by GRAPES, leading to an overestimation of *α* at high selfing levels. Very similar trends were observed when the folded SFS was used to estimate *α* (Supplementary Figures 23B & 24B). Overall, at high selfing levels, *α* values were substantially overestimated by GRAPES.

### Effects of population structure and sampling on the accuracy of inference

While selfing organisms tend to have highly structured populations, DFE inference in the context of population structure has been largely unexplored, though the effects of cryptic structure (without migration) were tested in Kim *et al*. (2017). Here, we simulated an island model, with five demes that exchanged migrants at a total rate *m* each generation. Multiple values of *m* were used to vary the degree of isolation between demes in this model, and the number of individuals in each deme (*N*_*deme*_) was varied so that the effective size of the metapopulation (*N*_*meta*_) was equivalent to 500,000 individuals (Supplementary Table 7). Under strict neutrality and while selfing was taken into account (see Methods), the neutral nucleotide diversity of the entire metapopulation (2*T*_*T*_*μ*), of individual subpopulations (2*T_S_μ*), and *F*_*ST*_ was equal to theoretical expectations (equations 2-4, Supplementary Table 8).

Population structure by itself did not result in mis-inference of the DFE when the rate of migration was sufficiently high (*i.e*., *N*_*deme*_*m* = 2, 1, and 0.5). High selfing rates with population structure again led to inferring a DFE (as population-scaled selection coefficients) skewed almost entirely towards mildly deleterious mutations, thus producing a very similar extent of mis-inference as that observed in a panmictic population (Supplementary Figure 25 A, B, and C, respectively). However, at low rates of migration between demes (*N*_*deme*_*m* of 0.1), the DFE was mis-inferred even at 0% selfing, with a bias towards inferring a lower efficacy of selection than the simulated DFE (Figure 7; Supplementary Figure 25D). This was largely because the relative increase in intermediate frequency alleles due to population structure as opposed to panmixia (Supplementary Figures 26-27) was falsely inferred as a bottleneck (mean fold change = 0.45-0.50; similar to Chikhi *et al*. 2010) by DFE-alpha, leading to incorrect estimates of the DFE. As expected, these distortions of intermediate frequency alleles in the SFS caused by population structure were observed to increase at higher selfing levels and at lower migration rates, with particularly large deviations from panmixia observed at *N*_*deme*_*m* = 0.1.

**Figure 7:**
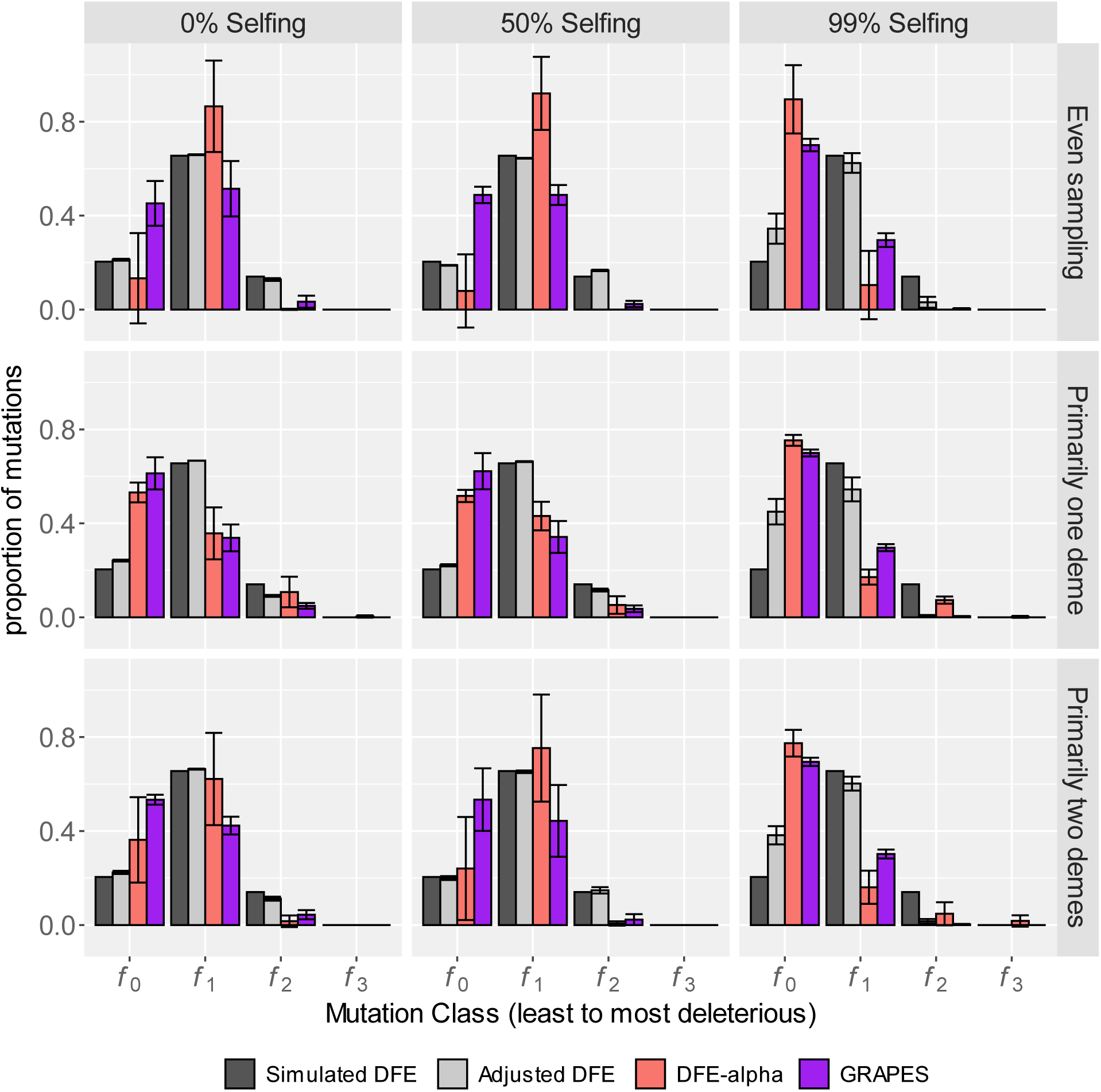
Effects of uneven sampling, population structure, and selfing on the inference of the DFE of new deleterious mutations using DFE-alpha (red bars) and GRAPES (purple bars). Results are shown in terms of the proportion of mutations in effectively neutral (*f*_0_), weakly deleterious (*f*_1_), moderately deleterious (*f*_2_) and strongly deleterious (*f*_3_) classes. Populations were simulated under an island model with five demes such that the metapopulation effective size was equal to 5000 at 0% selfing and *N*_*deme*_*m* was 0.1. Results are shown for the most weakly deleterious DFE (*i.e*., DFE1). The error bars denote the standard deviation of proportions estimated from 5 independent replicates. Rows represent the sampling scheme for genomes, where sampling was even, sampling was primarily from one deme, or sampling was primarily from two demes.

Empirical studies that draw from worldwide population samples may knowingly or unknowingly sample unevenly from structured populations when performing DFE inference. To investigate if this has unintended consequences on DFE inference, we repeated our experiments with population structure, altering our sampling strategy to primarily draw genomes from one deme (60%) or two demes (35% from each), with the remaining demes contributing only 10% of the sampled genomes each. As expected, the site frequency spectra from both unevenly sampled populations were observed to be even more distorted at intermediate frequencies than the evenly sampled populations with low migration rates (Supplementary Figures 28-31). For the sampling strategy primarily drawing from one deme, some DFE mis-inference towards nearly neutral and strongly deleterious mutations was observed at 0% and 50% selfing even for *N*_*deme*_*m* = 1 and more strongly at *N*_*deme*_*m* = 0.5, and this effect was stronger at the DFE with predominantly weakly deleterious mutations (Supplementary Figure 32). This effect was also observed for the sampling strategy primarily drawing from two demes, but to a lesser extent (Supplementary Figure 33). At the lowest migration rate, the inferred DFEs for both uneven sampling cases were substantially less deleterious at 0% and 50% selfing than for even sampling, particularly for the DFE with predominantly weakly deleterious mutations (Figure 7). At higher selfing levels, the inferred DFEs for all sampling schemes falsely inferred a larger proportion of weakly deleterious mutations than predicted by the DFE corrected for BGS (Figure 7), likely due to a combination of HRI and uneven sampling. The BGS-corrected DFE was unable to correct for the mis-inference of the DFE for all selfing levels at the lowest migration rate, reflecting the fact that DFE mis-inference is likely driven by changes in the SFS caused by population structure rather than changes in the level of BGS.

Overall, these results indicate both DFE inference approaches are fairly robust to population structure in an island model, even with uneven sampling, until the populations become extremely isolated from each other, or individuals are mostly drawn from one deme, especially when the true DFE has a significant proportion of mildly deleterious mutations.

Additionally, both low rates of migration and highly uneven sampling result in the inference of a DFE even more skewed towards weakly deleterious mutations than simulated. Testing DFE inference with more models of population structure and additional sampling strategies might be necessary to interpret empirically estimated DFEs from natural populations.

## DISCUSSION

Inference of the distribution of fitness effects (DFE) of new mutations is a critical step in understanding the forces that shape genetic variation in a population. Population genetic methods to infer the DFE that account for nonequilibrium demography usually assume independent segregation of putatively neutral and selected sites (Eyre-Walker and Keightley 2007; Keightley and Eyre-Walker 2007; Galtier 2016; Tataru et al. 2017). However, organisms that have little/ no recombination, for instance, those that undergo high rates of selfing or asexual reproduction, are likely to violate this assumption. A major consequence of strong linkage is the increase in HRI effects that lower the efficacy of selection. When the regime of classical BGS and sweeps is predominant, inferred population-scaled selection coefficients are biased, but accounting for the relative decrease in the rate of coalescence can provide reasonably accurate estimates of the selection coefficients. However, when interference between selected alleles is pervasive, the estimation of the distribution of selection coefficients is also severely mis-inferred. We also find that while purging of partially recessive deleterious mutations does not lead to mis-inference, the presence of cryptic population structure where there is a low rate of migration between subpopulations and/or sampling from predominantly one subpopulation leads to inferring a DFE (in terms of the population-scaled coefficients) skewed towards effectively neutral/mildly deleterious class of mutations. Moreover, the proportion of beneficial substitutions (*α*) is substantially overestimated in highly selfing species, suggesting caution when performing inference in species or genomic regions with little/no recombination.

### Why is the DFE estimated from *C. elegans* populations not skewed towards mildly deleterious mutations?

While our study suggests that DFE inference in highly selfing species should yield population-scaled selection coefficients skewed towards weakly/moderately deleterious mutations, surprisingly, the previous empirical estimate of the *C. elegans* DFE using DFE-alpha is heavily skewed towards the class of strongly deleterious mutations (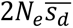 > 10,000, Gilbert *et al*.2022). One explanation of this discrepancy is that the mean 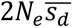 of the population is even greater than 10,000. Interestingly, direct measurements of fitness-related traits in mutation accumulation lines suggest that *C. elegans* has a highly deleterious DFE, with estimates of the mean deleterious selection coefficient ranging from -0.09 to -0.46 (Vassilieva et al. 2000; Baer et al. 2005; Katju et al. 2015). Thus, using previously calculated *C. elegans N*_*e*_ ranging from 5,000 to 80,000 (Table 4 in Teterina *et al*. 2023), the expected 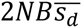 would range from 900 to 72,000, consistent with this explanation. However, there are a few major caveats to these observations. (a) The experimental estimates of selection coefficients are likely biased towards more deleterious selection coefficients due to the difficulty of measuring small fitness changes in experimental lines (Davies et al. 1999; Eyre-Walker and Keightley 2007), and might not reflect fitness effects in the wild. (b) It is unclear whether previous estimates of the population sizes of *C. elegans* populations using neutral nucleotide diversity (Barrière and Félix 2005; Cutter 2006; Rockman and Kruglyak 2009; Teterina et al. 2023) are accurate due to the joint effects of selfing, HRI, and demography (c) Empirical estimates of the deleterious DFE in *C. elegans* (Gilbert *et al*. 2022) were obtained by sampling worldwide rather than from local populations, and thus other factors such as complex demographic history and population structure could play a role as well. For instance, selfing populations are likely to follow extinction-recolonization dynamics (Charlesworth and Wright 2001; Ingvarsson 2002; Wright et al. 2013) and such models remain understudied.

A second explanation for the discrepancy between the expected and observed DFE in *C. elegans* is that the true selfing rate in wild *C. elegans* might be lower than 99%, thus reducing linked effects of selection. Though selfing rate estimates from unique populations range from 80% to ∼100% using various methods (Barrière and Félix 2005; Sivasundar and Hey 2005; Teterina et al. 2023), in general, the high LD in *C. elegans* populations (haplotype blocks of 2.5 Mb on average; Cutter 2006) along with low frequency of males in wild specimens (Barrière and Félix 2005; Richaud et al. 2018) suggest the outcrossing rate is very low in most populations. A recent study using whole-genome sequencing data focusing on Hawaiian *C. elegans* estimated a relatively lower selfing rate of 93%, suggesting that selfing rate can vary across populations (Teterina et al. 2023). Importantly, as the samples used in Gilbert *et al*. (2022) were unevenly sampled worldwide (with many samples from Hawaiian populations), variation in selfing rate across populations or through time could additionally influence DFE inference results.

A third possibility relates to the difficulty in obtaining the SFS using highly inbred strains. The SNP-calling pipeline used to obtain *C. elegans* population-genetic data converted heterozygous sites into homozygous reference or alternate alleles depending on if the mutant was above or below 50% (Cook et al. 2017). This method of SNP processing could lead to the inference of a more deleterious DFE if the heterozygote frequency is higher at neutral sites than selected sites by skewing the neutral SFS towards higher frequencies and the selected SFS towards lower frequencies, though the low frequency of heterozygotes in highly selfing populations makes this unlikely to be a large issue. Future studies working with inbred strains should evaluate the effect of employing such SNP processing approaches on population genomic inference.

Finally, we have only examined a population under equilibrium in the current study, whereas a recent population expansion/ contraction could decrease the observed effects of interference between selected alleles at multiple loci, allowing for a correct inference of selection coefficients. HRI effects in non-equilibrium populations remain to be studied more thoroughly.

Unfortunately, as linked effects of selection will also lead to falsely inferring recent population growth (Supplementary Table 1; Ewing and Jensen 2016; Schrider *et al*. 2016; Johri *et al*. 2021), there is no straightforward solution to accounting for the effects of the true demography (Johri, Eyre-Walker, et al. 2022). Thus performing a joint inference of demography and the DFE as employed by Johri *et al*. (2020) is likely to be the most promising approach to circumvent this challenge.

### How broadly do our results apply?

Our results have implications for the population genetics of other highly selfing organisms, many of which could fall into a parameter space where linked effects of selection are pervasive. A key factor that determines the extent of HRI is the number (*U*_*d*_) of new deleterious mutations per diploid genome per generation, with higher values of *NU*_*d*_ relative to *NR* causing more interference. Thus, the extent of HRI across species would likely differ due to differences in outcrossing rates, population sizes, mutation and recombination rates, and genome architecture (McVean and Charlesworth 2000; Comeron and Kreitman 2002; Kaiser and Charlesworth 2009; Good et al. 2014). Our Figure 3 suggests that the extent of background selection (measured by *B*) is a good predictor of the possible extent of HRI and that *B* < 0.5 could result in substantial misinference. Interestingly, a number of highly selfing species may have reasonably low *B* values (Table 2). In addition to *C. elegans*, a variety of species such as *C. briggsae*, *A. tauschii*, *A. thaliana*, *C. bursa-pastoris*, *C. orientalis,* and *S. lycopersicon* have their lowest *B* value estimates below 0.25, with *C. elegans, C. briggsae,* and *S. lycopersicon* also exhibiting values of *U*_*d*_⁄*R*_*self*_ > 5. Interestingly, *C. elegans* has been observed to have a U-shaped SFS in many samples (Freund et al. 2023), consistent with HRI, though in empirical data a variety of explanations such as ancestral allele misorientation, positive selection, and various demographic effects could explain a U-shaped SFS (Freund et al. 2023). This could pose problems for methods that infer the dominance coefficients of mutations by comparing the DFEs of closely related selfing and obligate outcrossing species, such as a recent study of *Arabidopsis* (Huber et al. 2018), and thus efforts to quantify the levels of HRI in highly selfing species merits additional investigations. In addition to selfing organisms, pathogenic species like *Plasmodium* (*U*_*d*_ = 0.25; Lynch *et al*. 2016) that have variable rates of sexual reproduction between genetically identical clones (and thus experience reduced rates of recombination), seem to have nearly identical neutral and selected site frequency spectra (Figure 3 in Parobek *et al*. 2016), suggesting that HRI effects might play an important role in shaping variation across their genomes. We therefore also expect DFE inference to be biased due to HRI in asexual organisms, mitochondrial genomes, and the Y chromosome, all of which have low effective recombination rates (Barton and Charlesworth 1998; Charlesworth and Charlesworth 2000; Weinreich and Rand 2000). Importantly, as structural variants (Adrion et al. 2017; Abel et al. 2020) and mutations in noncoding regions (Racimo and Schraiber 2014) are a significant source of deleterious mutations accounting for the input of other mutation types is likely to increase the estimated value of *U*_*d*_, making HRI more likely in populations with little/no recombination.

**Table 2:** Estimated levels of background selection (*B*) in highly selfing species for the range of previously reported selfing rates. All species are diploid except tetraploid *C. bursa-pastoris*. *U*_*d*_is the number of deleterious mutations per diploid genome per generation, while *R* is the map length in Morgans for each genome. The strength of BGS was calculated with 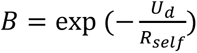, where *R*_*self*_ = *R* × *F*. All mutation rates, recombination rates, and genome annotation information used to construct this table is present in Supplementary Table 9.

| Species | Coding fraction ($f_{coding}$) | $\frac{U_d}{R}$ | Selfing rate | $\frac{U_d}{R_{self}}$ | Estimated $B$ | References |
| --- | --- | --- | --- | --- | --- | --- |
| *Aegilops tauschii* * | 0.01 | 0.022 | 0.90-0.995 | 0.25-4.45 | 0.01-0.78 | (Willenborg and Van Acker 2008; Luo et al. 2013; Wang et al. 2021; Burgarella et al. 2024) |
| *Arabidopsis thaliana* | 0.27 | 0.071 | 0.97-0.99 | 1.32-3.89 | 0.02-0.27 | (Snape and Lawrence 1971; Singer et al. 2006; Ossowski et al. 2010; Cheng et al. 2017) |
| *Caenorhabditis briggsae* * <sup>+</sup> | 0.22 | 0.031 | 0.95-0.999 | 0.33-15.50 | ~0-0.72 | (Zeitler and Gilbert 2024) |
| *Caenorhabditis elegans* | 0.3 | 0.042 | 0.93-0.999 | 0.32-21.00 | ~0-0.73 | (*C. elegans* Sequencing Consortium 1998; Denver et al. 2004; Saxena et al. 2019; Teterina et al. 2023) |
| *Capsella bursa-pastoris*<sup>+</sup> | 0.21 | 0.077 | 0.90-0.99 | 0.42-3.87 | 0.02-0.66 | (Linde et al. 2001: 200; Wesse et al. 2021; Penin et al. 2024) |
| *Capsella orientalis*<sup>+</sup> | 0.24 | 0.086 | 0.90-0.99 | 0.48-4.36 | 0.01-0.62 | (Ågren et al. 2014) |
| *Medicago truncatula* | 0.13 | 0.096 | 0.92-0.95 | 0.65-1.01 | 0.36-0.52 | (Maureira-Butler et al. 2008; Gorton et al. 2012; Pruitt et al. 2014; Jullien et al. 2021) |
| *Oryza sativa (indica)* | 0.12 | 0.017 | 0.95-0.99 | 0.17-0.83 | 0.43-0.84 | (Pruitt et al. 2014; Yang et al. 2015: 20; De Leon et al. 2016; Bah et al. 2017) |
| *Solanum lycopersicon* | 0.06 | 0.029 | 0.948-0.999 | 0.29-14.10 | ~0-0.75 | (Roselius et al. 2005; Viquez-Zamora et al. 2014; Horneburg and Becker 2018; Su et al. 2021) |
| *Triticum urartu* * <sup>+</sup> | 0.08 | 0.026 | 0.90-0.991 | 0.14-1.45 | 0.23-0.87 | (Willenborg and Van Acker 2008; Ling et al. 2018; Burgarella et al. 2024) |

\* mutation rate estimate of closely related species used for calculations (*A. thaliana* for plants, *C. elegans* for *C. briggsae*)
<sup>+</sup>recombination rate of closely related species used for calculations

### Considerations for inferring the DFE in highly selfing organisms

Given our observations, we here provide a few points to consider when inferring the DFE of new mutations in organisms with little/no recombination, where effects of BGS, selective sweeps, or HRI might be pervasive:

1. The DFE of new deleterious mutations estimated using the two-step population-genetic methods is most closely explained by the distribution of 2*NBs*_*d*_, which is equivalent to 2*N*_*emp*_(1 + *F*)*s*_*d*_ in partially selfing populations. Here *N*_*emp*_ represents the effective population size estimated using neutral nucleotide diversity, which includes the effects of BGS and is what is usually estimated using demographic inference approaches that do not account for linked effects of selection. Thus, if there exist experimentally validated estimates of selfing rates, it is possible to accurately infer the distribution of selection coefficients in populations with moderate levels of selfing. However, in many organisms selfing rates are inferred using allele frequencies and those estimates are likely to be biased by population history and effects of selection at linked sites. Moreover, this raises the question of how one can accurately simulate an estimated DFE (in terms of population-scaled selection coefficients) when multiple linked selected sites are to be simulated, which will be especially important to determine the extent of HRI effects in populations with little/no recombination.
2. Different models of population structure and the effects of demographic changes need to be considered to fully understand the robustness of DFE inference. For instance, it is known that local populations of selfing organisms are prone to recurrent extinction and recolonization events (Pannell and Charlesworth 1999; Ingvarsson 2002) which may bias DFE inference further. Indeed, recent field studies of *C. elegans* populations suggest that 3-10 individuals colonize a food source and quickly reproduce to census sizes of up to 10,000 individuals before dispersing in search of new food sources (Barrière and Félix 2005; Richaud et al. 2018). When sampling across many demes, we suggest also inferring the DFE in isolated populations to minimize biases due to cryptic population structure or uneven sampling and to downsample the number of genomes such that sampling is as even as possible across demes. Sampling strategies such as a scattered sampling scheme may be useful in removing such bias (Wakeley 1999; Städler et al. 2009), though the reduction in sample size may reduce the ability to accurately infer the moderate and deleterious portions of the DFE (Kim et al. 2017).
3. If possible, it could be helpful to infer the DFE in closely related non-selfing species or in different populations of the same species with known differences in selfing rates. For instance, *C. elegans* and *A. thaliana* populations vary in the estimated rates of selfing by ∼10%, and the mostly outcrossing *A. lyrata* has several populations that have evolved partial selfing (Mable et al. 2005; Bomblies et al. 2010; Teterina et al. 2023). Though differences such as population structure and demography need to be carefully accounted for, statistical comparisons across populations/species with flexible DFE inference frameworks (Sendrowski and Bataillon 2024) could also be used to approximate the extent of HRI in a selfing species.
4. Finally, in the cases where HRI is determined to be extensive, joint inference of demography and the DFE as employed by Johri *et al*. in *D. melanogaster* (2020) and human (2022) populations could be especially useful for DFE inference. They employed an approximate Bayesian computation (Beaumont et al. 2002) framework combined with forward simulations to jointly infer parameters of the DFE of new deleterious mutations and an arbitrary size change, while accounting for the effects of selection at linked sites. While highly successful, such an approach has been limited to a relatively simple demographic model of a single size change in one population due to the large number of parameters required to be jointly estimated. In addition, our work here indicates that simulation rescaling may have to be limited to accurately capture the dynamics of Hill-Robertson interference, which will further increase the computational burden of simulations. However, if computationally feasible, such an approach could yield the most likely distribution of selection coefficients, especially because, the distortions of the SFS from a neutral model caused by interference cannot be explained by any constant *N*_*e*_ (see discussion in Neher 2013; Good *et al*. 2014; Melissa *et al*. 2022). Thus, until theoretical models can accurately estimate the effects of HRI in organisms with little/no recombination with complex DFEs, simulation-based approaches will be the most helpful in inferring the DFE of new mutations in such populations, as well as obtaining the fixation probability of new mutations, which will be altered by HRI.

## ACKNOWLEDGEMENTS

We thank Brian Charlesworth for many helpful discussions related to the project and for providing suggestions and comments that improved the manuscript. We are also thankful to Matthew Hartfield for providing helpful comments to improve the manuscript and to Matt Rockman for many helpful discussions and for pointing us to relevant resources available for *C. elegans*. We thank an anonymous reviewer and Thomas Batallion for their helpful suggestions and for improving the manuscript. The research in this study was conducted using computational resources provided by ITS Research Computing at the University of North Carolina at Chapel Hill. A.T.D. received support from NIGMS predoctoral training grant 5T32 GM067553.

Research reported in this publication was supported by the National Institute of General Medical Sciences of the National Institutes of Health under award number R35GM154969 to P.J. The authors declare no conflicts of interests.

## DATA AVAILABILITY

The following data are publicly available on https://github.com/JohriLab/DFESelfing: (1) Data and scripts to reproduce figures; (2) Config files used for DFE inference; (3) Simulation outputs in SLiM format; (4) templates and example scripts for launching SLiM simulations.

## SUPPLEMENTARY FIGURES AND TABLES

**Supplementary Figure 1:**
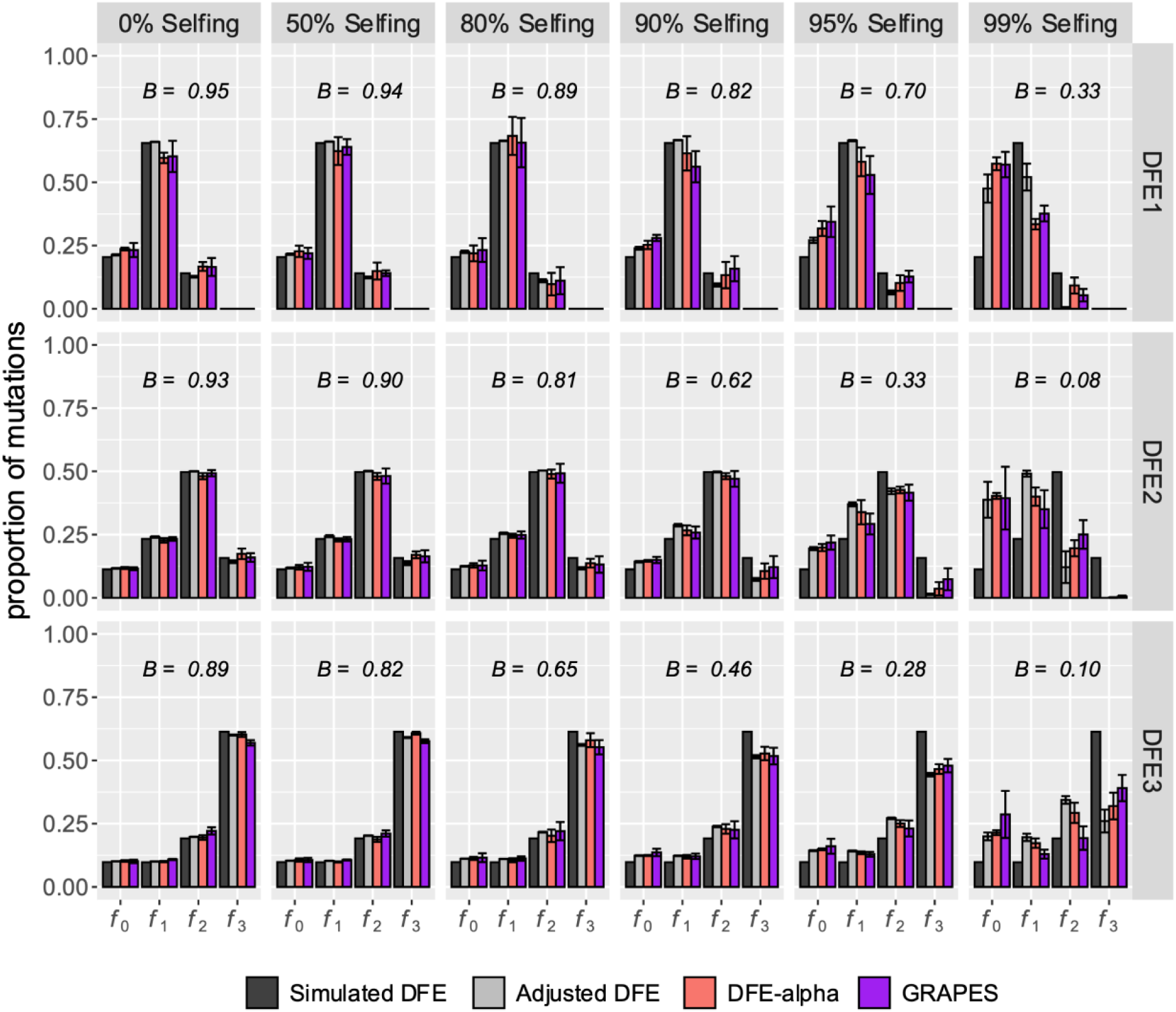
Effects of selfing on the inference of the DFE of new deleterious mutations using DFE-alpha (red bars) and GRAPES (purple bars). Results are shown in terms of the proportion of mutations in effectively neutral (*f*_0_), weakly deleterious (*f*_1_), moderately deleterious (*f*_2_) and strongly deleterious (*f*_3_) classes. Selfing rates were varied between 0 and 99 %. The nucleotide site diversity with background selection (*B*) relative to its expectation under strict neutrality is shown in each panel. The simulated (black bars) and adjusted (grey bars) DFE represents the distribution of 2*Ns*_*d*_ and 2*NBs*_*d*_ respectively. The error bars denote the standard deviation of proportions estimated from 5 independent replicates.

**Supplementary Figure 2:**
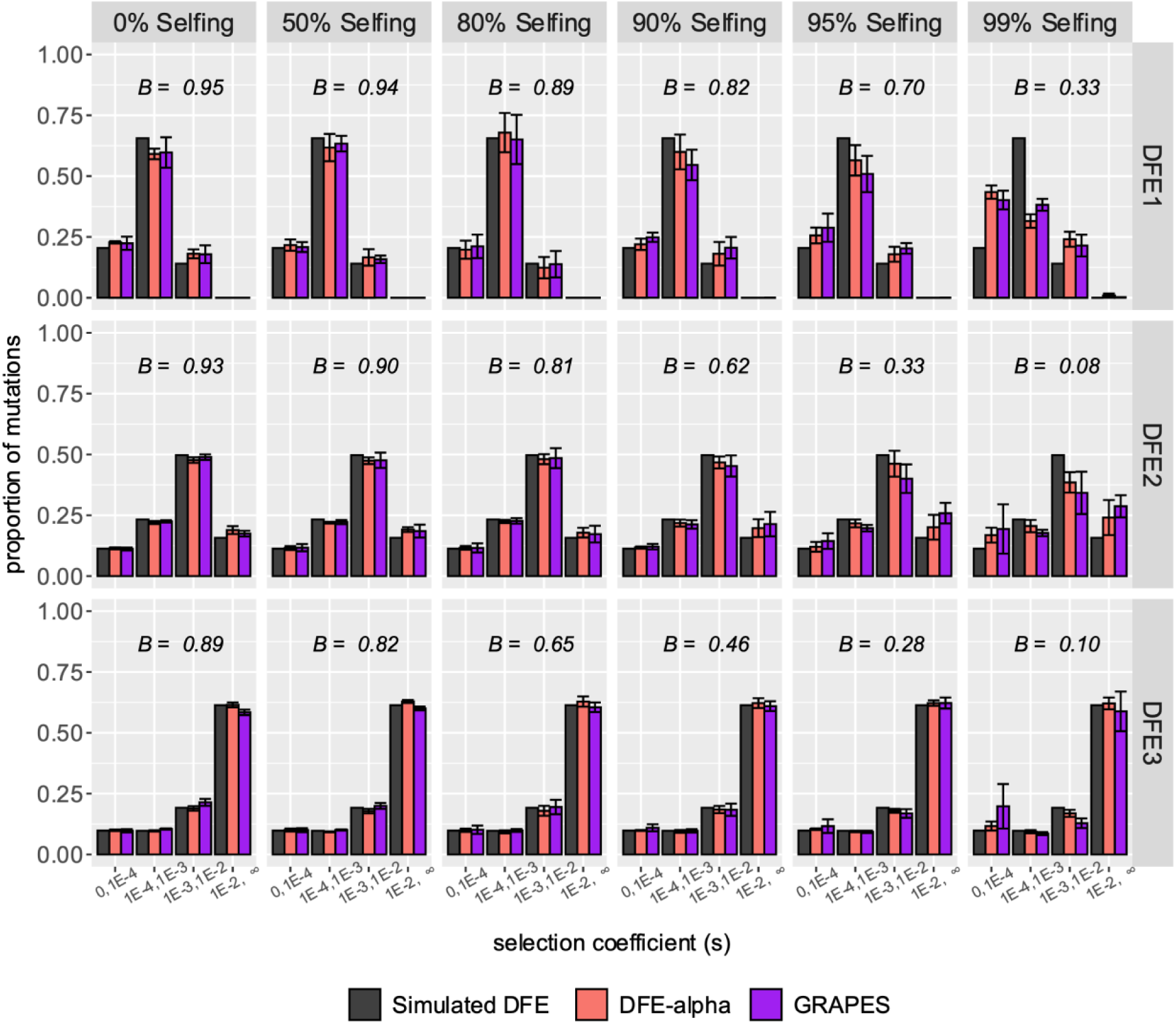
Effects of selfing on the inference of the DFE of new deleterious mutations using DFE-alpha (red bars) and GRAPES (purple bars). Here the DFE is displayed as the distribution of *s*_*d*_, which was created by dividing the mean of the gamma distribution (2*N*_*e*_*s*_*d*_) by 2*BN*. Results are shown in terms of the proportion of mutations in effectively neutral (0 ≤ *s*_*d*_ < 1 × 10^−4^), weakly deleterious (1 × 10^−4^ ≤ *s*_*d*_ < 1 × 10^−3^), moderately deleterious (1 × 10^−3^ ≤ *s*_*d*_ < 1 × 10^−2^) and strongly deleterious (1 × 10^−2^ ≤ *s*_*d*_ < ∞) classes. Selfing rates were varied between 0 and 99 %. The black bars represent the simulated distribution of *s*_*d*_. The error bars denote the standard deviation of proportions estimated from 5 independent replicates.

**Supplementary Figure 3:**
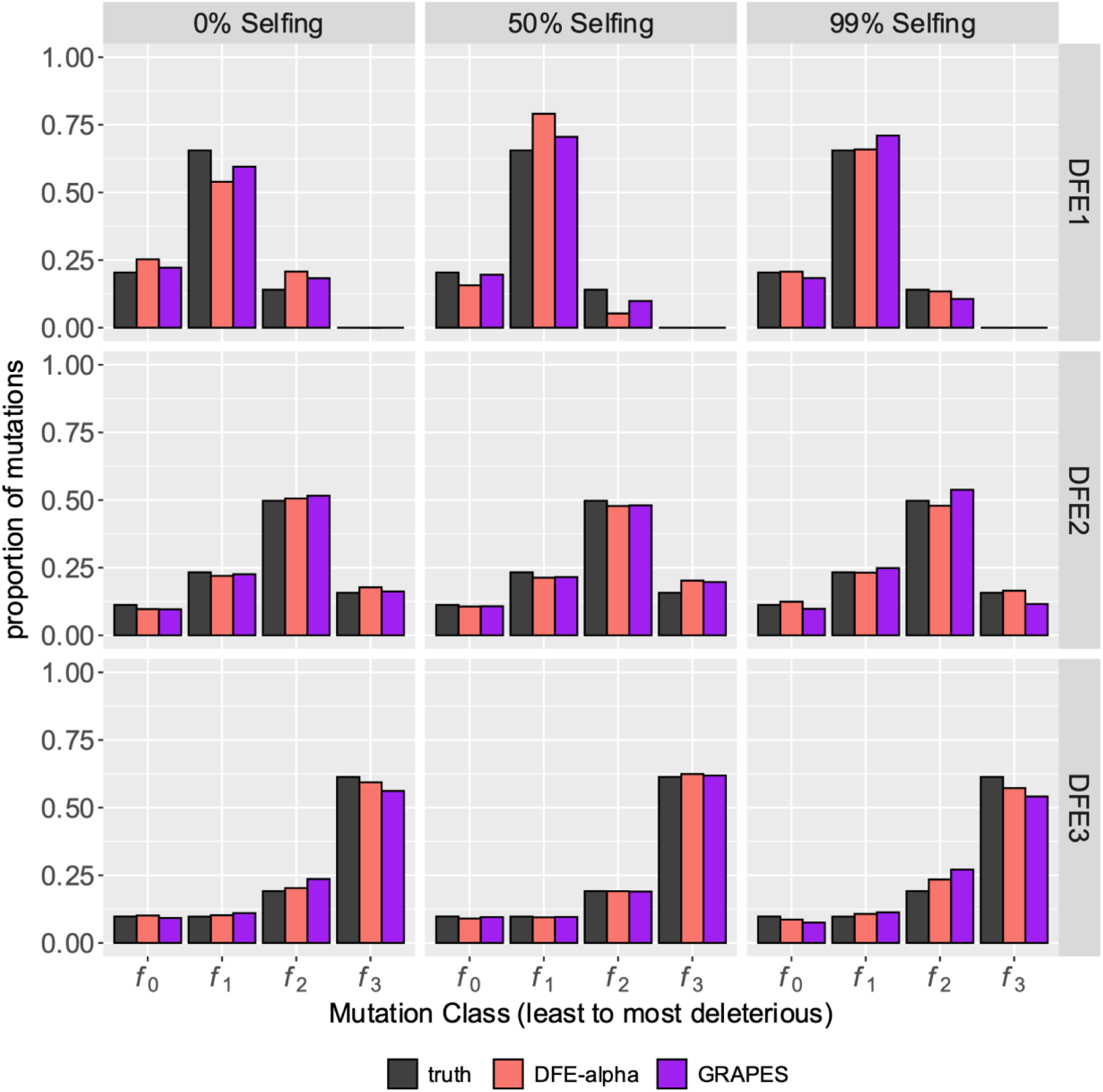
Inference of the DFE of new deleterious mutations in partially selfing populations using DFE-alpha and GRAPES for simulations with no linkage. Here, 50,000 selected mutations were simulated individually (as single locus simulations) in various selfing rates without linkage to other selected mutations. Results are shown in terms of the proportion of mutations in effectively neutral (*f*_0_), weakly deleterious (*f*_1_), moderately deleterious (*f*_2_) and strongly deleterious (*f*_3_) classes.

**Supplementary Figure 4:**
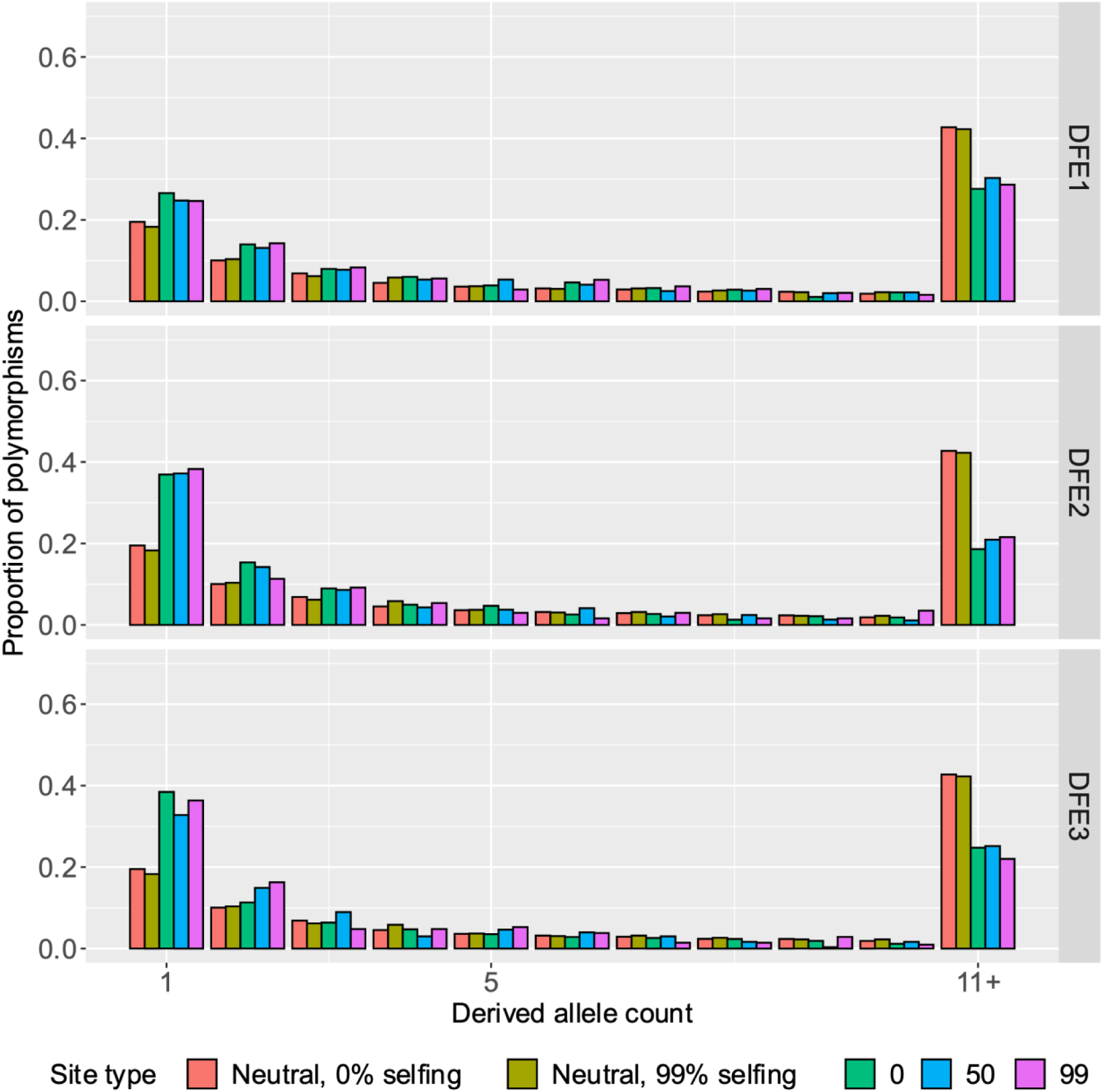
Site frequency spectra of neutral and selected alleles in partially selfing populations from single-site simulations. 100 genomes were sampled and all selected mutations were additive and deleterious The *y*-axis represents the proportion of segregating polymorphisms that fall into the given allele-count class. The last class (11+) refers to the derived allele counts 11-100. The SFS of neutral sites is shown in red, while the SFS of selected sites in simulations with 0%, 50%, and 99% selfing are shown as green, blue, and purple bars respectively.

**Supplementary Figure 5:**
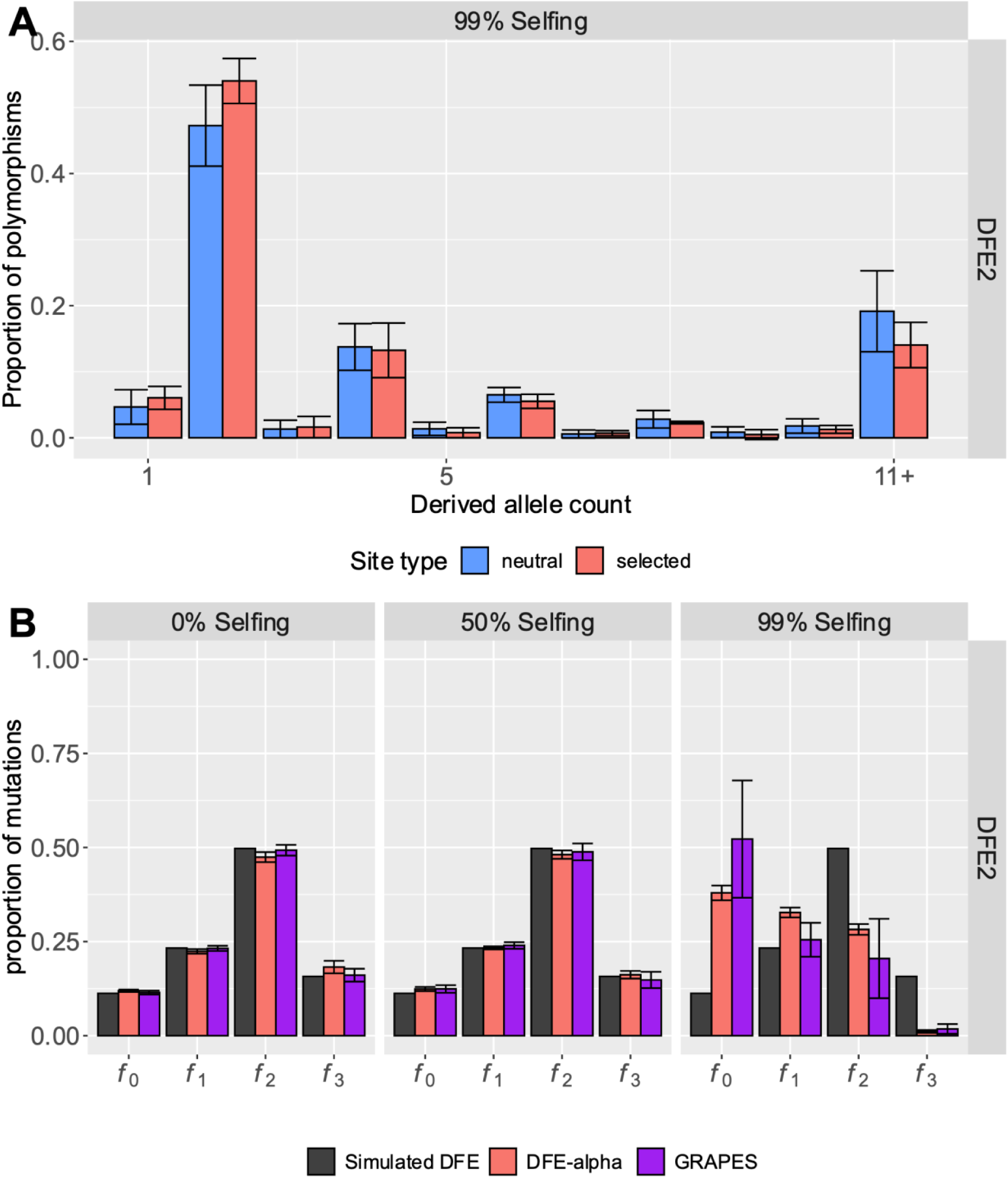
(A) The SFS at neutral (blue) and selected (red) sites when 50 diploid individuals are sampled (instead of 100 random genomes). (B) The accuracy of DFE inference when 50 diploid individuals are sampled (instead of 100 random genomes). Results are shown for a predominantly moderately deleterious DFE (DFE2). Results are shown in terms of the proportion of mutations in effectively neutral (*f*_0_), weakly deleterious (*f*_1_), moderately deleterious (*f*_2_) and strongly deleterious (*f*_3_) classes. The error bars denote the standard deviation of proportions estimated from 5 independent replicates.

**Supplementary Figure 6:**
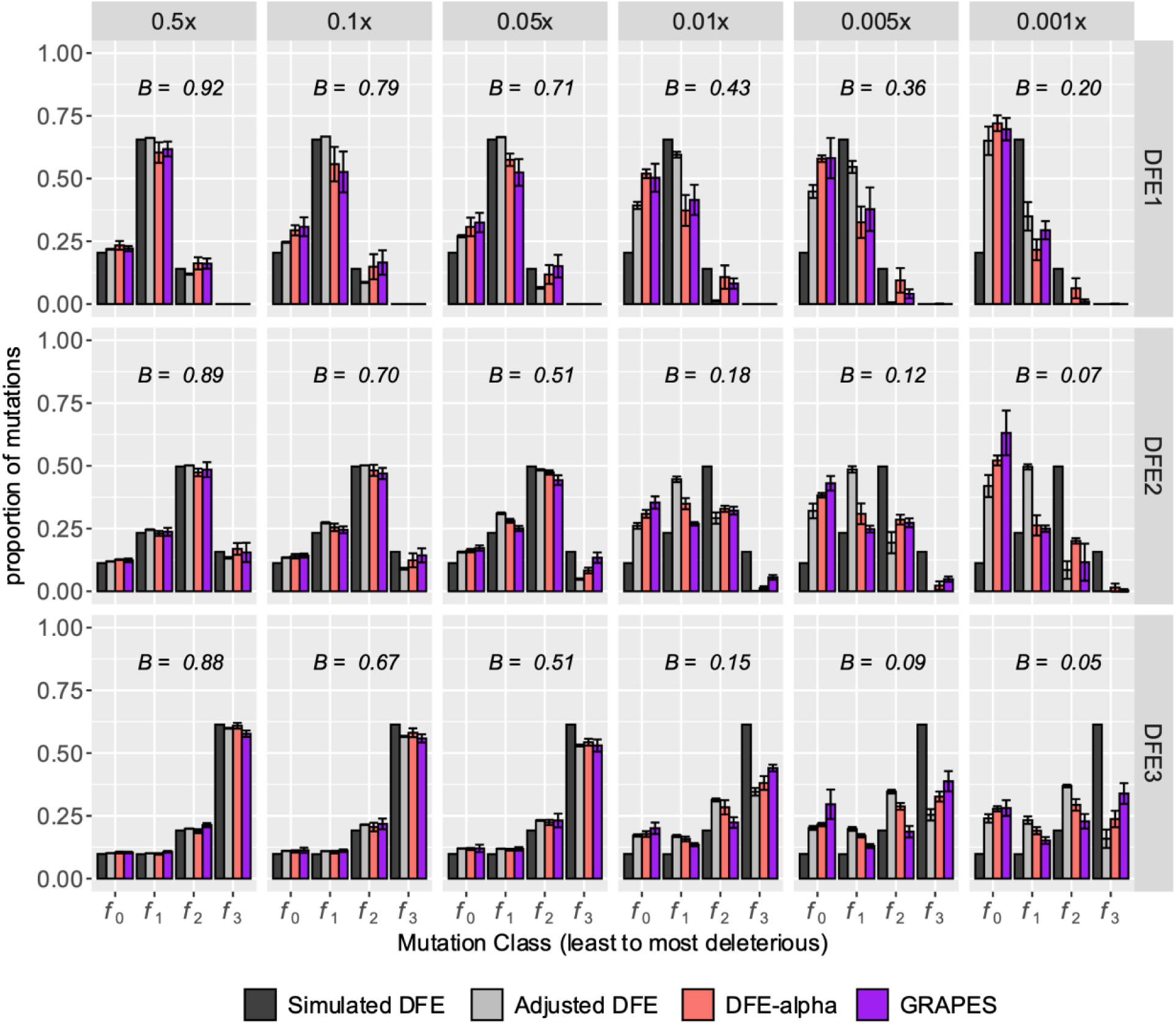
Effects of low recombination rate on the inference of the DFE of new deleterious mutations using DFE-alpha (red bars) and GRAPES (purple bars). Results are shown in terms of the proportion of mutations in effectively neutral (*f*_0_), weakly deleterious (*f*_1_), moderately deleterious (*f*_2_) and strongly deleterious (*f*_3_) classes. Recombination rates were varied between 0.5 × mean and 0.001 × mean, where the mean recombination rate was 3.12 × 10^−8^ per site/generations. The nucleotide site diversity with background selection (*B*) relative to its expectation under strict neutrality is shown in each panel. The simulated DFE (shown in black) and adjusted DFE (shown in grey) represents the distribution of 2*Ns*_*d*_ and 2*NBs*_*d*_ respectively. The error bars denote the standard deviation of proportions estimated from 5 independent replicates.

**Supplementary Figure 7:**
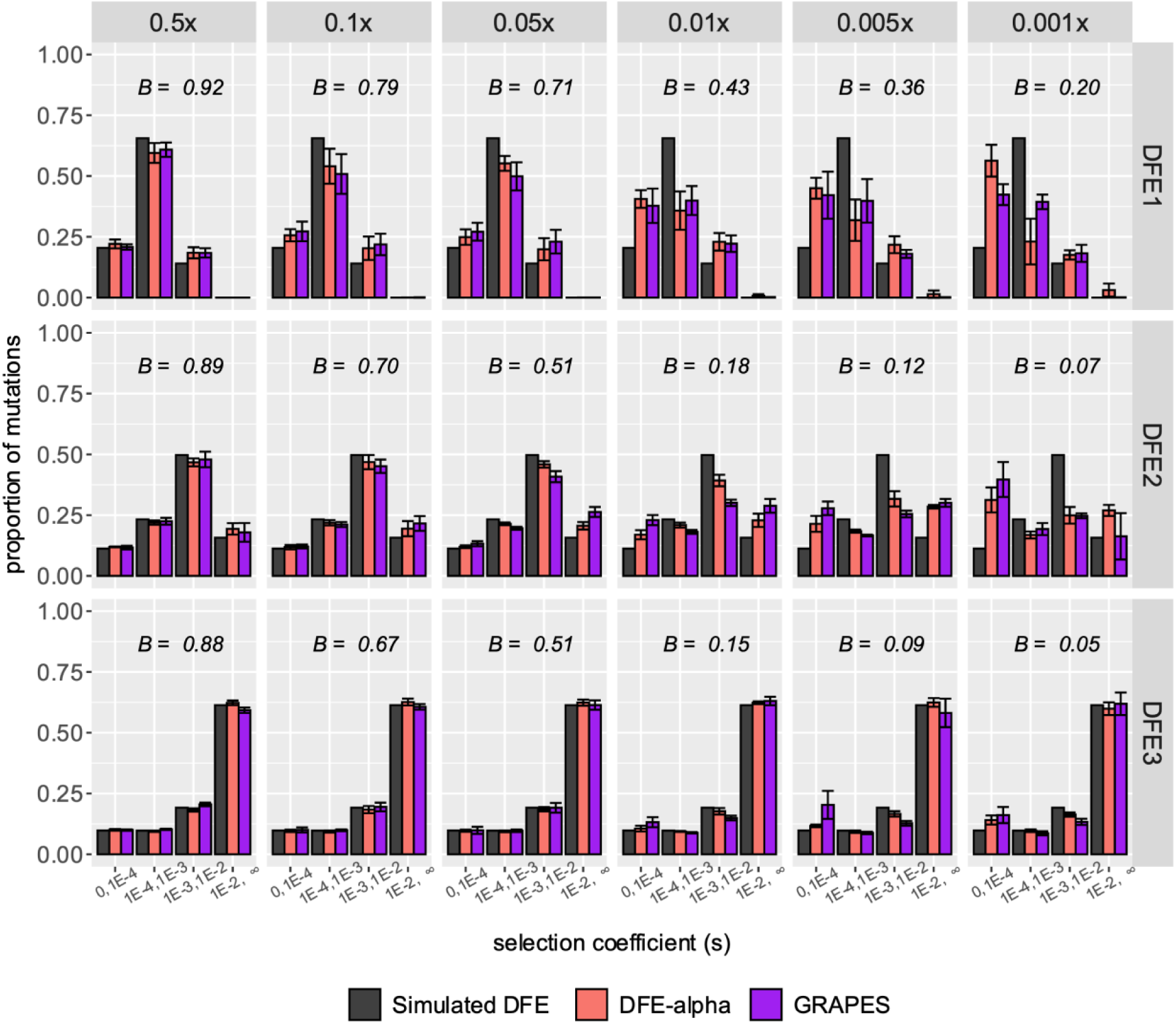
Effects of low recombination rate on the inference of the DFE of new deleterious mutations using DFE-alpha (red bars) and GRAPES (purple bars). Here the DFE is displayed as the distribution of *s*_*d*_, which was created by dividing the mean of the gamma distribution (2*N*_*e*_ *s*_*d*_) by 2*BN*. Results are shown in terms of the proportion of mutations in effectively neutral (0 ≤ *s*_*d*_ < 1 × 10^−4^), weakly deleterious (1 × 10^−4^ ≤ *s*_*d*_ < 1 × 10^−3^), moderately deleterious (1 × 10^−3^ ≤ *s*_*d*_ < 1 × 10^−2^) and strongly deleterious (1 × 10^−2^ ≤ *s*_*d*_ <∞) classes. Recombination rates were varied between 0.5 × mean and 0.001 × mean, where the mean recombination rate was 3.12 × 10^−8^ per site/generations. The black bars represent the simulated distribution of *s*_*d*_. The error bars denote the standard deviation of proportions estimated from 5 independent replicates.

**Supplementary Figure 8:**
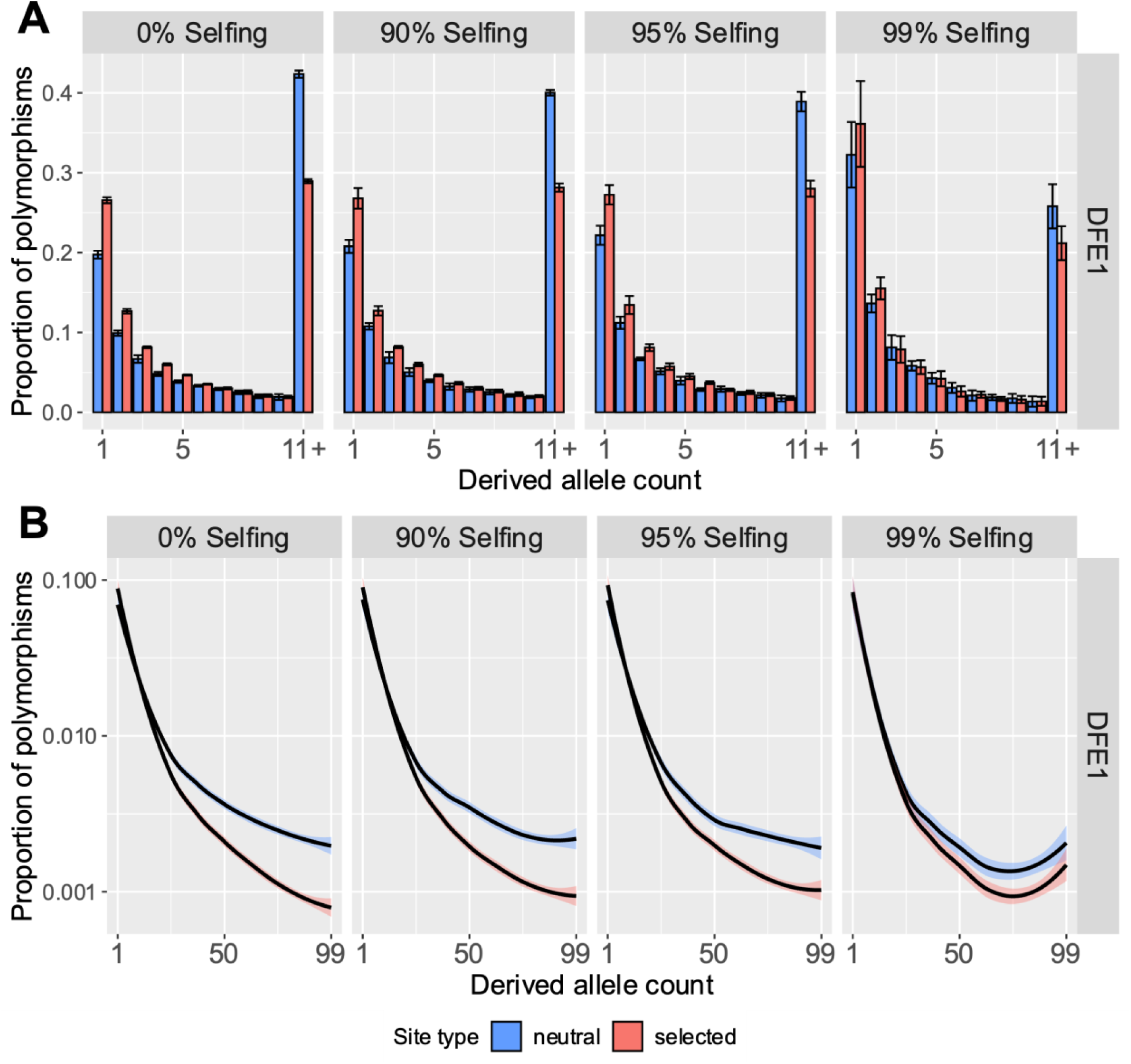
Site frequency spectra of neutral and selected alleles from 100 genomes sampled from simulations where all selected mutations were deleterious, for the weakly deleterious DFE. For **(A)** the y-axis represents the proportion of segregating polymorphisms that fall into the given derived allele count. The last class (11+) refers to the derived allele counts 11-99. The error bars denote the standard deviation of proportions estimated from 5 independent replicates. For **(B)** the y-axis is plotted on a log scale, with all derived allele counts displayed. Lines were smoothed with LOESS.

**Supplementary Figure 9:**
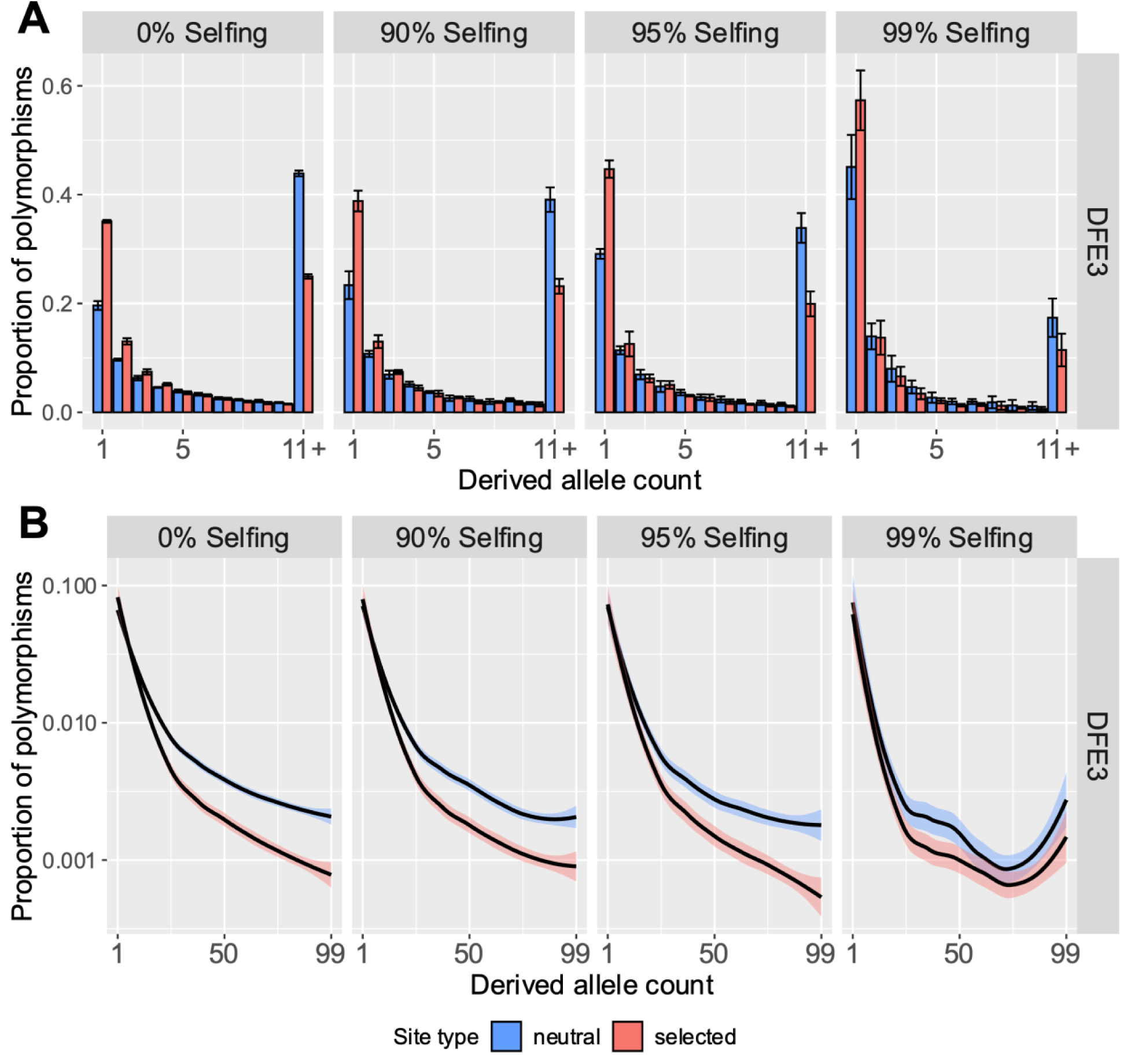
Site frequency spectra of neutral and selected alleles from 100 genomes sampled from simulations where all selected mutations were deleterious, for the strongly deleterious DFE. For **(A)** the y-axis represents the proportion of segregating polymorphisms that fall into the given derived allele count. The last class (11+) refers to the derived allele counts 11-99. The error bars denote the standard deviation of proportions estimated from 5 independent replicates. For **(B)** the y-axis is plotted on a log scale, with all derived allele counts displayed. Lines were smoothed with LOESS.

**Supplementary Figure 10:**
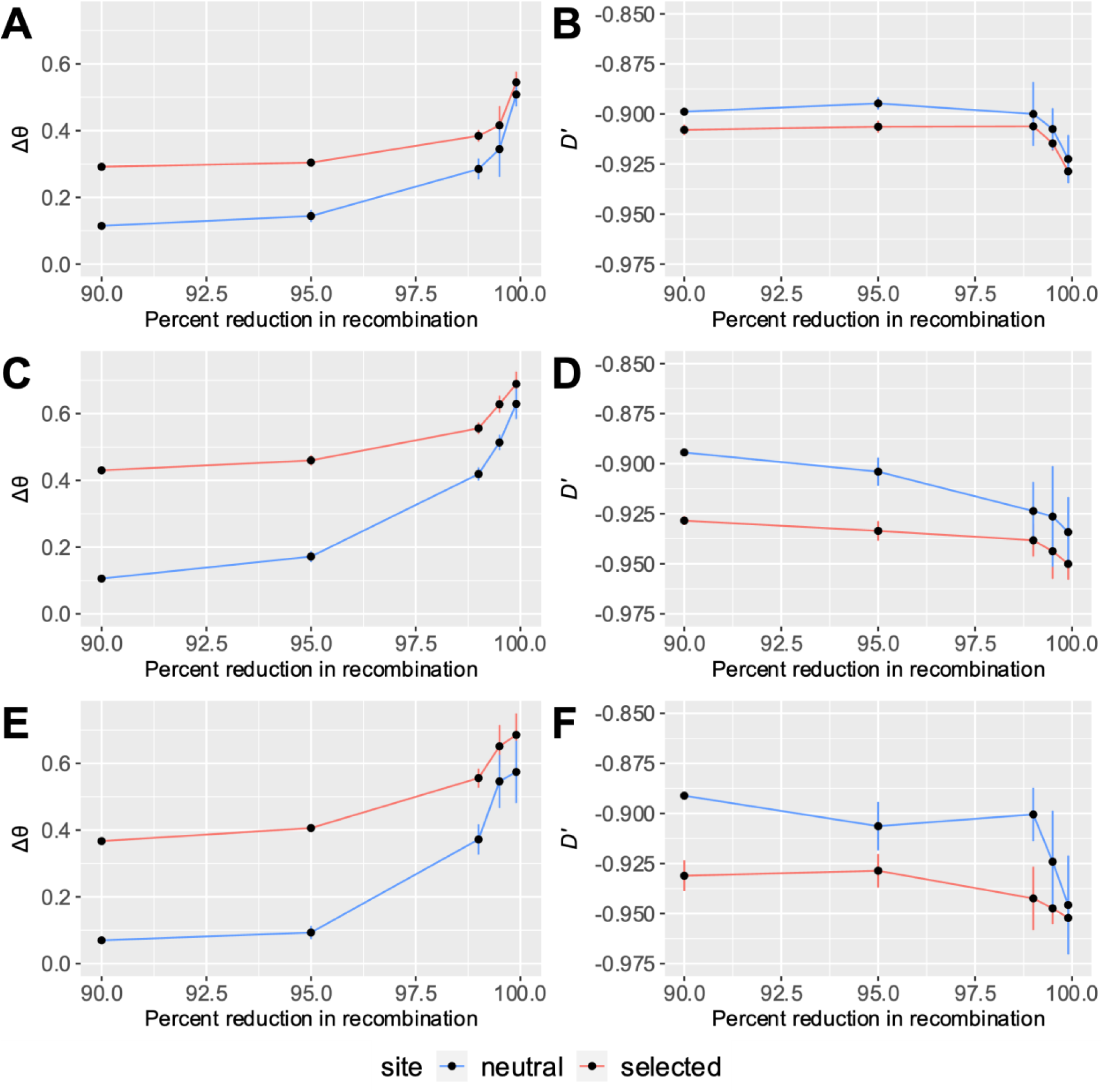
Effects of low recombination on the SFS and linkage disequilibrium (LD) at neutral (red lines) and selected (blue lines) sites. The skew in the SFS was quantified using Δ*θ* for DFE1 (**A**), DFE2 (**C**), and DFE3 (**E**), calculated using alleles at all frequencies, while LD was summarized by the statistic *D*′ for DFE1 (**B**), DFE2 (**D**), and DFE3 (**F**), calculated for alleles at frequencies of 1%-5%. The error bars denote the standard deviation of the genome-wide average of each summary statistic estimated from 5 independent replicates.

**Supplementary Figure 11:**
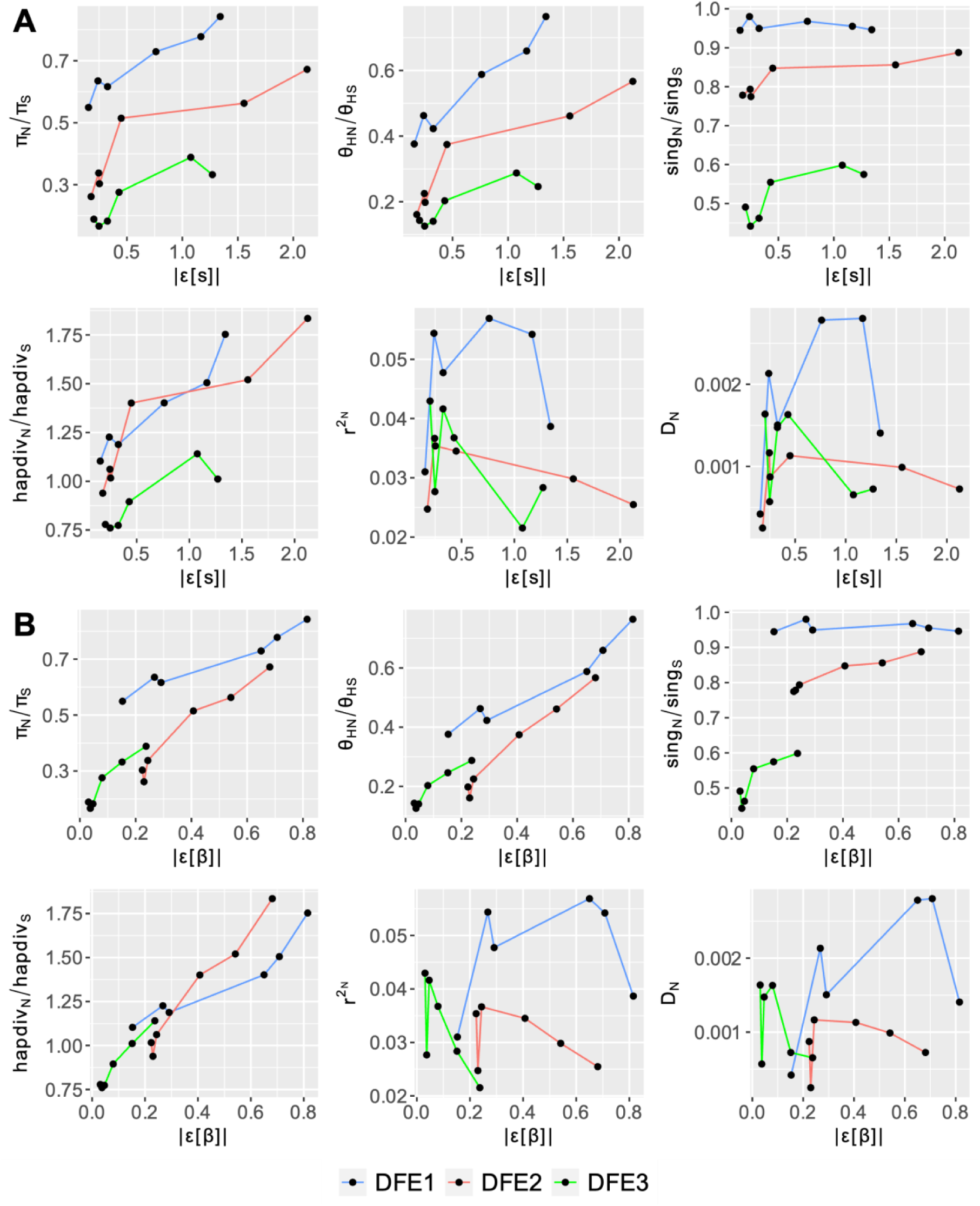
Correlation of error in inference of the parameters 2*Ns* (**A**) and *β* (**B**) by DFE-alpha with various summary statistics calculated by pylibseq. Data points represent the average error and statistic value averaged over five replicates. The skew in the SFS was quantified using *π* (per site), *θ*_*H*_ (per site), and singleton density (*sing*), while LD was summarized by the statistics haplotype diversity (*hapdiv*), *D*, and *r*^2^ using minor alleles across the window. Most statistics were calculated in 5000 bp windows for all allele frequencies, and values for nonsynonymous sites were divided by synonymous sites. However, *D* and *r*^2^ were calculated for only nonsynonymous alleles at frequencies of 1%-5%.

**Supplementary Figure 12:**
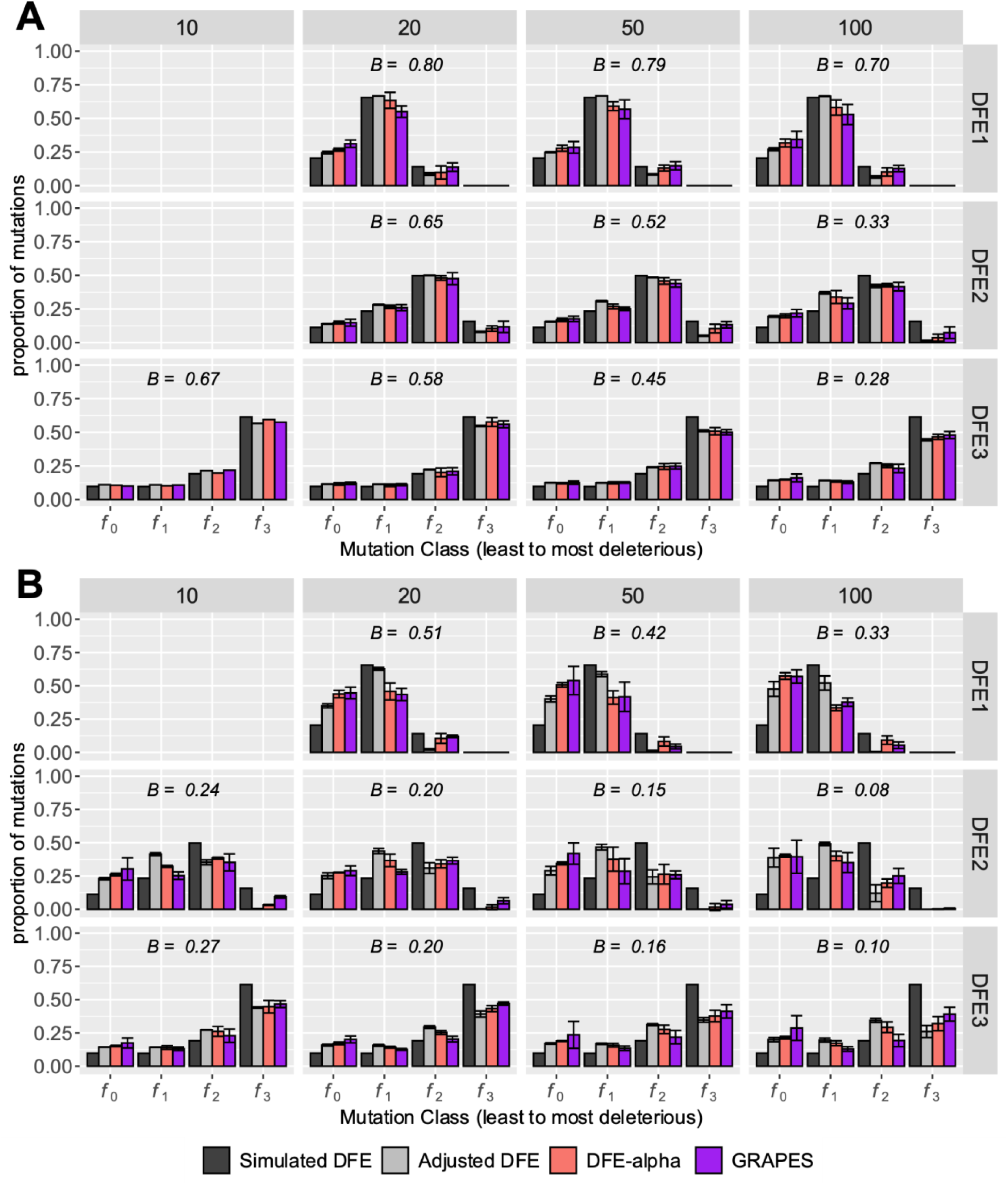
Effects of varying the rescaling factor on the estimated DFE of new deleterious mutations in partially selfing organisms using DFE-alpha and GRAPES at selfing rates of (**A**) 95% and (**B**) 99%. The rescaling factors were reduced from the original 100 to 50, 20, and 10 (denoted on top of the columns). The error bars denote the standard deviation of proportions estimated from 5 independent replicates, except for the rescaling factor of 10, where only one replicate of DFE3 at 95% selfing and three replicates each for DFE2 and DFE3 at 99% selfing were completed.

**Supplementary Figure 13:**
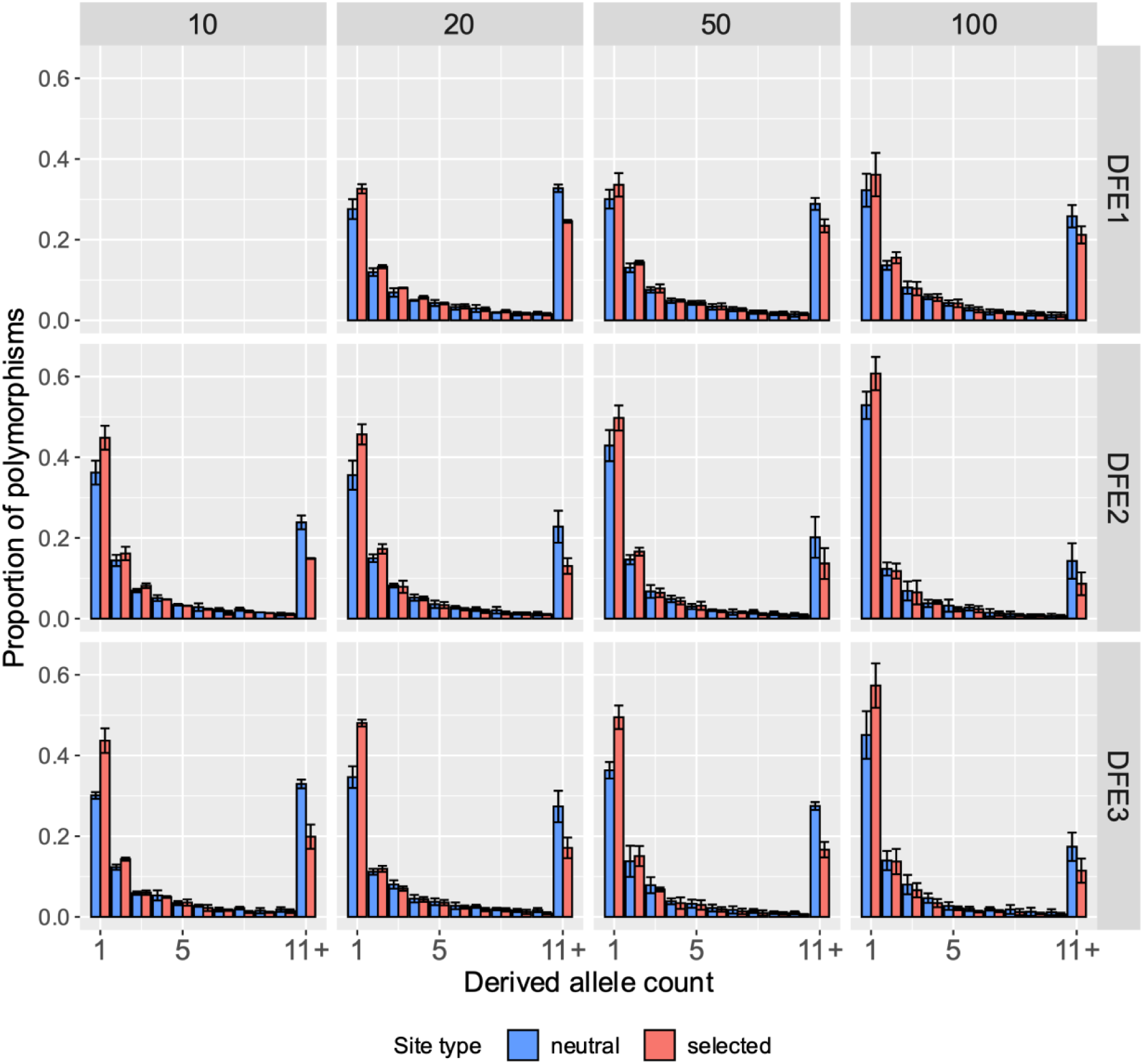
Site frequency spectra of neutral and selected alleles from 100 genomes sampled from simulations with 99% selfing where all selected mutations were deleterious, with varying rescaling factors plotted on the x-axis. The y-axis represents the proportion of segregating polymorphisms that fall into the given derived allele count. The last class (11+) refers to the derived allele counts 11-99. The error bars denote the standard deviation of proportions estimated from 5 independent replicates, except for the rescaling factor of 10, where only three replicates each for DFE2 and DFE3 were completed.

**Supplementary Figure 14:**
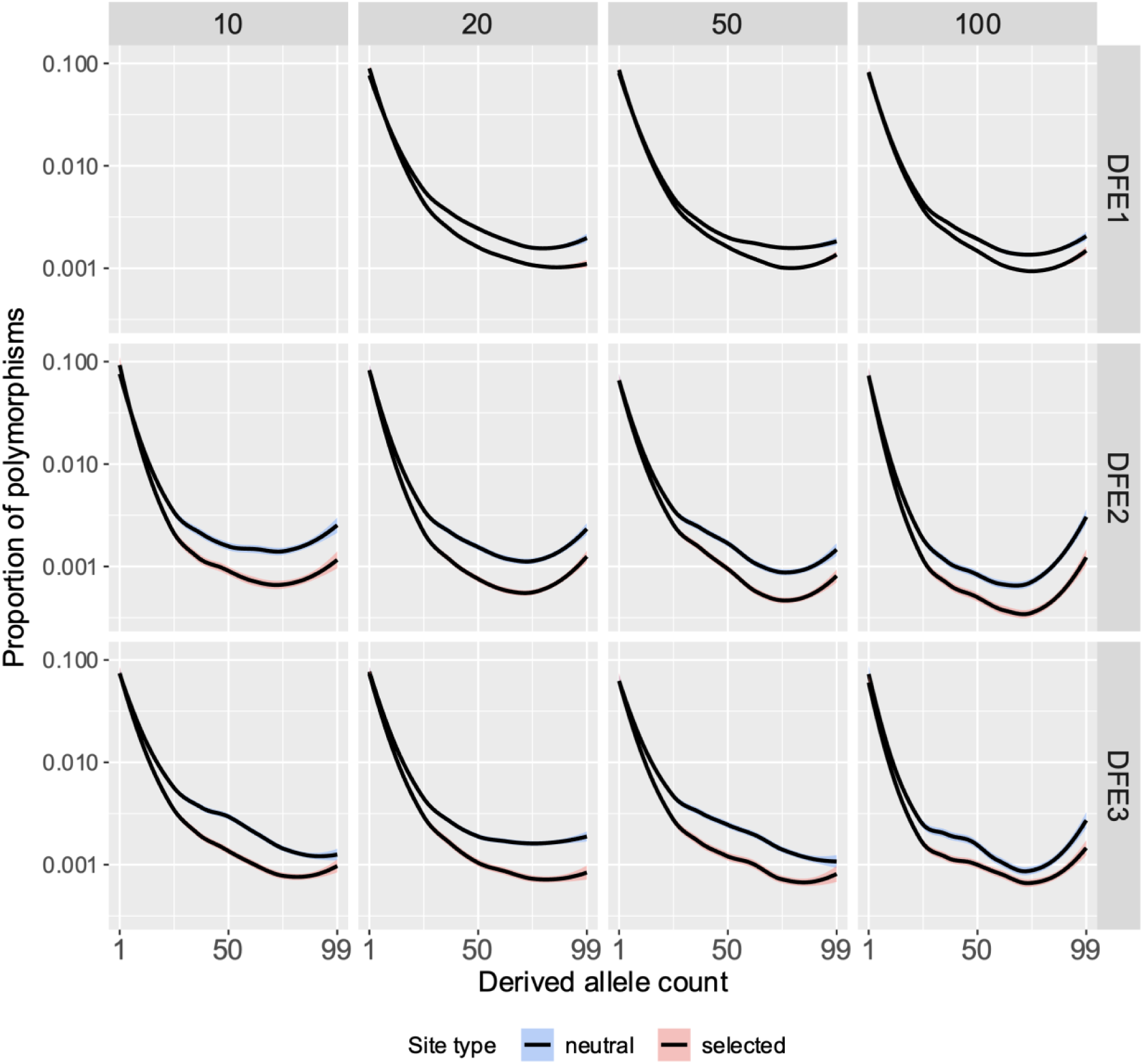
Site frequency spectra of neutral and selected alleles from 100 genomes sampled from simulations where all selected mutations were deleterious, with varying rescaling factors plotted on the x-axis. The y-axis is plotted on a log scale, with all derived allele counts (frequencies 1-99) displayed. Lines were smoothed with LOESS.

**Supplementary Figure 15:**
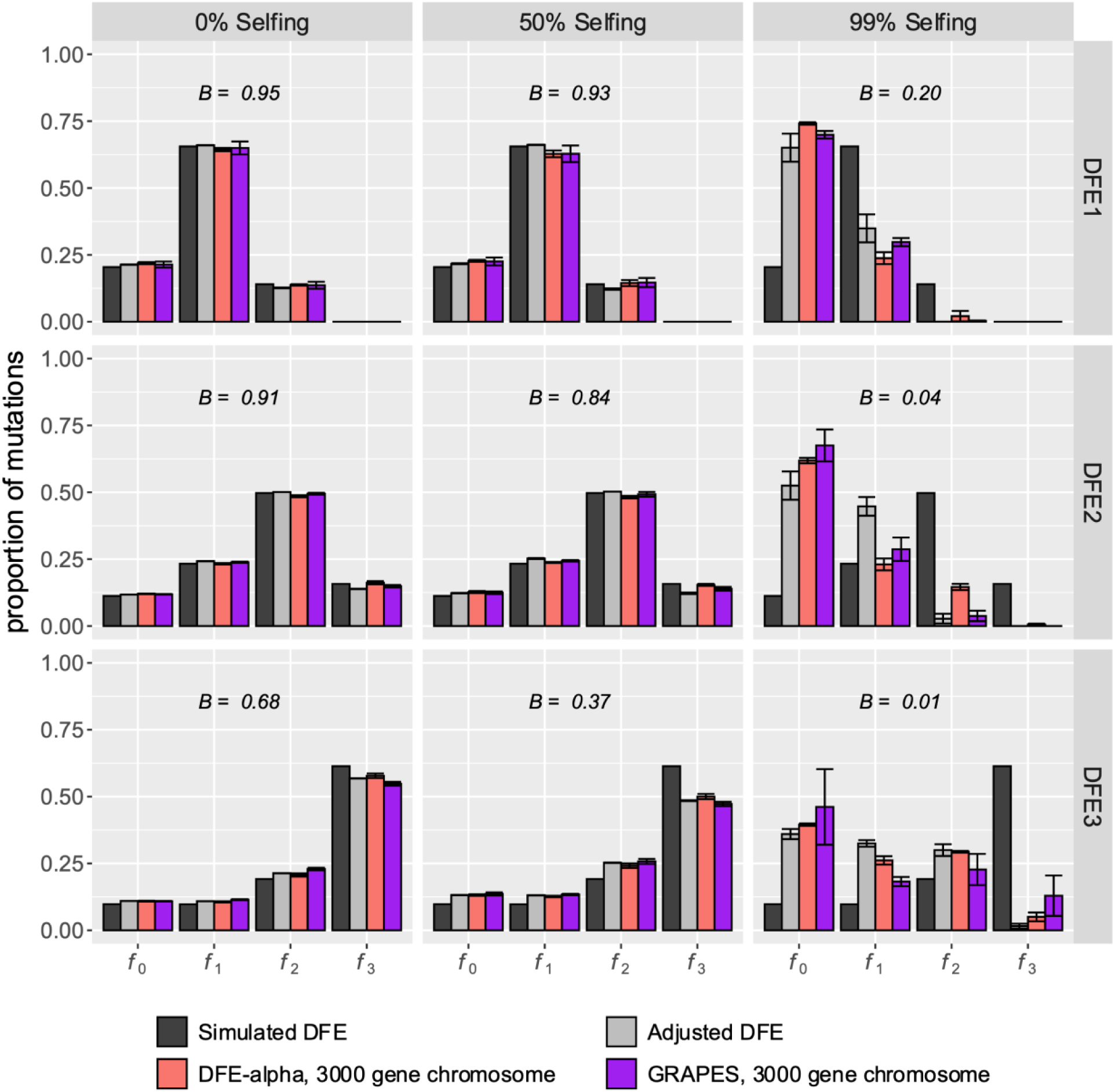
The inference of the deleterious DFE when varying chromosome sizes were simulated: a chromosome with 500 genes (shown in darker shades) vs a chromosome with 3000 genes (shown in lighter shades). Results are shown in terms of the proportion of mutations in effectively neutral (*f*_0_), weakly deleterious (*f*_1_), moderately deleterious (*f*_2_) and strongly deleterious (*f*_3_) classes. The nucleotide site diversity with background selection (*B*) relative to its expectation under strict neutrality is shown in each panel. The adjusted DFE represents the distribution of 2*NBs*_*d*_. Inference was performed using DFE-alpha (red bars) and GRAPES purple bars). The error bars denote the standard deviation of proportions estimated from 5 independent replicates.

**Supplementary Figure 16:**
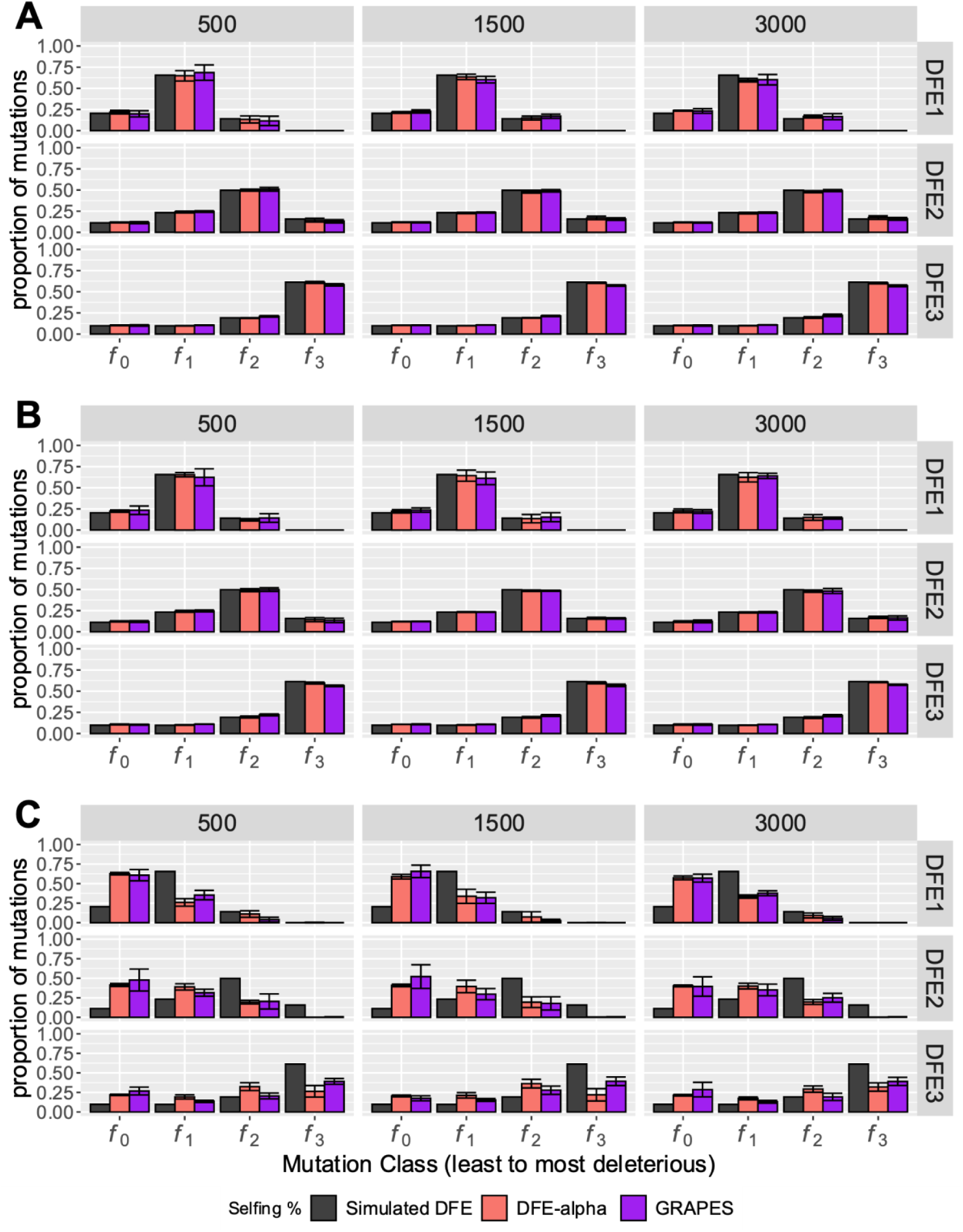
Effects of varying the coding density on the estimated DFE of new deleterious mutations in partially selfing organisms using DFE-alpha and GRAPES at selfing rates of (**A**) 0%, (**B**) 50%, and (**C**) 99%. The intergenic regions were reduced from the original 3000 bp to 1500 bp and 500 bp (denoted on top of the columns). The error bars denote the standard deviation of proportions estimated from 5 independent replicates.

**Supplementary Figure 17:**
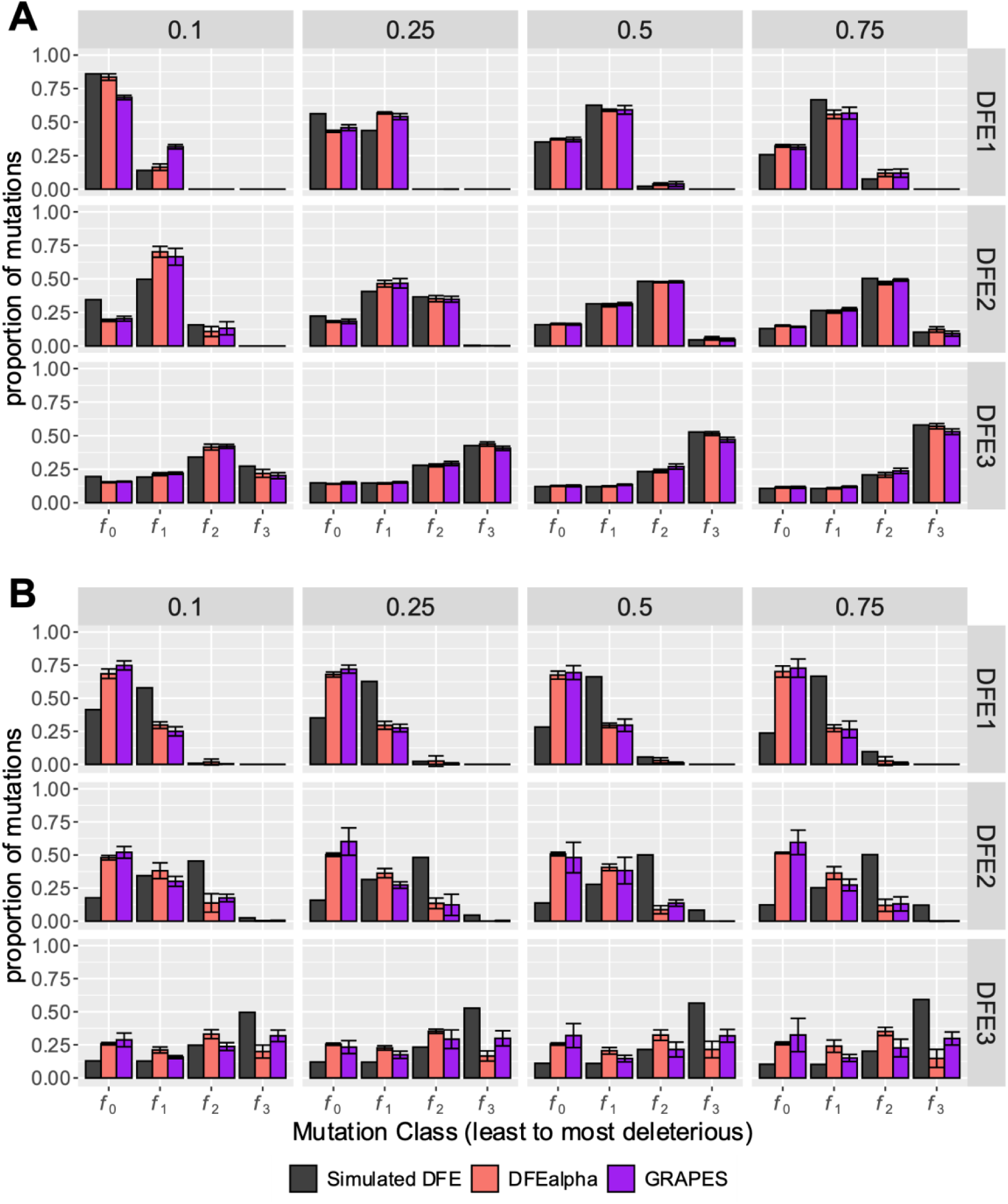
Effects of selfing on DFE inference of new deleterious mutations with varying dominance coefficients using DFE-alpha (red bars) and GRAPES (purple bars). The dominance of selected mutations is noted at the top of each column. Results are shown for populations with (**A)** 0% selfing and (**B)** 99% selfing. Here the effectively neutral (*f*_0_), weakly deleterious (*f*_1_), moderately deleterious (*f*_2_) and strongly deleterious (*f*_3_) classes represent 0 < 2*N*_*self*_ *s*_*d*_*h*_*self*_ < 1, 1 < 2*N*_*self*_ *s*_*d*_*h*_*self*_ < 10, 10 < 2*N*_*self*_ *s*_*d*_*h*_*self*_ < 100, and 100 < 2*N*_*self*_ *s*_*d*_*h*_*self*_ < 2*N*_*self*_, respectively. The error bars denote the standard deviation of proportions estimated from 5 independent replicates.

**Supplementary Figure 18:**
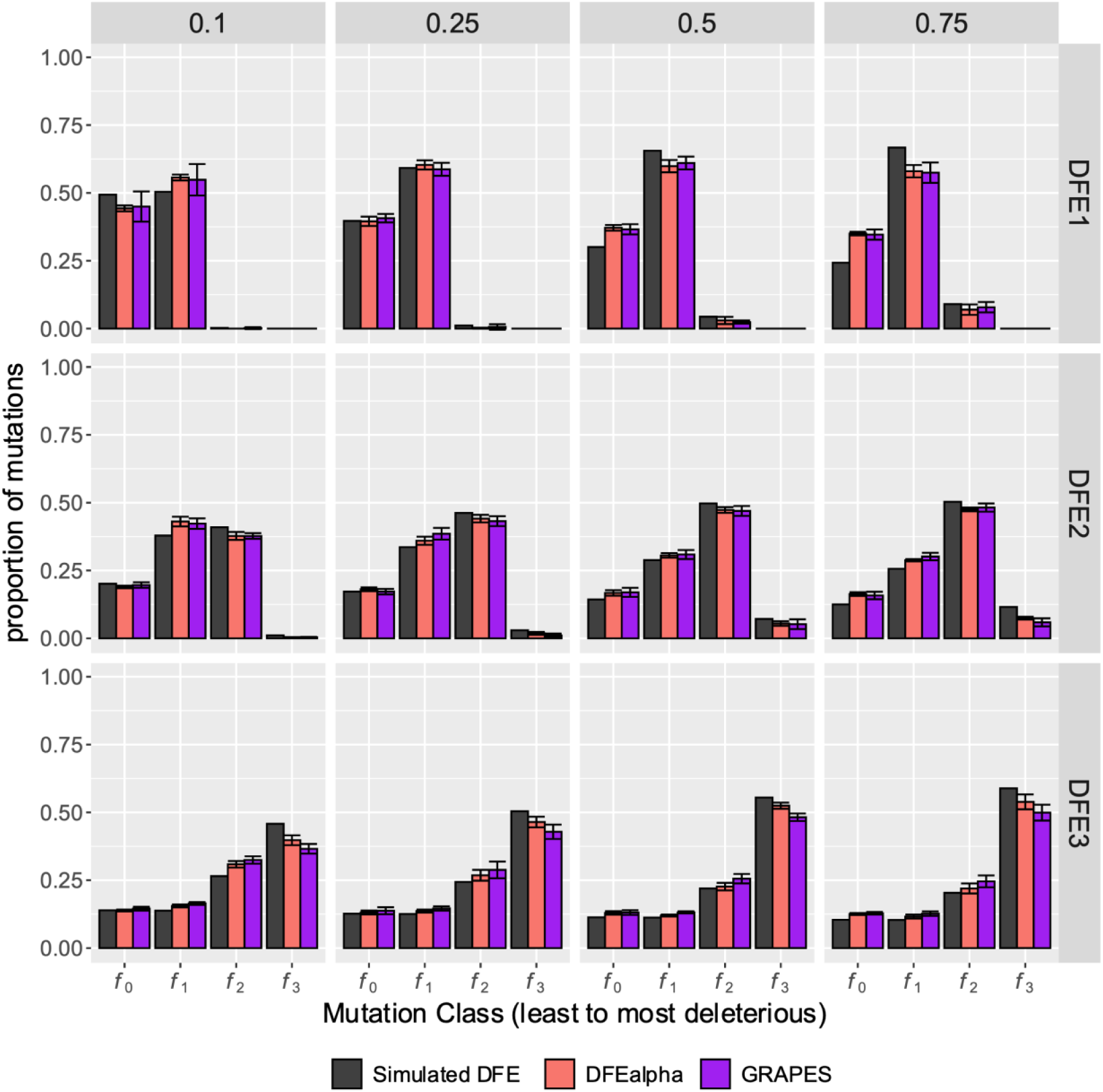
Effects of selfing (50%) on the inference of the DFE of new deleterious mutations with varying dominance coefficients using DFE-alpha and GRAPES. The dominance of all new mutations is noted at the top of each column. The error bars denote the standard deviation of proportions estimated from 5 independent replicates.

**Supplementary Figure 19:**
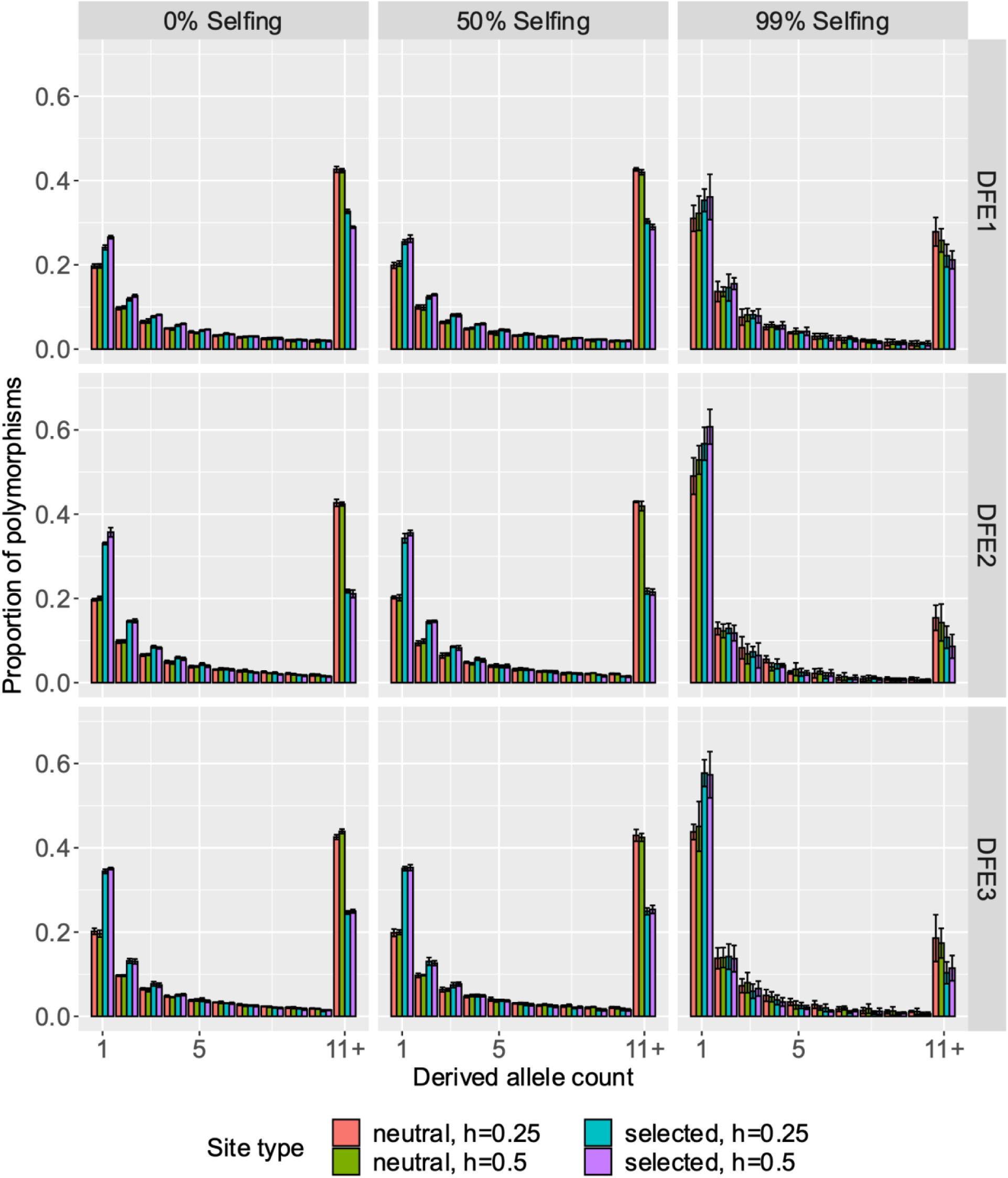
Site frequency spectra of neutral and selected alleles for varying rates of selfing, dominance coefficients, and deleterious DFEs. 100 genomes were sampled from simulations. Selected mutations had a dominance coefficient *h* = 0.25 or 0.5.The y-axis represents the proportion of segregating polymorphisms that fall into the given allele-frequency class. The last class (11+) refers to the derived allele counts 11-100.

**Supplementary Figure 20:**
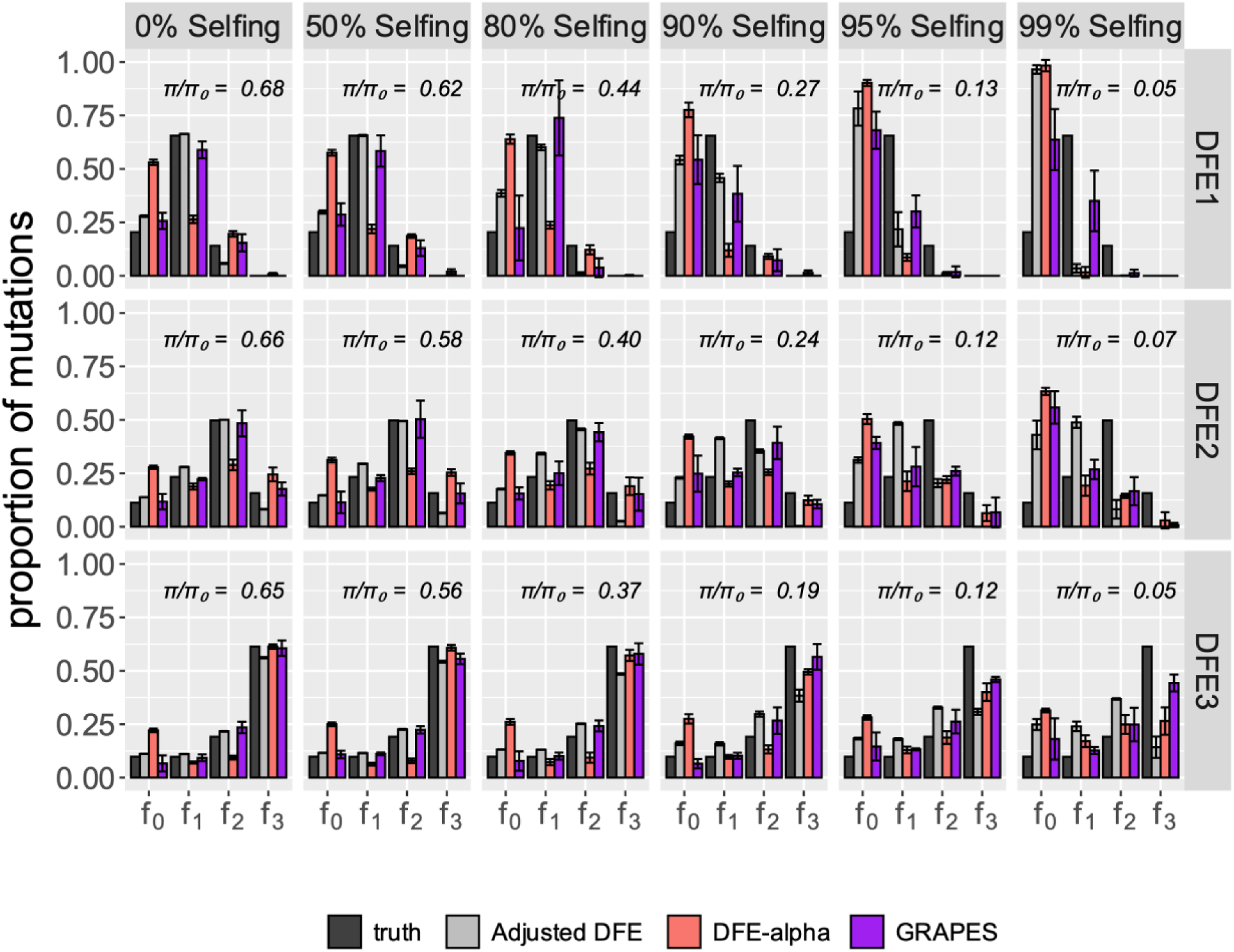
Effects of selfing on the inference of the DFE of new deleterious mutations using DFE-alpha and GRAPES (shown in purple bars). Results are shown in terms of the proportion of mutations in effectively neutral (*f*_0_), weakly deleterious (*f*_1_), moderately deleterious (*f*_2_) and strongly deleterious (*f*_3_) classes. The adjusted DFE represents the distribution of 2*N*(*π*_*obs*_⁄*π*_*exp*_)*s*_*d*_, where *π*_*obs*_ is the neutral nucleotide site diversity observed in each scenario and *π*_*exp*_ is the neutral nucleotide site diversity expected in a population of size *N* at strict neutrality. 0.1% of new exonic mutations were beneficial and exponentially distributed with a mean 2*Ns*_*a*_ = 200. Here, the unfolded SFS was used for inference. The error bars denote the standard deviation of proportions estimated from 5 independent replicates.

**Supplementary Figure 21:**
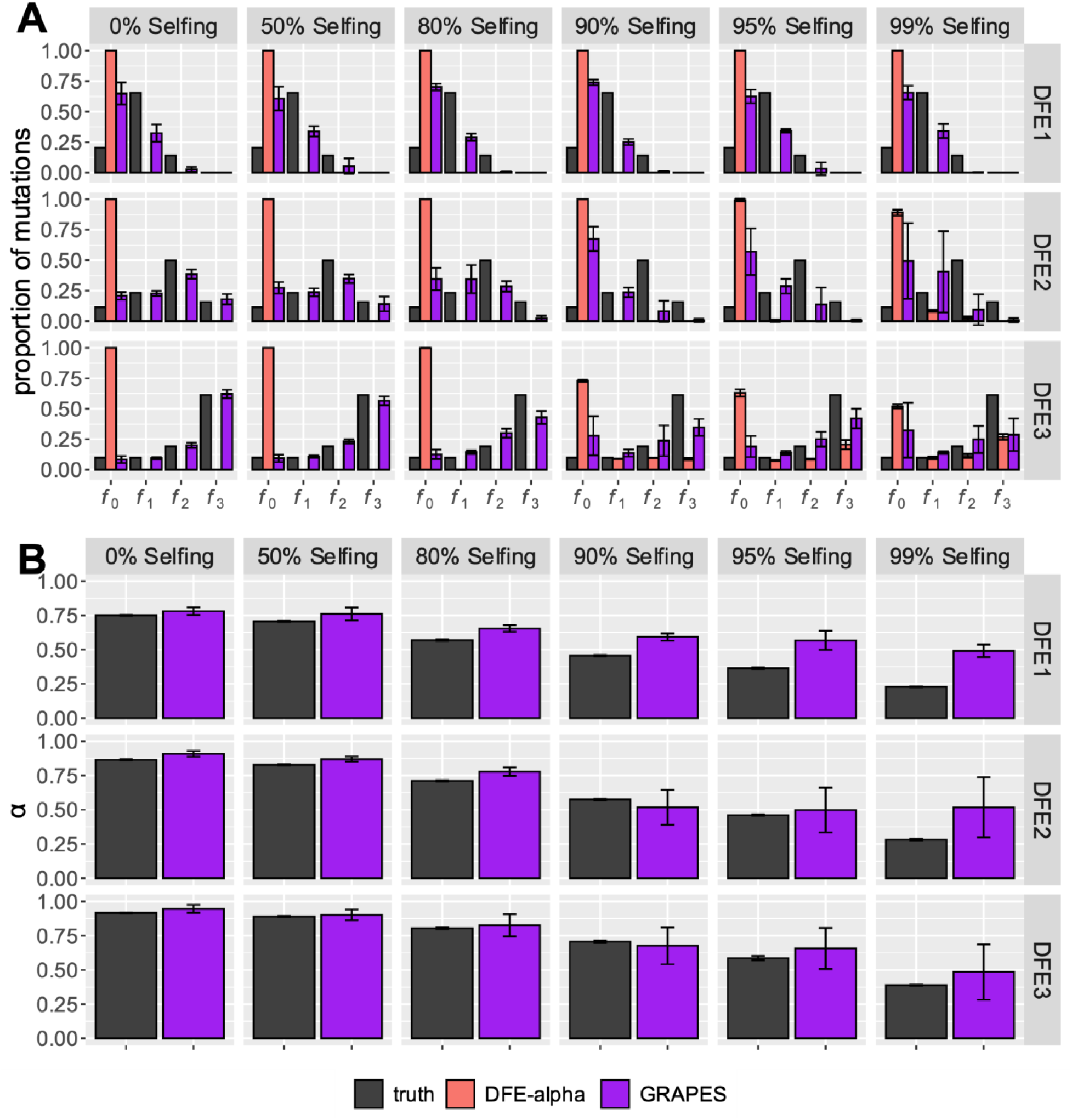
Effects of selfing on the inference of (A) the DFE of new deleterious mutations and (B) the proportion of beneficial fixations (α) using DFE-alpha and GRAPES when beneficial mutations are common (1%). Results for A are shown in terms of the proportion of mutations in effectively neutral (*f*_0_), weakly deleterious (*f*_1_), moderately deleterious (*f*_2_) and strongly deleterious (*f*_3_) classes. Here, 1.0% of new exonic mutations were beneficial and exponentially distributed with a mean 2*Ns*_*a*_ = 200. Here, the unfolded SFS was used for inference. The error bars denote the standard deviation of proportions estimated from 5 independent replicates.

**Supplementary Figure 22:**
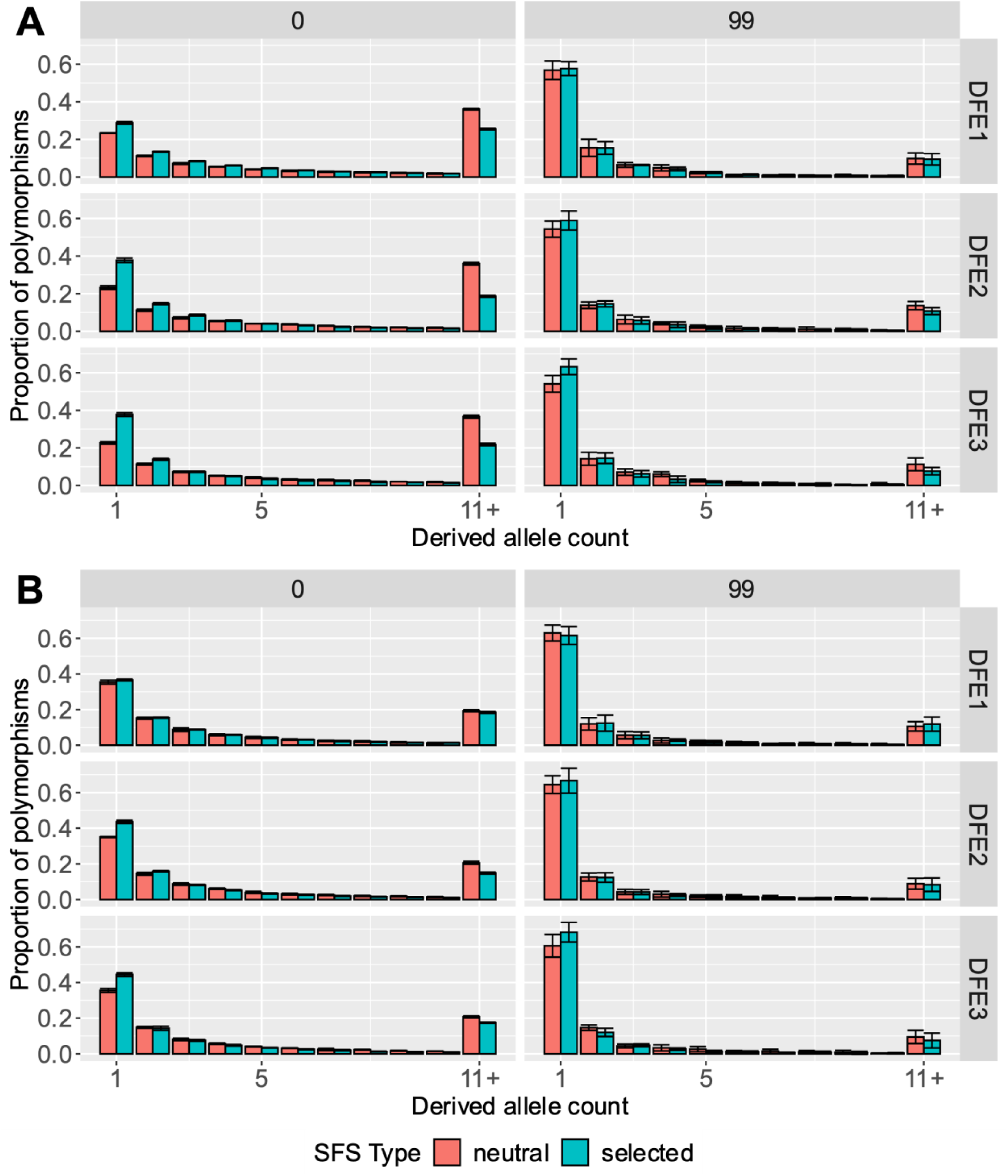
Site frequency spectra of neutral vs selected sites when a full DFE was simulated (*i.e*., both deleterious and beneficial mutations were present). 100 genomes were sampled and the y-axis depicts the proportion of segregating polymorphisms that fall into the given allele-count class. Panel **A** shows simulations where 0.1% of new mutations at selected sites were beneficial and Panel **B** shows simulations where 1% of new mutations at selected sites were beneficial. The error bars denote the standard deviation of proportions estimated from 5 independent replicates.

**Supplementary Figure 23:**
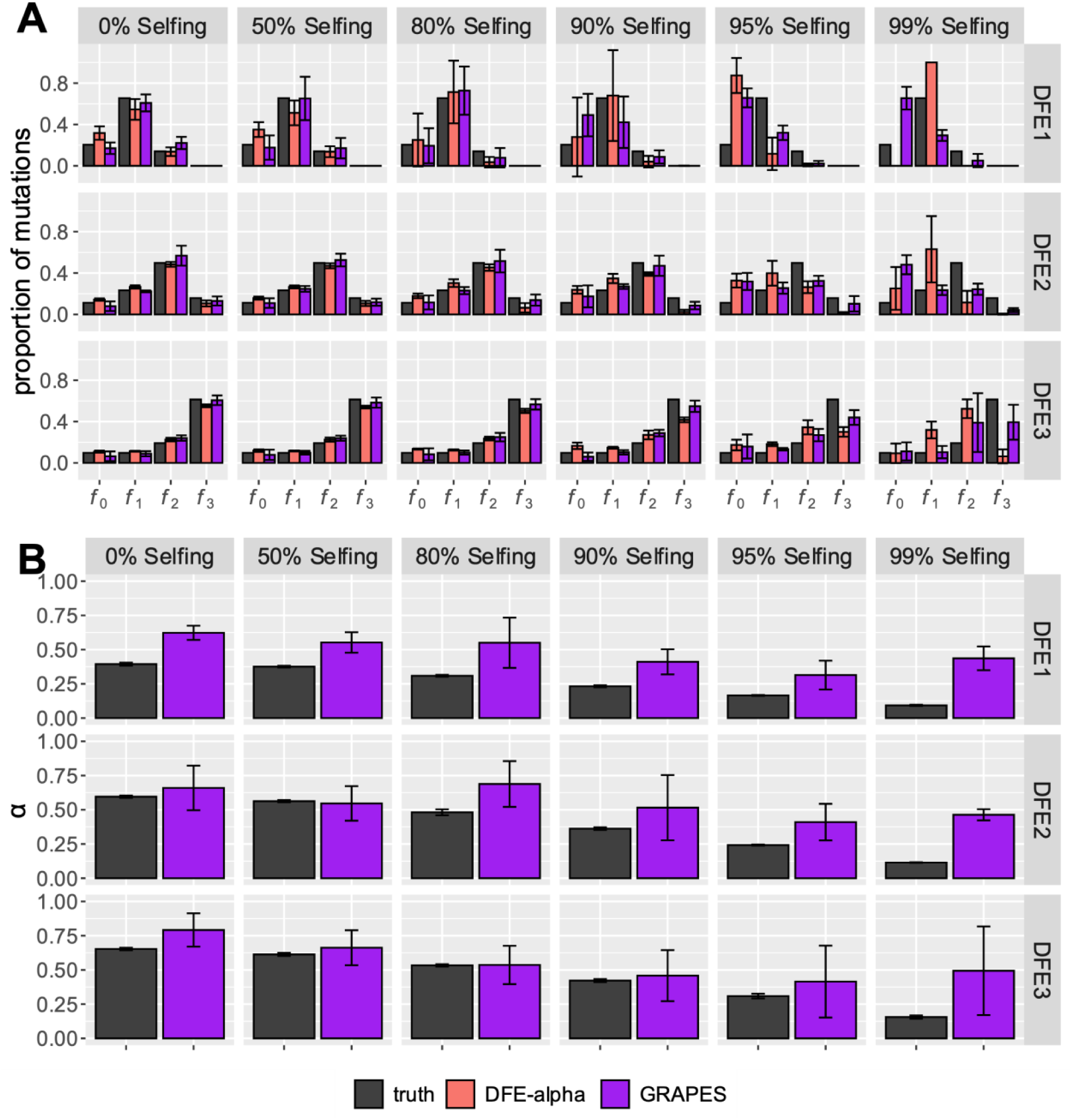
Effects of selfing on the inference of (A) the DFE of new deleterious mutations and (B) the proportion of beneficial fixations (α) using DFE-alpha and GRAPES, when beneficial mutations were rare. Results for A are shown in terms of the proportion of mutations in effectively neutral (*f*_0_), weakly deleterious (*f*_1_), moderately deleterious (*f*_2_) and strongly deleterious (*f*_3_) classes. Here, 0.1% of new exonic mutations were beneficial and exponentially distributed with a mean 2*Ns*_*a*_=200. Here, the folded SFS was used for inference. The error bars denote the standard deviation of proportions estimated from 5 independent replicates.

**Supplementary Figure 24:**
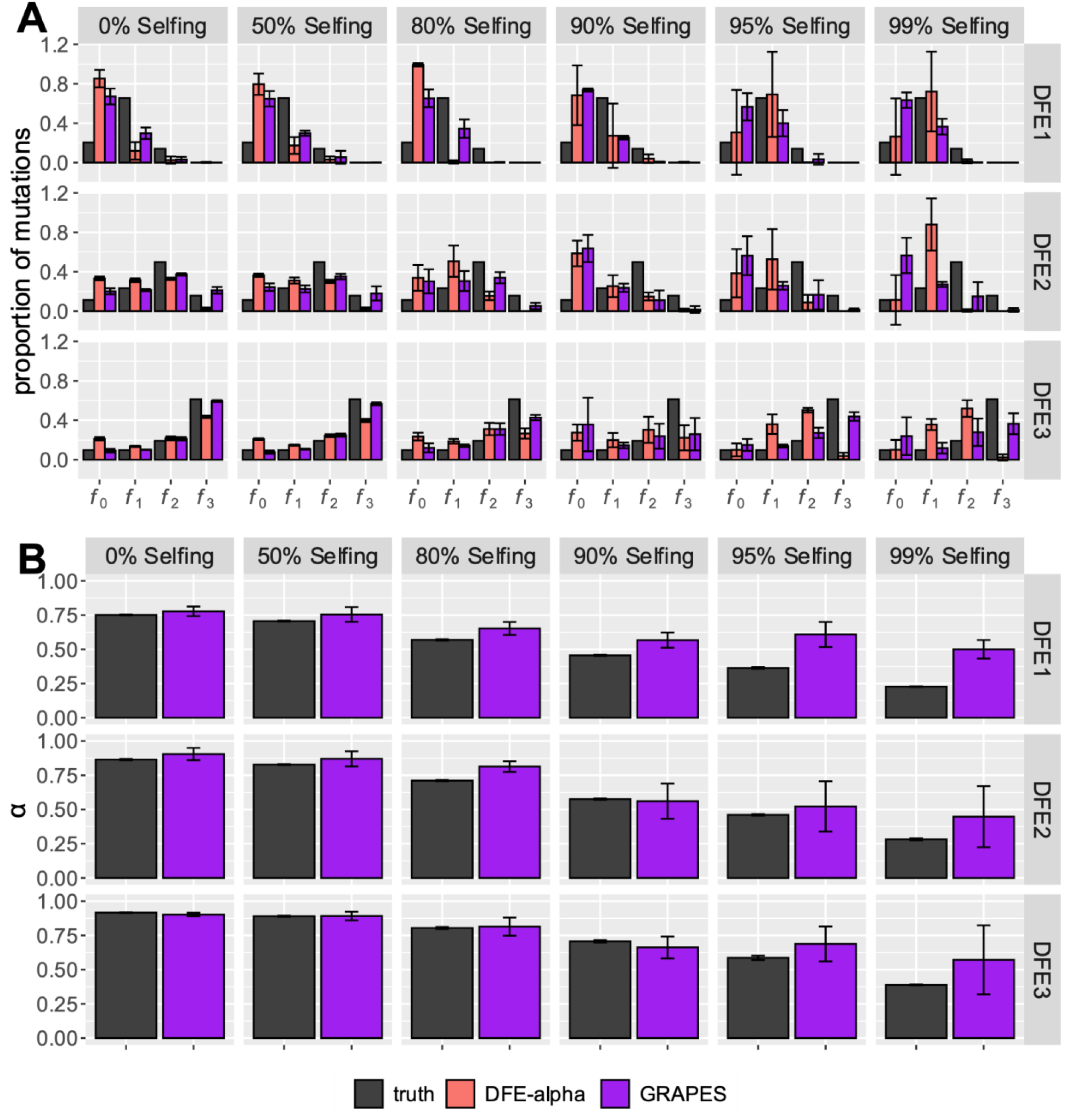
Effects of selfing on the inference of (A) the DFE of new deleterious mutations and (B) the proportion of beneficial fixations (α) using DFE-alpha and GRAPES, when beneficial mutations were common. Results for A are shown in terms of the proportion of mutations in effectively neutral (*f*_0_), weakly deleterious (*f*_1_), moderately deleterious (*f*_2_) and strongly deleterious (*f*_3_) classes. Here, 1.0% of new exonic mutations were beneficial and exponentially distributed with a mean 2*Ns*_*a*_=200. Here, the folded SFS was used for inference. The error bars denote the standard deviation of proportions estimated from 5 independent replicates.

**Supplementary Figure 25:**
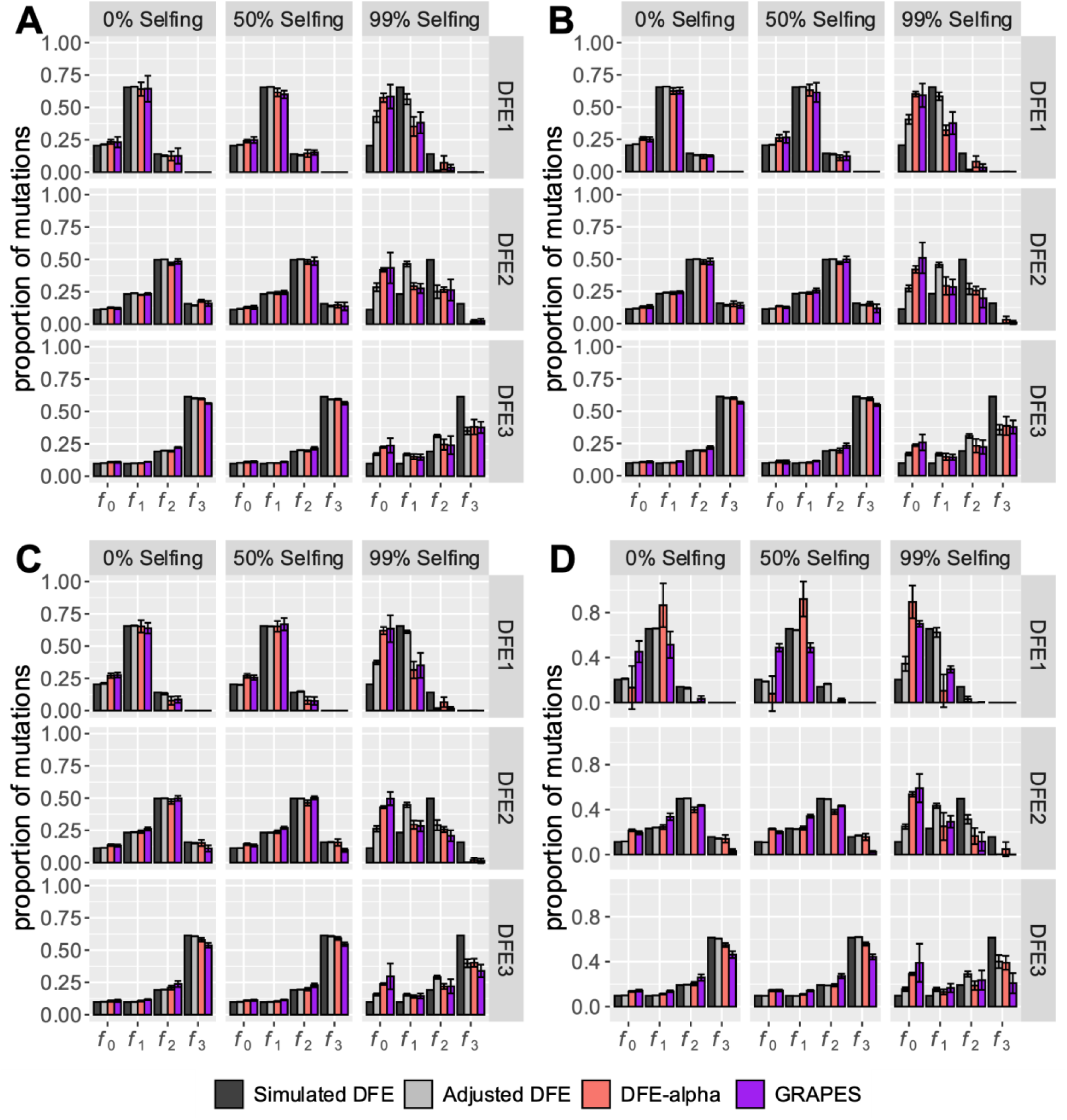
Effects of population structure and selfing on the inference of the DFE of new deleterious mutations using DFE-alpha and GRAPES. Populations were simulated under an island model with five demes, and genomes were sampled evenly from each deme. Here the metapopulation effective size was equal to 5000 at 0% selfing (see methods). Results are shown when *N*_*deme*_ *m* was (**A)** 2 **(B)** 1, (**C)** 0.5, and (**D)** 0.1. Results are shown in terms of the proportion of mutations in effectively neutral (*f*_0_), weakly deleterious (*f*_1_), moderately deleterious (*f*_2_) and strongly deleterious (*f*_3_) classes. The nucleotide site diversity with background selection (*B*) relative to its expectation under strict neutrality is shown in each panel. The adjusted DFE represents the distribution of 2*NBs*_*d*_. The error bars denote the standard deviation of proportions estimated from 5 independent replicates.

**Supplementary Figure 26:**
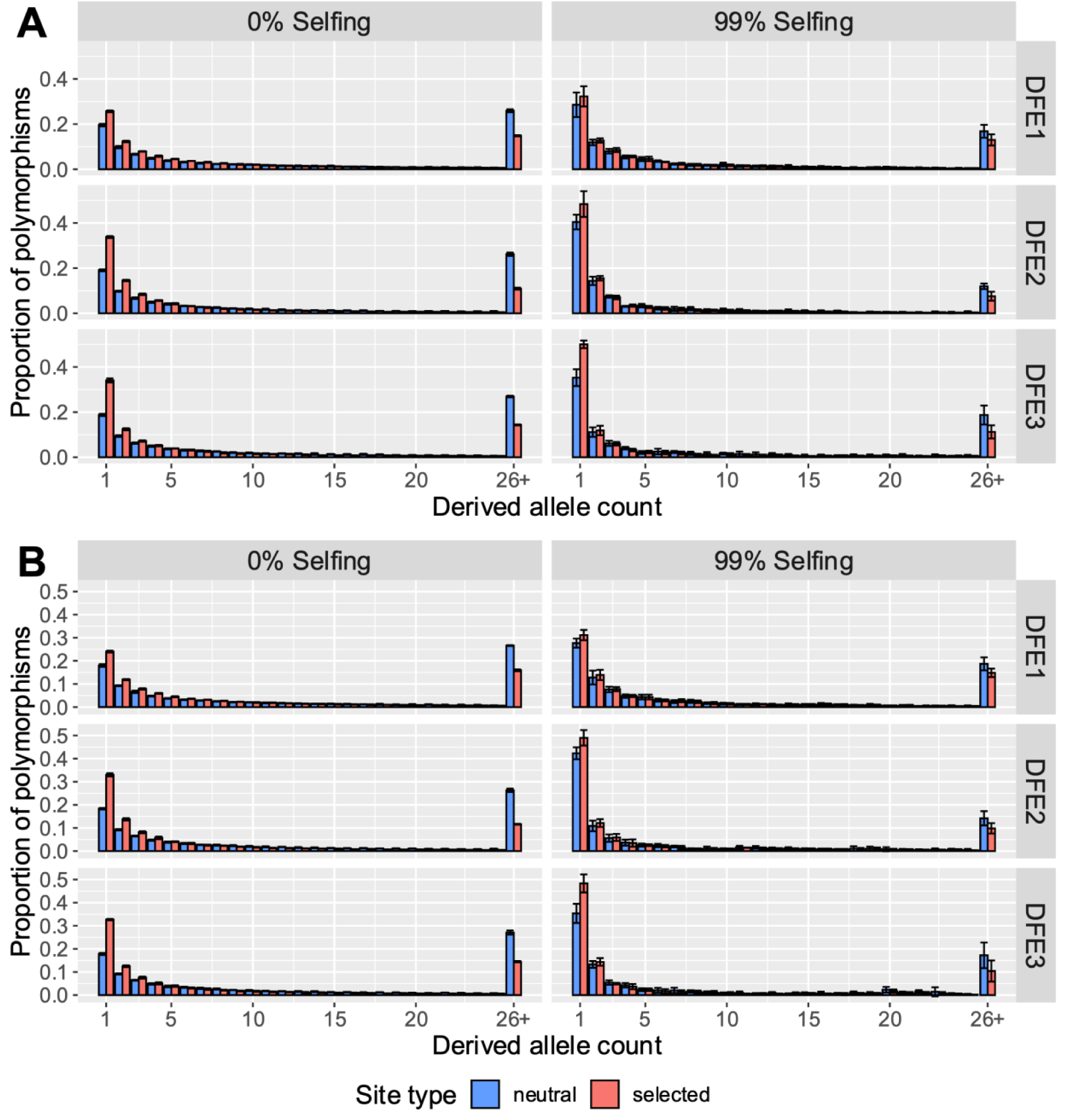
Effects of population structure and selfing on the SFS. Populations were simulated under an island model with five demes, and genomes were sampled evenly from each deme. Here the metapopulation effective size was equal to 5000 at 0% selfing (see methods). Results are shown when *N*_*deme*_ *m* was (**A)** 2 **(B)** 1. The error bars denote the standard deviation of proportions estimated from 5 independent replicates. The last class (26+) refers to the derived allele counts 26-100.

**Supplementary Figure 27:**
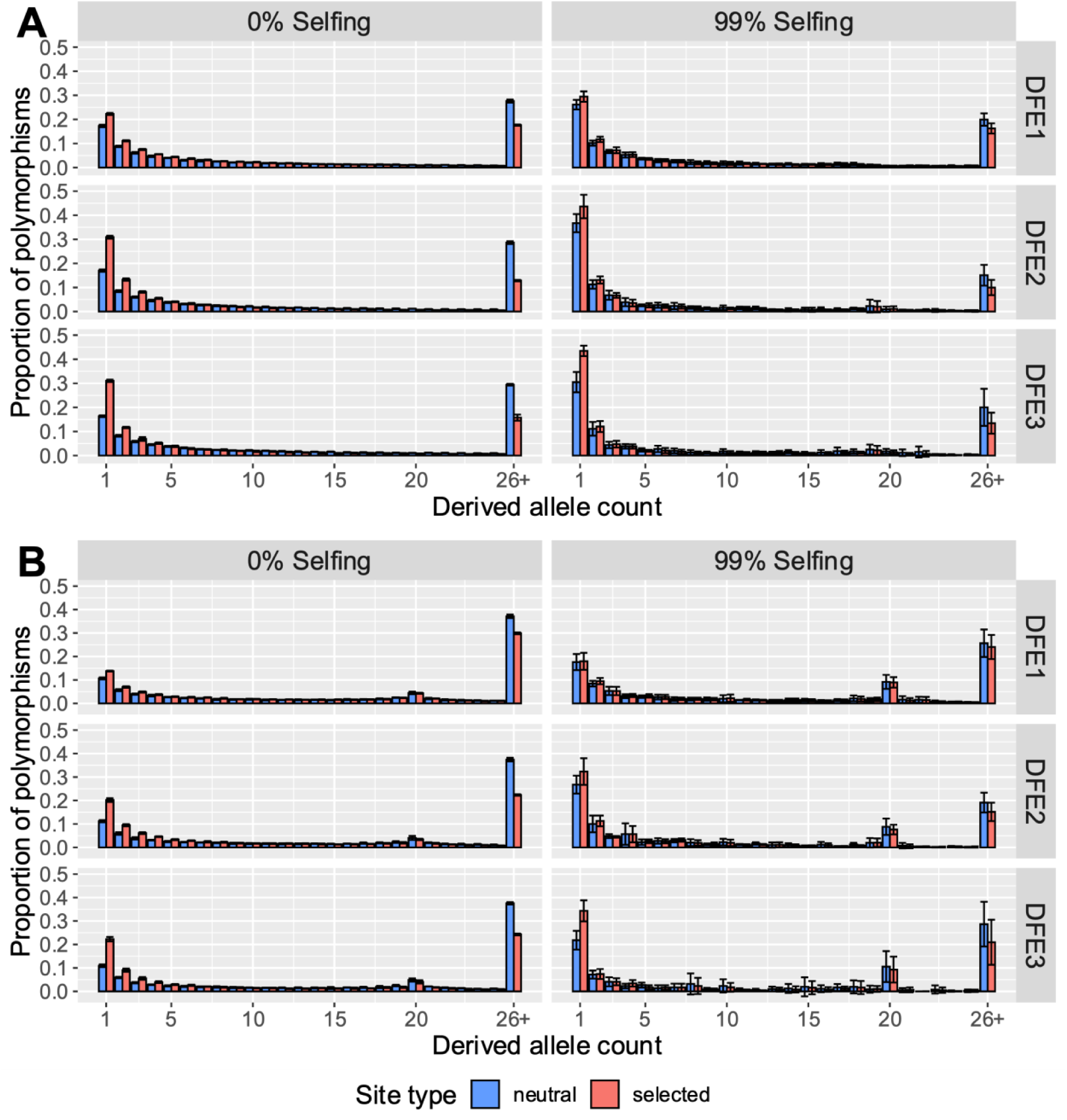
Effects of population structure and selfing on the SFS with even sampling. Populations were simulated under an island model with five demes, and genomes were sampled evenly from each deme. Here the metapopulation effective size was equal to 5000 at 0% selfing (see methods). Results are shown when *N*_*deme*_*m* was (**A)** 0.5 **(B)** 0.1. The error bars denote the standard deviation of proportions estimated from 5 independent replicates. The last class (26+) refers to the derived allele counts 26-100.

**Supplementary Figure 28:**
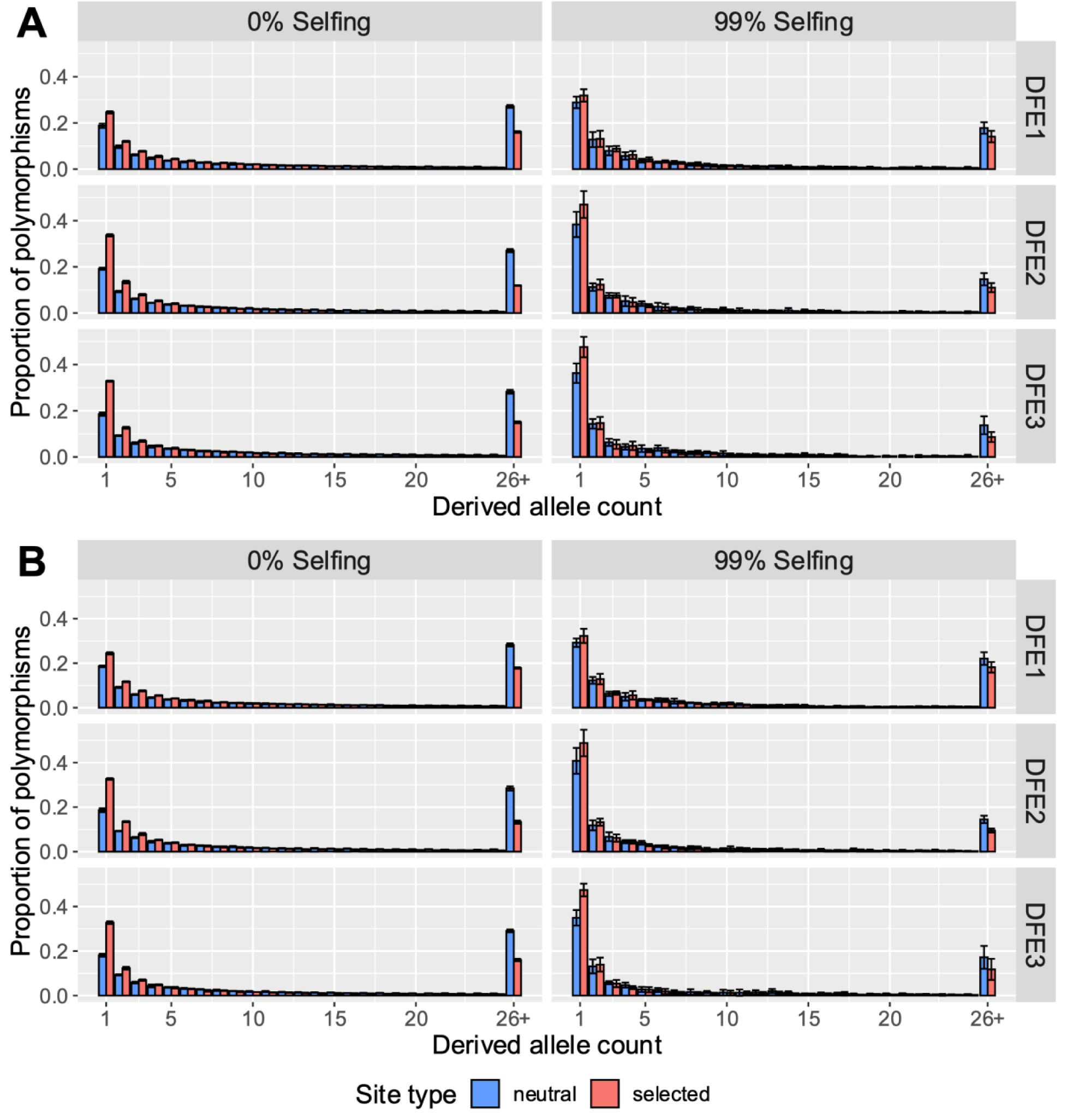
Effects of uneven sampling, population structure, and selfing on the SFS at high migration rates. Populations were simulated under an island model with five demes, and genomes were sampled unevenly (60 genomes from one deme, and 10 from the remaining four demes). Here the metapopulation effective size was equal to 5000 at 0% selfing (see methods). Results are shown when *N*_*deme*_ *m* was (**A)** 2 **(B)** 1. The error bars denote the standard deviation of proportions estimated from 5 independent replicates. The last class (26+) refers to the derived allele counts 26-100.

**Supplementary Figure 29:**
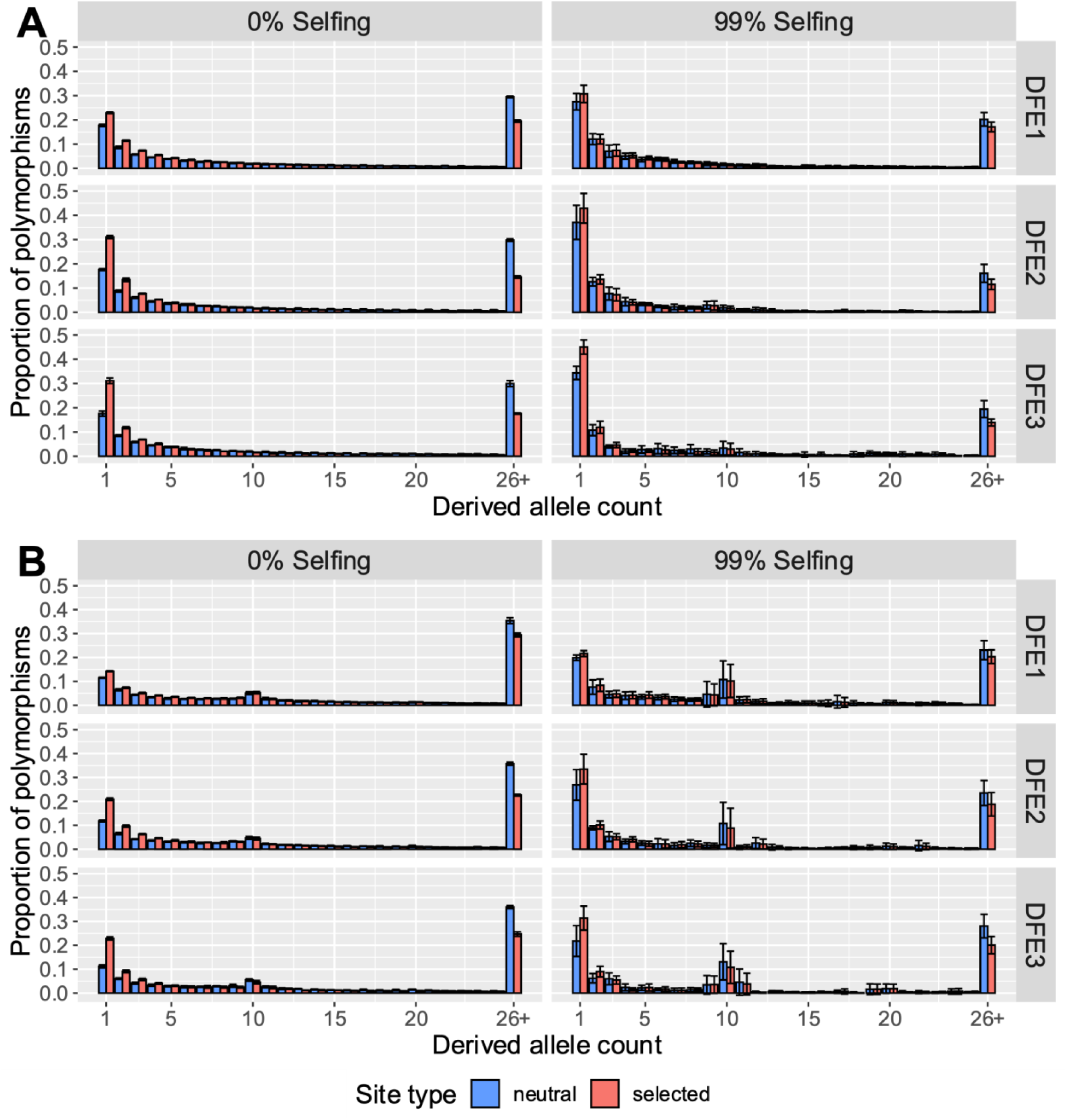
Effects of uneven sampling, population structure, and selfing on the SFS at lower migration rates. Populations were simulated under an island model with five demes, and genomes were sampled unevenly (60 genomes from one deme, and 10 from the remaining four demes). Here the metapopulation effective size was equal to 5000 at 0% selfing (see methods). Results are shown when *N*_*deme*_ *m* was (**A)** 0.5 **(B)** 0.1. The error bars denote the standard deviation of proportions estimated from 5 independent replicates. The last class (26+) refers to the derived allele counts 26-100.

**Supplementary Figure 30:**
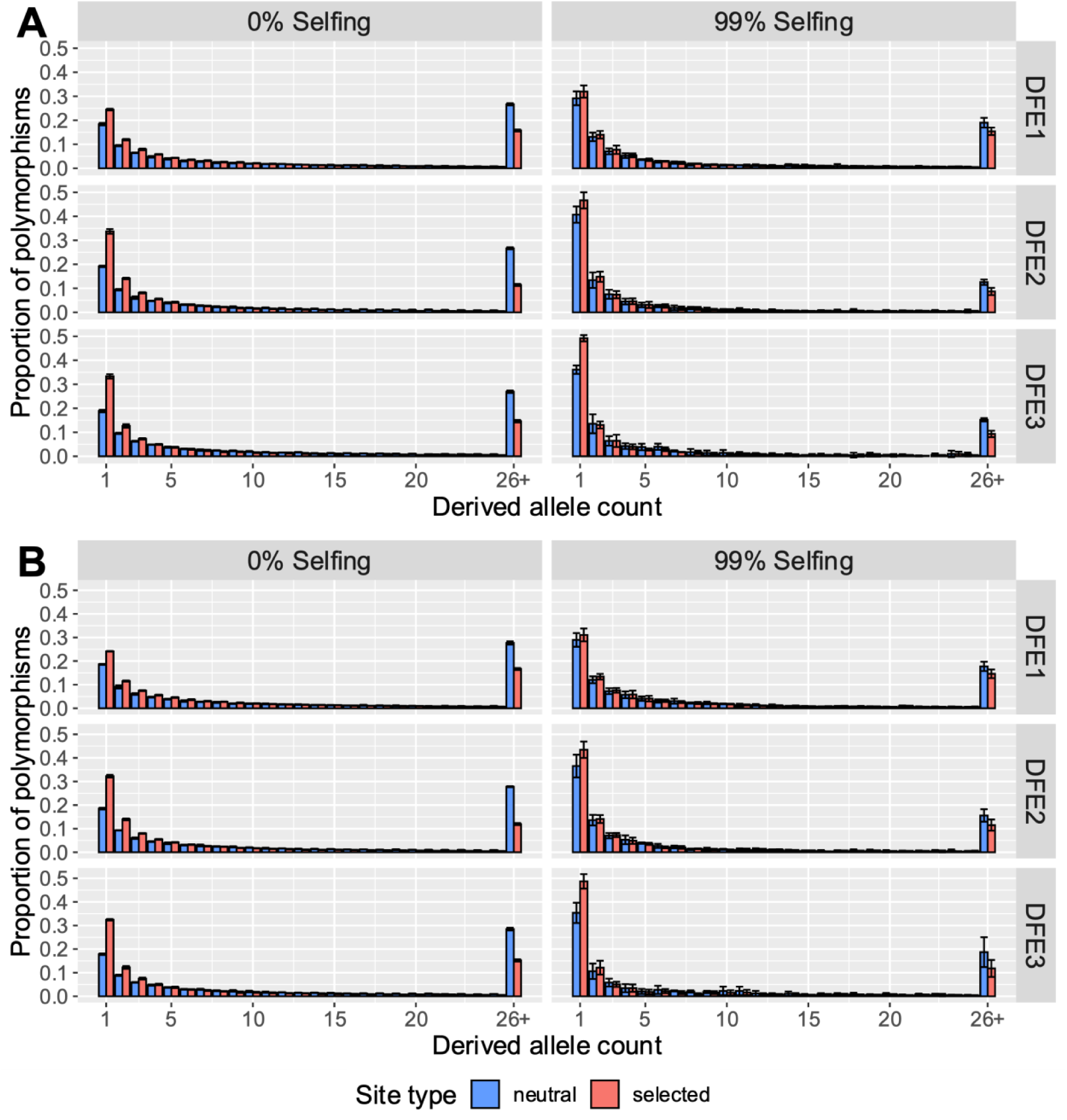
Effects of uneven sampling, population structure, and selfing on the SFS at high migration rates. Populations were simulated under an island model with five demes, and genomes were sampled unevenly (35 genomes from two demes, and 10 from the remaining four demes). Here the metapopulation effective size was equal to 5000 at 0% selfing (see methods). Results are shown when *N*_*deme*_ *m* was (**A)** 2 **(B)** 1. The error bars denote the standard deviation of proportions estimated from 5 independent replicates. The last class (26+) refers to the derived allele counts 26-100.

**Supplementary Figure 31:**
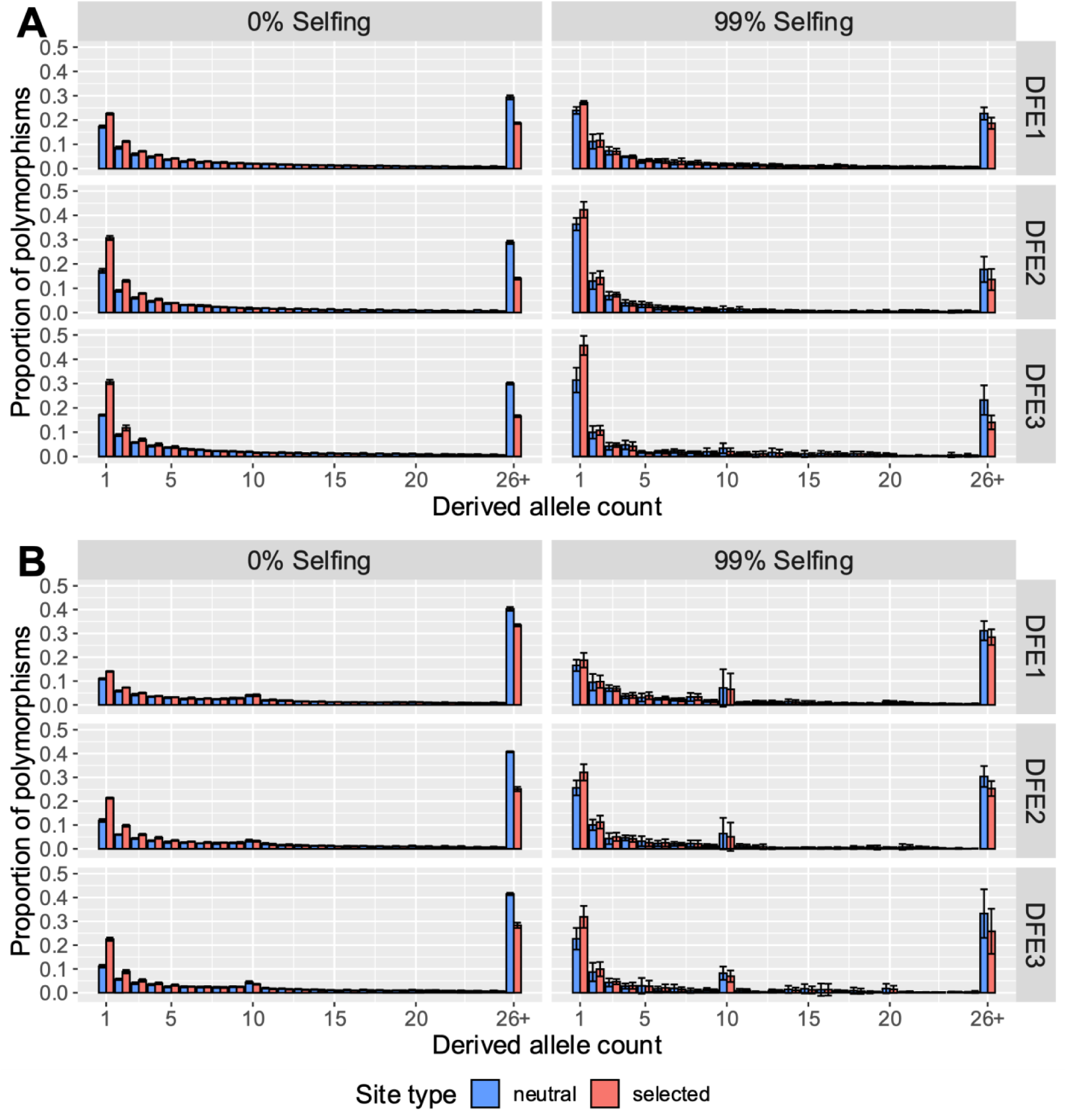
Effects of uneven sampling, population structure, and selfing on the SFS when rates of migration are low. Populations were simulated under an island model with five demes, and genomes were sampled unevenly (35 genomes from two demes, and 10 from the remaining four demes). Here the metapopulation effective size was equal to 5000 at 0% selfing (see methods). Results are shown when *N*_*deme*_ *m* was **(A)** 0.5 **(B)** 0.1. The error bars denote the standard deviation of proportions estimated from 5 independent replicates. The last class (26+) refers to the derived allele counts 26-100.

**Supplementary Figure 32:**
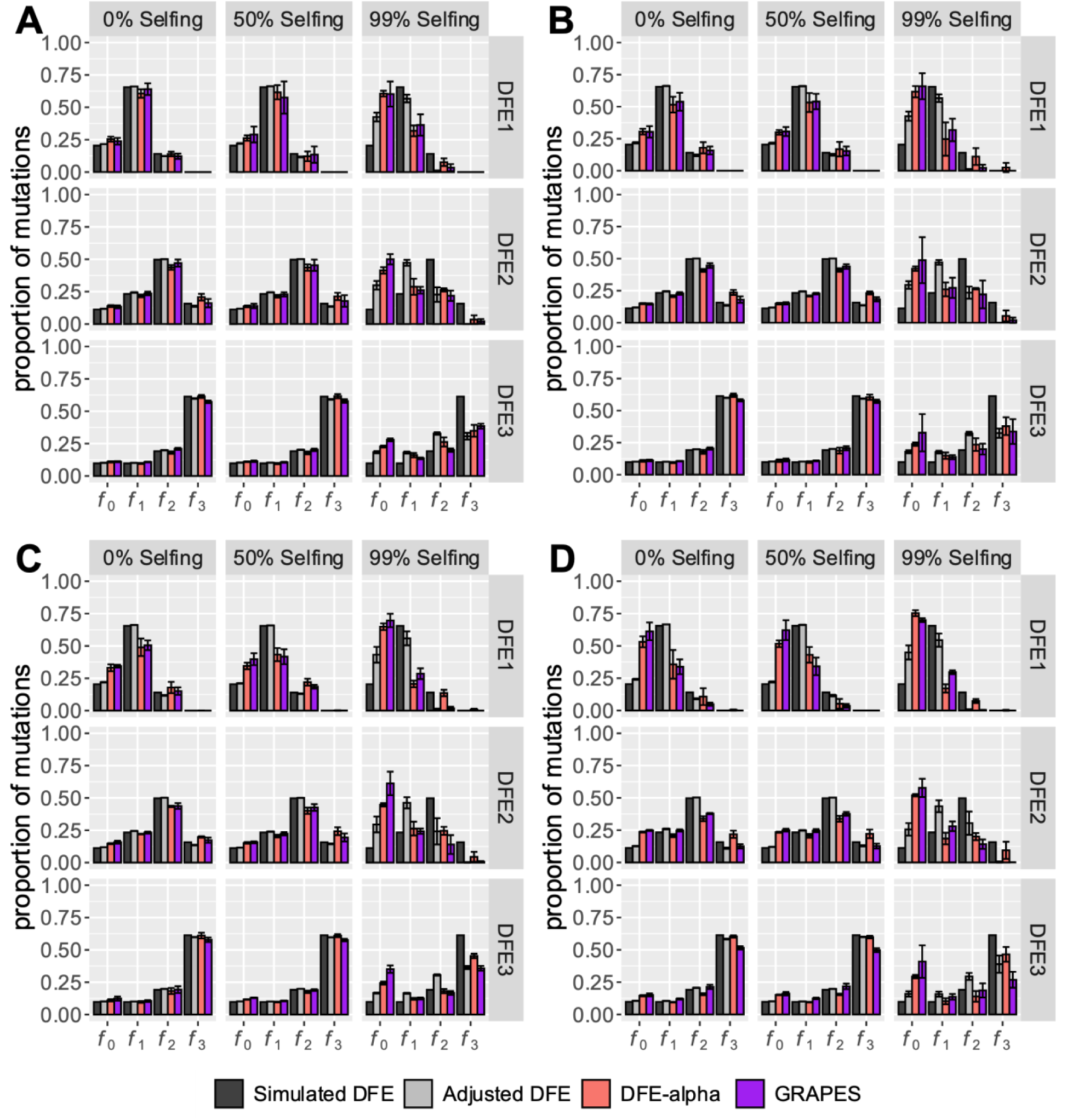
Effects of uneven sampling, population structure, and selfing on the inference of the DFE of new deleterious mutations using DFE-alpha and GRAPES. Populations were simulated under an island model with five demes, and genomes were sampled unevenly (60 genomes from one deme, and 10 from the remaining four demes). Here the metapopulation

**Supplementary Figure 33:**
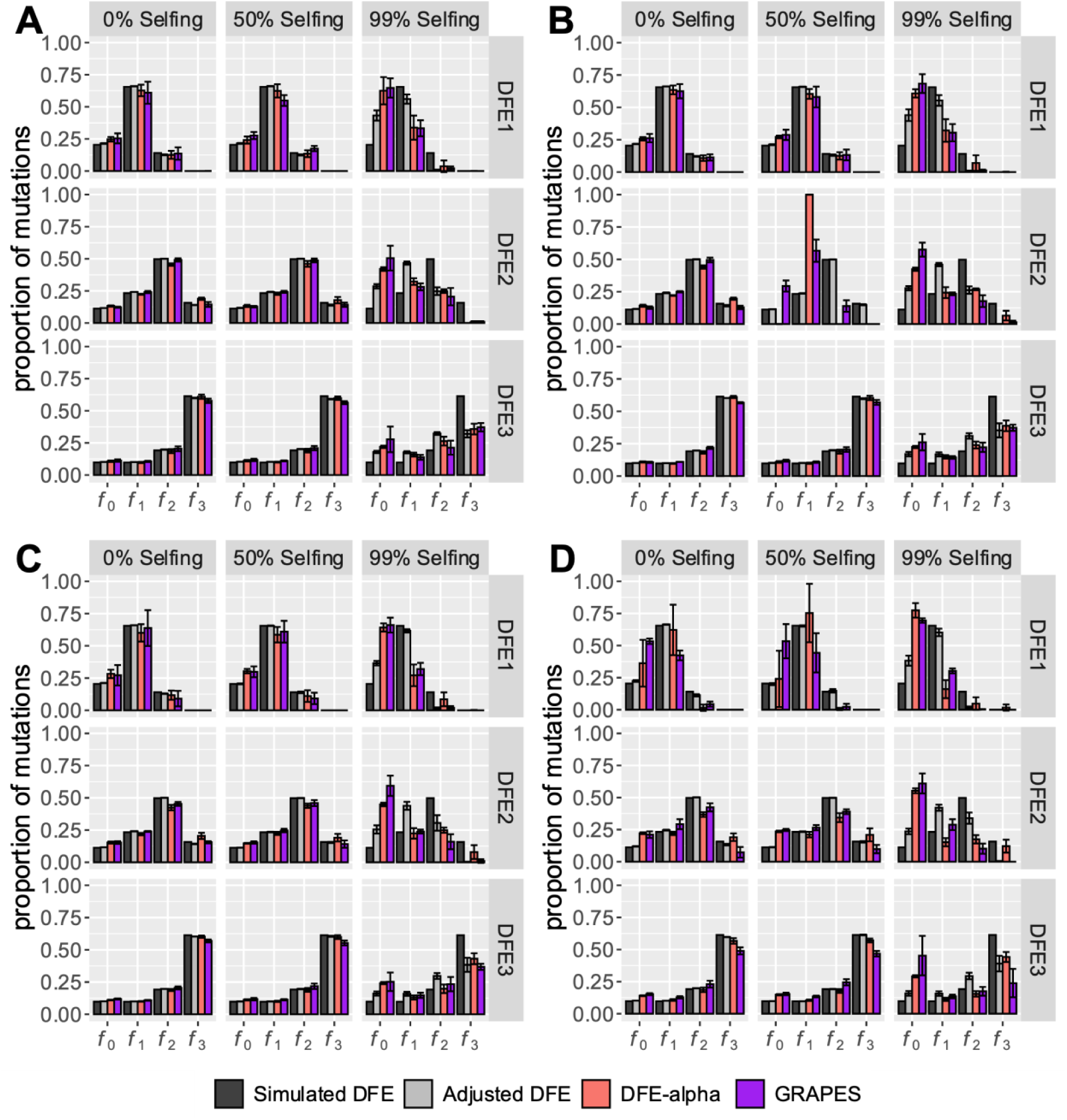
Effects of uneven sampling, population structure and selfing on the inference of the DFE of new deleterious mutations using DFE-alpha and GRAPES. Populations were simulated under an island model with five demes, and genomes were sampled unevenly (35 genomes from two demes, and 10 from the remaining three demes). Here the metapopulation

**Supplementary Table 1.** Effects of selfing on the mean values of parameters of the DFE of new deleterious mutations, estimated by DFE-alpha. Here *E*[*s*] represents the mean efficacy of selection, before being scaled by a 2-epoch model of population size change, where *N*1 represents the ancestral population size, *N*2 represents the relative population size after the first epoch, and *t*2 represents the duration of time after the population size change (in generations).

| | Selfing<br>rate | $E[s]$ | $N1$ | $N2$ | $t2$ |
| --- | --- | --- | --- | --- | --- |
| DFE1 | Truth | -0.025 | 100 | NA | NA |
|  | 0% | -0.027 | 100 | 145 | 16 |
|  | 50% | -0.025 | 100 | 153 | 7 |
|  | 80% | -0.021 | 100 | 162 | 5 |
|  | 90% | -0.023 | 100 | 150 | 17 |
|  | 95% | -0.019 | 100 | 179 | 29 |
|  | 99% | -0.013 | 100 | 394 | 77 |
| DFE2 | Truth | -0.25 | 100 | 100 | NA |
|  | 0% | -0.269 | 100 | 158 | 5 |
|  | 50% | -0.263 | 100 | 150 | 6 |
|  | 80% | -0.223 | 100 | 168 | 5 |
|  | 90% | -0.187 | 100 | 238 | 5 |
|  | 95% | -0.099 | 100 | 322 | 22 |
|  | 99% | -0.026 | 100 | 1000 | 57 |
| DFE3 | Truth | -5 | 100 | 100 | NA |
|  | 0% | -4.712 | 100 | 144 | 5 |
|  | 50% | -5.465 | 100 | 156 | 5 |
|  | 80% | -4.62 | 100 | 162 | 5 |
|  | 90% | -2.665 | 100 | 198 | 11 |
|  | 95% | -1.722 | 100 | 319 | 17 |
|  | 99% | -0.687 | 100 | 798 | 59 |

**Supplementary Table 2.** Estimated parameter values of the deleterious DFE by GRAPES and DFE-alpha for simulations with only semi-dominant deleterious mutations.

| | | $\beta$ | | | | $2N_e s_d$ | | | |
| --- | --- | --- | --- | --- | --- | --- | --- | --- | --- |
|  |  | GRAPES |  | DFE-alpha |  | GRAPES |  | DFE-alpha |  |
|  | Selfing rate | mean | SD | mean | SD | mean | SD | mean | SD |
| DFE1 | Truth | 0.9 | 0 | 0.9 | 0 | 5 | 0 | 5 | 0 |
|  | 0% | 0.77 | 0.16 | 0.74 | 0.04 | 5.43 | 0.58 | 5.4 | 0.33 |
|  | 50% | 0.85 | 0.09 | 0.82 | 0.16 | 4.97 | 0.15 | 5.12 | 0.5 |
|  | 80% | 0.93 | 0.30 | 1 | 0.29 | 4.5 | 0.81 | 4.27 | 0.59 |
|  | 90% | 0.65 | 0.11 | 0.78 | 0.16 | 5.25 | 1.07 | 4.8 | 0.92 |
|  | 95% | 0.57 | 0.15 | 0.68 | 0.14 | 4.42 | 0.39 | 4 | 0.48 |
|  | 99% | 0.34 | 0.03 | 0.26 | 0.03 | 2.34 | 0.59 | 3.2 | 0.9 |
| DFE2 | Truth | 0.5 | 0 | 0.5 | 0 | 50 | 0 | 50 | 0 |
|  | 0% | 0.50 | 0.01 | 0.48 | 0.01 | 50.6 | 4.12 | 54.23 | 5.35 |
|  | 50% | 0.48 | 0.04 | 0.47 | 0.02 | 52 | 6.64 | 53.12 | 3.45 |
|  | 80% | 0.49 | 0.06 | 0.48 | 0.03 | 44.4 | 7.33 | 45.14 | 3.73 |
|  | 90% | 0.45 | 0.05 | 0.47 | 0.03 | 41.87 | 10.37 | 37.87 | 6.4 |
|  | 95% | 0.39 | 0.07 | 0.46 | 0.07 | 29.44 | 9.1 | 21.12 | 5.89 |
|  | 99% | 0.33 | 0.15 | 0.35 | 0.04 | 8.93 | 1.38 | 6.39 | 1.41 |
| DFE3 | Truth | 0.3 | 0 | 0.3 | 0 | 1000 | 0 | 1000 | 0 |
|  | 0% | 0.32 | 0.02 | 0.3 | 0.01 | 627.15 | 107.89 | 949.56 | 138.64 |
|  | 50% | 0.30 | 0.02 | 0.29 | 0.02 | 730.72 | 133.39 | 1102.76 | 243.83 |
|  | 80% | 0.30 | 0.05 | 0.29 | 0.03 | 743.11 | 415.73 | 933.55 | 448.17 |
|  | 90% | 0.28 | 0.04 | 0.29 | 0.01 | 663.41 | 468.89 | 546.89 | 156.22 |
|  | 95% | 0.26 | 0.04 | 0.28 | 0.01 | 512.23 | 233.45 | 363.49 | 62.58 |
|  | 99% | 0.17 | 0.06 | 0.26 | 0.03 | 619.02 | 344.14 | 171.07 | 99.41 |

**Supplementary Table 3.** The number of segregating sites and reduction in population size due to BGS and selfing for varying selfing rates and DFEs as observed in our simulations. Here *N*=5000 and *B* = *π*_*neu*_ ⁄(4*N*_*self*_ *μ*), where *μ* is the mutation rate per site/generation.

| | Selfing rate | Number of deleterious segregating sites | | $\pi_{neu}$ | $N_{emp}$ | $N_{self}$ | $B$ |
| --- | --- | --- | --- | --- | --- | --- | --- |
|  |  | mean | SD |  |  |  |  |
| DFE1 | 0% | 13564 | 137 | 0.00626 | 4746 | 5000 | 0.949 |
|  | 50% | 10077 | 121 | 0.00464 | 3512 | 3750 | 0.936 |
|  | 80% | 7950 | 112 | 0.00352 | 2664 | 3000 | 0.888 |
|  | 90% | 6997 | 131 | 0.00299 | 2264 | 2750 | 0.823 |
|  | 95% | 6260 | 190 | 0.00244 | 1850 | 2625 | 0.705 |
|  | 99% | 4281 | 293 | 0.0011 | 835 | 2525 | 0.331 |
| DFE2 | 0% | 7930 | 103 | 0.00614 | 4654 | 5000 | 0.931 |
|  | 50% | 5837 | 105 | 0.00448 | 3392 | 3750 | 0.905 |
|  | 80% | 4463 | 95 | 0.00322 | 2440 | 3000 | 0.813 |
|  | 90% | 3559 | 137 | 0.00225 | 1706 | 2750 | 0.620 |
|  | 95% | 2656 | 244 | 0.00114 | 865 | 2625 | 0.329 |
|  | 99% | 1628 | 135 | 0.00028 | 212 | 2525 | 0.084 |
| DFE3 | 0% | 4273 | 49 | 0.0059 | 4470 | 5000 | 0.894 |
|  | 50% | 3003 | 36 | 0.00406 | 3078 | 3750 | 0.821 |
|  | 80% | 2048 | 68 | 0.00257 | 1946 | 3000 | 0.649 |
|  | 90% | 1602 | 25 | 0.00165 | 1254 | 2750 | 0.456 |
|  | 95% | 1213 | 54 | 0.00097 | 737 | 2625 | 0.281 |
|  | 99% | 818 | 69 | 0.00032 | 241 | 2525 | 0.096 |

**Supplementary Table 4.**
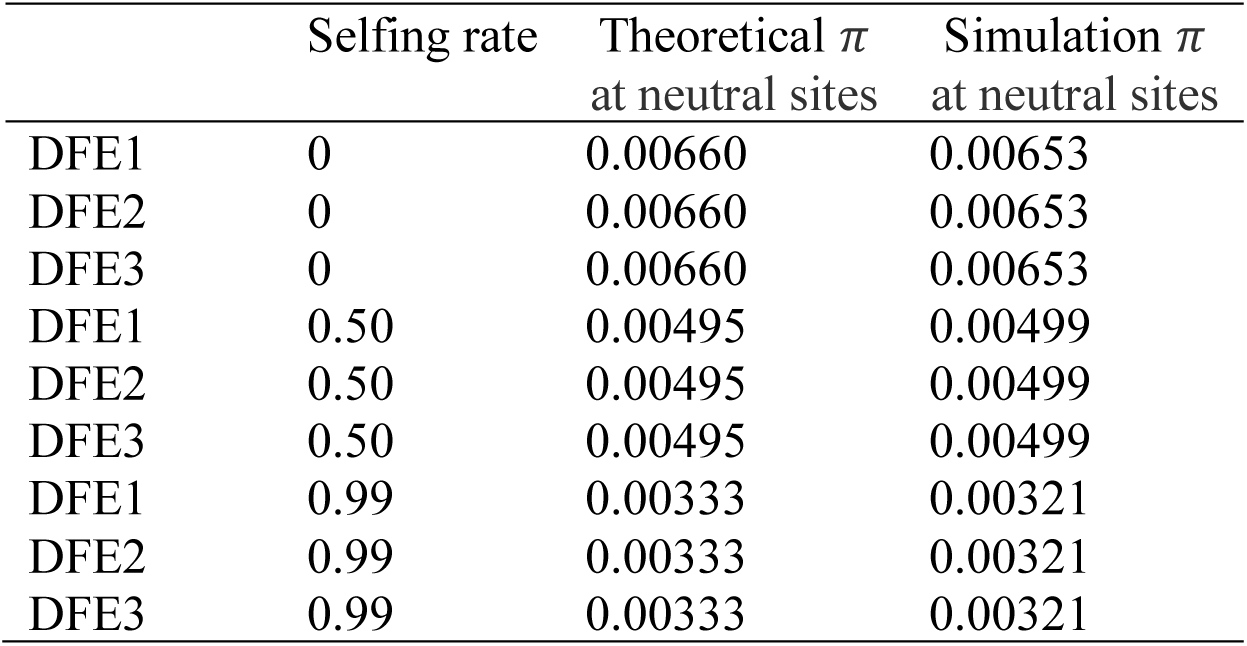
Nucleotide site diversity (*π*) from single-site simulations of neutral and semidominant deleterious mutations in populations with varying selfing rates.

| | Selfing rate | Theoretical $\pi$<br>at neutral sites | Simulation $\pi$<br>at neutral sites |
| --- | --- | --- | --- |
| DFE1 | 0 | 0.00660 | 0.00653 |
| DFE2 | 0 | 0.00660 | 0.00653 |
| DFE3 | 0 | 0.00660 | 0.00653 |
| DFE1 | 0.50 | 0.00495 | 0.00499 |
| DFE2 | 0.50 | 0.00495 | 0.00499 |
| DFE3 | 0.50 | 0.00495 | 0.00499 |
| DFE1 | 0.99 | 0.00333 | 0.00321 |
| DFE2 | 0.99 | 0.00333 | 0.00321 |
| DFE3 | 0.99 | 0.00333 | 0.00321 |

**Supplementary Table 5.** Estimates of the parameters of the deleterious DFE (a gamma distribution with mean 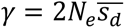 and shape parameter *β*) inferred by DFE-alpha and GRAPES. Since DFE-alpha and GRAPES assume semidominance, the mean strength of selection *γ* was multiplied by 0.5 to be equivalent to 2*N*_*self*_ *s*_*d*_*h*_*self*_.

| DFE | Selfing rate | $2N_{self}s_dh_{self}$<br>(expected) | DFE-alpha<br>$\gamma h$ | GRAPES<br>$\gamma h$ | $\beta$<br>(expected) | DFE-alpha<br>$\beta$ | GRAPES<br>$\beta$ | $h$ |
| --- | --- | --- | --- | --- | --- | --- | --- | --- |
| DFE1 | 0 | 3.75 | 4.29 | 4.33 | 0.9 | 0.62 | 0.64 | 0.75 |
| DFE1 | 0.50 | 4.04 | 3.32 | 3.49 | 0.9 | 0.68 | 0.66 | 0.75 |
| DFE1 | 0.99 | 4.17 | 1.40 | 1.01 | 0.9 | 0.42 | 0.34 | 0.75 |
| DFE2 | 0 | 37.5 | 41.45 | 34.99 | 0.5 | 0.44 | 0.48 | 0.75 |
| DFE2 | 0.50 | 40.38 | 30.68 | 27.77 | 0.5 | 0.46 | 0.49 | 0.75 |
| DFE2 | 0.99 | 41.67 | 4.23 | 4.54 | 0.5 | 0.30 | 0.19 | 0.75 |
| DFE3 | 0 | 750.00 | 791.63 | 493.84 | 0.3 | 0.29 | 0.31 | 0.75 |
| DFE3 | 0.50 | 807.69 | 630.82 | 429.23 | 0.3 | 0.28 | 0.30 | 0.75 |
| DFE3 | 0.99 | 833.32 | 50.93 | 222.49 | 0.3 | 0.29 | 0.18 | 0.75 |
| DFE1 | 0 | 2.50 | 2.70 | 2.71 | 0.9 | 0.74 | 0.77 | 0.50 |
| DFE1 | 0.50 | 3.08 | 2.56 | 2.48 | 0.9 | 0.82 | 0.85 | 0.50 |
| DFE1 | 0.99 | 3.33 | 1.60 | 1.17 | 0.9 | 0.26 | 0.34 | 0.50 |
| DFE2 | 0 | 25.00 | 27.11 | 25.30 | 0.5 | 0.48 | 0.49 | 0.50 |
| DFE2 | 0.50 | 30.77 | 26.56 | 26.00 | 0.5 | 0.47 | 0.48 | 0.50 |
| DFE2 | 0.99 | 33.33 | 3.19 | 4.46 | 0.5 | 0.35 | 0.33 | 0.50 |
| DFE3 | 0 | 500.00 | 474.78 | 313.57 | 0.3 | 0.30 | 0.32 | 0.50 |
| DFE3 | 0.50 | 615.38 | 551.38 | 365.36 | 0.3 | 0.29 | 0.30 | 0.50 |
| DFE3 | 0.99 | 666.64 | 85.53 | 309.51 | 0.3 | 0.26 | 0.17 | 0.50 |
| DFE1 | 0 | 1.25 | 1.28 | 1.33 | 0.9 | 2.66 | 2.07 | 0.25 |
| DFE1 | 0.50 | 2.12 | 1.66 | 1.74 | 0.9 | 1.54 | 1.49 | 0.25 |
| DFE1 | 0.99 | 2.50 | 1.56 | 0.98 | 0.9 | 0.32 | 0.39 | 0.25 |
| DFE2 | 0 | 12.50 | 10.73 | 10.62 | 0.5 | 0.63 | 0.64 | 0.25 |
| DFE2 | 0.50 | 21.15 | 18.06 | 15.95 | 0.5 | 0.51 | 0.56 | 0.25 |
| DFE2 | 0.99 | 25.00 | 4.50 | 4.65 | 0.5 | 0.29 | 0.20 | 0.25 |
| DFE3 | 0 | 250.00 | 263.78 | 217.20 | 0.3 | 0.31 | 0.31 | 0.25 |
| DFE3 | 0.50 | 423.08 | 316.34 | 253.83 | 0.3 | 0.31 | 0.32 | 0.25 |
| DFE3 | 0.99 | 499.97 | 54.93 | 183.24 | 0.3 | 0.28 | 0.25 | 0.25 |
| DFE1 | 0 | 0.50 | 0.91 | 0.88 | 0.9 | 100 | 16.45 | 0.10 |
| DFE1 | 0.50 | 1.54 | 1.24 | 1.31 | 0.9 | 3.44 | 9.15 | 0.10 |
| DFE1 | 0.99 | 2.00 | 1.32 | 0.84 | 0.9 | 0.40 | 0.36 | 0.10 |
| DFE2 | 0 | 5.00 | 4.54 | 4.92 | 0.5 | 1.05 | 0.94 | 0.10 |
| DFE2 | 0.50 | 15.38 | 12.19 | 12.42 | 0.5 | 0.58 | 0.56 | 0.10 |
| DFE2 | 0.99 | 20.00 | 5.02 | 6.41 | 0.5 | 0.32 | 0.22 | 0.10 |
| DFE3 | 0 | 100.00 | 69.53 | 63.27 | 0.3 | 0.38 | 0.39 | 0.10 |
| DFE3 | 0.50 | 307.69 | 192.87 | 159.62 | 0.3 | 0.33 | 0.33 | 0.10 |
| DFE3 | 0.99 | 399.96 | 72.07 | 213.35 | 0.3 | 0.26 | 0.19 | 0.10 |

**Supplementary Table 6.** Effects of selfing on the number of fixations of deleterious and beneficial mutations. Here, deleterious mutations were drawn from the DFEs defined in Table 1, while 0.1% of new exonic mutations were beneficial and exponentially distributed with a mean 2*Ns*_*a*_ = 200.

| Selfing rate | DFE | Deleterious substitutions avg. | Deleterious substitutions SD | Beneficial substitutions avg. | Beneficial substitutions SD | $\alpha$ avg. | $\alpha$ SD |
| --- | --- | --- | --- | --- | --- | --- | --- |
| 0 | DFE1 | 1438.4 | 31.48 | 935.8 | 32.09 | 0.39 | 0.01 |
| 50 | DFE1 | 1514.2 | 56.92 | 912.2 | 30.19 | 0.38 | 0.01 |
| 80 | DFE1 | 1820.8 | 28.03 | 813.4 | 17.9 | 0.31 | 0.01 |
| 90 | DFE1 | 2253.6 | 68.2 | 681.8 | 14.75 | 0.23 | 0.01 |
| 95 | DFE1 | 2785 | 77.11 | 552.4 | 19.55 | 0.17 | 0 |
| 99 | DFE1 | 3363.8 | 56.91 | 343.4 | 15.53 | 0.09 | 0 |
| 0 | DFE2 | 618.2 | 7.82 | 912 | 40.21 | 0.6 | 0.01 |
| 50 | DFE2 | 682 | 28.57 | 878.2 | 33.25 | 0.56 | 0.01 |
| 80 | DFE2 | 810 | 46.98 | 753.4 | 33.07 | 0.48 | 0.02 |
| 90 | DFE2 | 1087.4 | 31.5 | 617.6 | 20.02 | 0.36 | 0.01 |
| 95 | DFE2 | 1451.2 | 28.01 | 464 | 5.7 | 0.24 | 0 |
| 99 | DFE2 | 2133 | 26.54 | 276.6 | 7.83 | 0.11 | 0 |
| 0 | DFE3 | 460.8 | 11.76 | 869.2 | 24.38 | 0.65 | 0.01 |
| 50 | DFE3 | 509 | 16.39 | 806.6 | 25.75 | 0.61 | 0.01 |
| 80 | DFE3 | 543.2 | 27.22 | 620.6 | 18.99 | 0.53 | 0.01 |
| 90 | DFE3 | 635.6 | 34.25 | 464.8 | 25.96 | 0.42 | 0.01 |
| 95 | DFE3 | 762.2 | 24.94 | 341.4 | 20.95 | 0.31 | 0.02 |
| 99 | DFE3 | 1040.8 | 36.62 | 191.6 | 18.77 | 0.16 | 0.01 |

**Supplementary Table 7.** Parameters used for five-deme island model simulations.

| $N_{\text{deme}}m$ | $m$ | $N_{\text{deme}}$ |
| --- | --- | --- |
| 10 | 0.0102 | 984 |
| 2 | 0.00216 | 926 |
| 1 | 0.00116 | 862 |
| 0.5 | 0.00066 | 785 |
| 0.1 | 0.00026 | 385 |

**Supplementary Table 8.** Table showing how theoretical expectations for *T*_*T*_, *T_S_*, and *F*_*st*_ match values estimated from neutral simulations.

| $N_{deme}m$ | Selfing rate | $\pi_{meta}$ expected | $\pi_{meta}$ observed | $\pi_{deme}$ expected | $\pi_{deme}$ observed | $F_{st}$ expected | $F_{st}$ observed |
| --- | --- | --- | --- | --- | --- | --- | --- |
| 0.1 | 0 | 0.00660 | 0.00659 | 0.00254 | 0.00259 | 0.61515 | 0.60713 |
| 0.1 | 0.50 | 0.00597 | 0.00599 | 0.00191 | 0.00211 | 0.68063 | 0.64825 |
| 0.1 | 0.99 | 0.00534 | 0.00567 | 0.00128 | 0.00140 | 0.75991 | 0.76400 |
| 0.5 | 0 | 0.00678 | 0.00661 | 0.00518 | 0.00507 | 0.23595 | 0.23265 |
| 0.5 | 0.50 | 0.00549 | 0.00537 | 0.00389 | 0.00375 | 0.29166 | 0.30229 |
| 0.5 | 0.99 | 0.00422 | 0.00425 | 0.00262 | 0.00291 | 0.37947 | 0.31402 |
| 1.0 | 0 | 0.00660 | 0.00659 | 0.00569 | 0.00564 | 0.13794 | 0.14426 |
| 1.0 | 0.50 | 0.00518 | 0.00519 | 0.00427 | 0.00434 | 0.17584 | 0.16317 |
| 1.0 | 0.99 | 0.00378 | 0.00402 | 0.00287 | 0.00305 | 0.24062 | 0.24016 |
| 2.0 | 0 | 0.00660 | 0.00657 | 0.00611 | 0.00611 | 0.07407 | 0.06951 |
| 2.0 | 0.50 | 0.00507 | 0.00508 | 0.00458 | 0.00457 | 0.09638 | 0.09950 |
| 2.0 | 0.99 | 0.00358 | 0.00345 | 0.00309 | 0.00295 | 0.13674 | 0.14377 |

**Supplementary Table 9.** Estimated levels of background selection (*B*) in highly selfing species for the range of previously reported selfing rates. All species are diploid except *C. bursa-pastoris*. The number of deleterious mutations per genome per generation (*U*_*d*_) was estimated by multiplying the number of coding sites by 0.7. The strength of BGS was calculated with 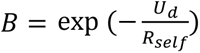, where *R*_self_ = *R* × *F*. This supplementary table contains all mutation and recombination rate data used for the calculations in Table 2 of the main text. (table stored in a separate attachment)

